# The Myc-Like Mlx Network Impacts Aging and Metabolism

**DOI:** 10.1101/2023.11.26.568749

**Authors:** Huabo Wang, Taylor Stevens, Jie Lu, Alexander Roberts, Clinton Van’t Land, Radhika Muzumdar, Zhenwei Gong, Jerry Vockley, Edward V. Prochownik

## Abstract

The “Mlx” and “Myc” Networks share many common gene targets. Just as Myc’s activity depends upon its heterodimerization with Max, the Mlx Network requires that the Max-like factor Mlx associate with the Myc-like factors MondoA or ChREBP. We show here that body-wide *Mlx* inactivation, like that of *Myc,* accelerates numerous aging-related phenotypes pertaining to body habitus and metabolism. The deregulation of numerous aging-related Myc target gene sets is also accelerated. Among other functions, these gene sets often regulate ribosomal and mitochondrial structure and function, genomic stability and aging. Whereas “*Myc*KO” mice have an extended lifespan because of a lower cancer incidence, “*Mlx*KO” mice have normal lifespans and a somewhat higher cancer incidence. Like Myc, Mlx, MondoA and ChREBP expression and that of their target genes, deteriorate with age in both mice and humans, underscoring the importance of life-long and balanced cross-talk between the two Networks to maintain normal aging.

**Teaser:** Inactivation of the Myc-like “Mlx Network” in mice leads to phenotypic and molecular signs of premature aging and a cancer predisposition.

## Introduction

The bHLH-ZIP transcription factor (TF) c-Myc (Myc) is de-regulated and/or over-expressed in most human cancers, is highly transforming in various *in vivo* contexts and can be linked functionally to most canonical “Cancer Hallmarks” (*1–3*). Upon heterodimerizing with its bHLH-ZIP partner Max, Myc binds one or more “E box” sites (CACGTG) that usually reside in the proximal promoters of its many direct target genes. It then recruits an assortment of companion co-factors such as histone acetylases and H3K4 methylases thereby facilitating transcriptional initiation and read-through and reducing pausing by RNA pol II (*4–6*). These activities are countered by four tissue-restricted bHLH-ZIP “Mxd” members (Mxd1-4) or the more distantly related factors Mnt and Mga. These also heterodimerize with Max and compete with one another and with Myc-Max for E-box occupancy and negate Myc target gene activation by recruiting histone deacetylases (*7, 8*). Myc-Max-driven interactions are favored during rapid proliferation when Myc is highly expressed whereas Mxd/Mnt/Mga-Max interactions predominate during periods of quiescence or in differentiated cells when Myc levels are low or declining (*7, 8*). Negative transcriptional regulation by Myc, which accounts for nearly half of its effects on gene expression, is indirect, E-box-independent and mediated by inhibitory interactions between Myc-Max and TFs such as Miz1 and Sp1/3, which bind at their own distinct promoter-proximal Miz1 and SP1 sites, respectively (*9, 10*).

Fundamental processes supervised by the above “Myc Network” include ribosomal biogenesis and translation, cell cycle progression and energy production (*7, 11–14*). The latter encompasses glycolysis, glutaminolysis, fatty acid oxidation (FAO), oxidative phosphorylation (Oxphos) and the maintenance of mitochondrial structure (*1, 13–18*). Collectively, these processes support biomass accumulation and the maintenance of a reduced intracellular environment, all of which decline with age as does the expression of Myc itself and many of its direct target genes (*11, 19–24*).

A second group of bHLH-ZIP TFs with structural and functional relatedness to Myc Network members are those which comprise the “Mlx Network” (*13, 14, 25, 26*). They regulate a smaller direct target gene repertoire than do Myc Network TFs although considerable overlap exists (*13, 14, 24–29*). The positively-acting, Myc-like members of this Network, ChREBP and MondoA, interact with the Max-like protein Mlx to form transcriptionally active heterodimers. Unlike Myc, however, their nuclear import and transcriptional activity are dependent upon their binding of glucose-6-phosphate, thus making them nutrient-responsive and subservient to metabolic cues (*26*). While these heterodimers recognize E boxes (*13, 14, 25*), they classically bind to more complex “carbohydrate response elements” (ChoREs) consisting of tandem E box-like elements separated by 5 nucleotides (*13, 14, 30*). This binding site flexibility is reinforced by the fact that Mxd1, Mxd4 and Mnt can also heterodimerize with Mlx, displace ChREBP/MondoA-Mlx heterodimers and counter their positive effects on transcription (*7, 25, 31, 32*). In addition to being fewer in number, Mlx Network direct target genes were initially believed to be more functionally restricted than those of the Myc Network and were primarily viewed as participating in lipid and carbohydrate metabolism (*7, 25, 31–35*). More recently, however, extended roles for the Mlx Network in maintaining ribosomal and mitochondrial structure and function, both alone and in collaboration with Myc, have been reported(*13, 14, 24, 28, 29*).

The mid-gestational embryonic lethality associated with *Myc* gene inactivation has until recently stymied a complete understanding of its role(s) in normal physiology, particularly over the course of a lifetime (*36*)*. In vivo* studies therefore have been restricted to examining the consequences of such inactivation in individual tissues or in *Myc+/-* mice, both of which are compatible with long-term survival (*15, 37–39*)*. Myc+/-* animals are smaller than their WT counterparts, age less rapidly, live significantly longer and are generally healthier. These studies showed that Myc impacts aging although precisely how and the degree to which the smaller size of the animals might have contributed to this was not addressed (*38, 40*). They also suggested that a more complete loss of Myc would reveal additional phenotypes.

We recently reported the long-term consequences of near-complete body-wide *Myc* gene inactivation in mice initiated at the time of weaning (*23, 40*). The survival of these “*Myc*KO” mice permitted the long-term consequences of the gene’s loss to be studied and its impact on aging to be determined. Unlike *Myc+/-* mice, *Myc*KO mice were of normal size or even somewhat larger than their wild-type (WT), *Myc+/+* counterparts. However, they gradually developed many features of premature aging that coincided with the dysregulation of numerous gene sets involved in ribosomal and mitochondrial structure and function, oxidative stress, redox balance, mRNA splicing and DNA damage response and repair. They also prematurely altered the expression of multiple sets of genes typically associated with normal or premature aging and senescence. Yet despite their pronounced premature aging profiles, *Myc*KO mice lived significantly longer than WT mice, most likely as a result of their having a 3-4-fold lower lifetime incidence of cancer, the generation and maintenance of which is highly Myc-dependent (*3*). The infrequent tumors that did arise tended to originate from minority cell populations that had escaped complete *Myc* excision. Remarkably, normal mouse and human tissues commonly showed significant age-related declines in Myc expression and an even more extensive loss of properly maintained Myc target gene expression. Together, these findings showed that the long-known intimate association between aging and cancer could be broken and was maintained to a large degree by a single gene, namely *Myc* (*23, 41*).

Given the substantial overlap of the target genes regulated by the Myc and Mlx Networks (*13, 14, 25*), we sought to determine here whether and to what extent the body-wide inactivation of *Mlx* recapitulates any of the life-long consequences previously described in *Myc+/-* or *Myc*KO mice (*23, 38, 40*). Because the loss of Myc can either slow or accelerate aging in a manner than reflects gene dosage (*23, 24, 40*), we paid particular attention to age-related phenotypes. In doing so, we chose to *inactivate Mlx* in the same manner as we previously did for *Myc* (i.e. at the time of weaning) so as to ensure that any observed differences were unlikely to be attributable to the timing or extent of target gene knockout or the manner by which this was achieved. This allowed us to demonstrate that *Mlx*KO mice display some of the same whole body, tissue and molecular changes as their age-matched *Myc*KO counterparts while also manifesting distinct differences that are consistent with the interplay between the two Networks.

## Results

### Post-natal excision of the *Mlx* locus is highly efficient and permanent

*Mlx^LoxP/LoxP^* C578Bl6 mice containing either one or 2 copies of the ROSA-CreER transgene were generated as described previously and as summarized in Supplementary Figure 1A-D (*23, 24*). Control mice were the progeny of crosses between ROSA-CreER strain and C57Bl6 mice with unmodified *Mlx* loci that are hereafter referred to as “WT” (wild-type) mice. At the time of weaning and upon attaining weights of at least 15 g, all mice were subjected to 5 daily injections of tamoxifen as previously described (75 mg/Kg/day i.p. in corn oil) (*23*) and the degree of *Mlx* gene excision was evaluated 2 weeks later. On average, excisional efficiencies were higher in mice bearing 2 copies of the CreER transgene and correlated with both Mlx transcript and protein expression (Supplementary Figure 1E&F and Supplementary Table 1). They were therefore used in all subsequent studies and are referred to as “*Mlx*KO” mice. *Mlx* gene excision persisted throughout life and no evidence was obtained to indicate a selective expansion of cells with retained *Mlx* alleles (Supplementary Figure 1E and Supplementary Table 1).

### *Mlx*KO mice show signs of accelerated aging but have normal lifespans

WT and *Mlx*KO mice of both sexes initially showed similar growth rates and maintained indistinguishable weights and fat:lean mass ratios until ∼10 months of age. At this point, females diverged, gaining weight more rapidly and attaining their maximum adult weights ∼4 months earlier than WT mice (∼20 months vs. 24 months). Fat:lean mass ratios also peaked ∼4 months earlier in female *Myc*KO mice and then subsequently declined at similar rates (Figure 1A).

**Figure 1.**
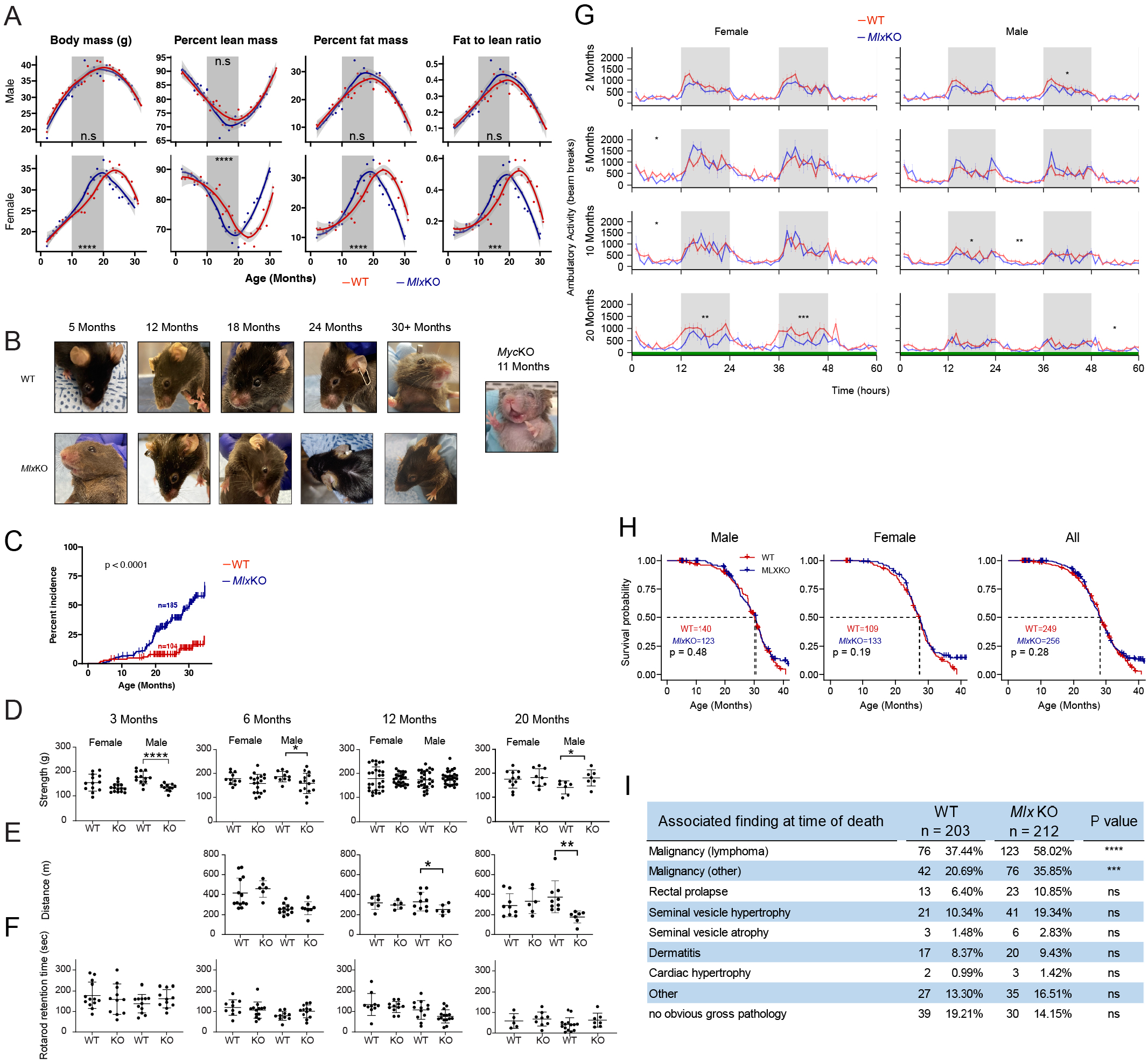
*Mlx*KO mice show attributes of premature aging and a higher cancer incidence but have normal lifespans. **(A)** Weight and body composition of WT and *Mlx*KO mice during the course of life. Measurements were made as previously described beginning at the time of weaning when the *Mlx* gene was first inactivated (*23*). Total body fat and lean mass content were determined by EchoMRI scanning at the time of weighing. Each point represents the mean of measurements from 10-20 animals performed over the course of 2-3 days. Significant differences between the 2 groups are indicated by areas of shading. *=P<0.05, **=P<0.01; ***=P<0.001; ****P<0.0001. **(B)** Overall appearance of *Mlx*KO mice at various ages along with age-matched WT mice. Note typical examples of corneal opacifications (leucoma simplex). The image at the extreme right shows an example of an 11 month old *Myc*KO mouse displaying substantial graying and loss of fur (*23*). **(C)** Cumulative occurrence of both unilateral and bilateral leucoma simplex in WT and *Mlx*KO mice. **(D)** Grip strength of WT and *Mlx*KO mice at the indicated ages. **(E)** Treadmill endurance testing of WT and *Mlx*KO mice at the indicated ages. **(F)** Rotarod testing of WT and *Mlx*KO mice at the indicated ages. In **D-F**, each point represents the mean of 3 tests performed on the same mouse on consecutive days. **(G)** Total diurnal ambulatory activity in 5 month old 18 month old WT and *Mlx*KO mice measured in metabolic cages (*23*). **(H)** Survival of WT and *Myc*KO mice. **(I)** Pathologies associated with WT and *Mlx*KO mice at the time of death.

Premature fur loss (alopecia) and graying (achomotricia) previously observed in *Myc*KO mice, and first noted at ∼5-6 months of age, was not observed in *Mlx*KO mice. However, over two-thirds of the latter animals of both sexes eventually developed unilateral or bilateral corneal opacifications (leucoma simplex) compared to <20% of WT animals (Figure 1B and C). Thus, *Mlx*KO mice developed another external feature of premature aging that was not seen in *Myc*KO mice.

Young male *Mlx*KO mice were significantly weaker than WT mice as measured by grip strength testing (*23*), although this improved with age such that the oldest mice were actually somewhat stronger than their age-matched WT counterparts (Figure 1D). These mice also showed a progressive deterioration in endurance when subjected to treadmill testing (Figure 1E). In contrast, female *Mlx*KO and WT mice could not be distinguished by these tests, irrespective of age. Nor were any differences in balance noted among age-matched cohorts of either sex based on Rotarod testing (Figure 1F). Finally, overall diurnal ambulatory activity, measured during the course of metabolic cage studies (see below), showed no differences until 18-20 months of age when female mice became significantly less active (Figure 1G). Collectively, the results presented in Figure 1A-G show that *Mlx*KO mice prematurely developed several mild features and behaviors associated with aging, although they were somewhat distinct from those seen in *Myc*KO mice (*23*).

The longer survival of *Myc*KO mice has been attributed in part to their >3-fold lower lifetime cancer incidence owing to the virtual absence of this important oncogenic driver (*1, 3, 23, 40*). In contrast *Mlx*KO mice and WT mice of both sexes demonstrated identical lifespans although *Mlx*KO mice had 1.5-fold higher incidences of lymphoma and a 1.7-fold fold higher incidence of other tumors at the time of death (Figure 1H and I). This was in keeping with previous reports that Mlx can function as a suppressor (*23, 31, 40*).

### *Mlx*KO mice accumulate excessive hepatic lipid and malabsorb dietary fat

The incidence of hepatic fat accumulation (steatosis) by both mice and humans increases with age, particularly in association with obesity and insulin resistance (*42, 43*)*. Myc*KO mice also manifest a higher incidence and earlier onset of steatosis and both the genetic and pharmacologic inhibition of Myc in a variety of cell types is associated with excessive neutral lipid accumulation (*15, 23, 24, 28, 29, 44*). Similarly, mice with hepatocyte-specific knockout of *Chrebp* or *Mlx* develop steatosis although precisely when it begins and its natural history have not been investigated (*28, 29*). We confirmed and extended these findings by showing that *Mlx*KO mice as young as 5 months showed more intense hepatic staining with Oil Red O and accumulated ∼4 times more triglyceride (Figure 2A and B). As mice aged, this disparity lessened, although it remained elevated in the latter animals throughout the duration of the study.

**Figure 2.**
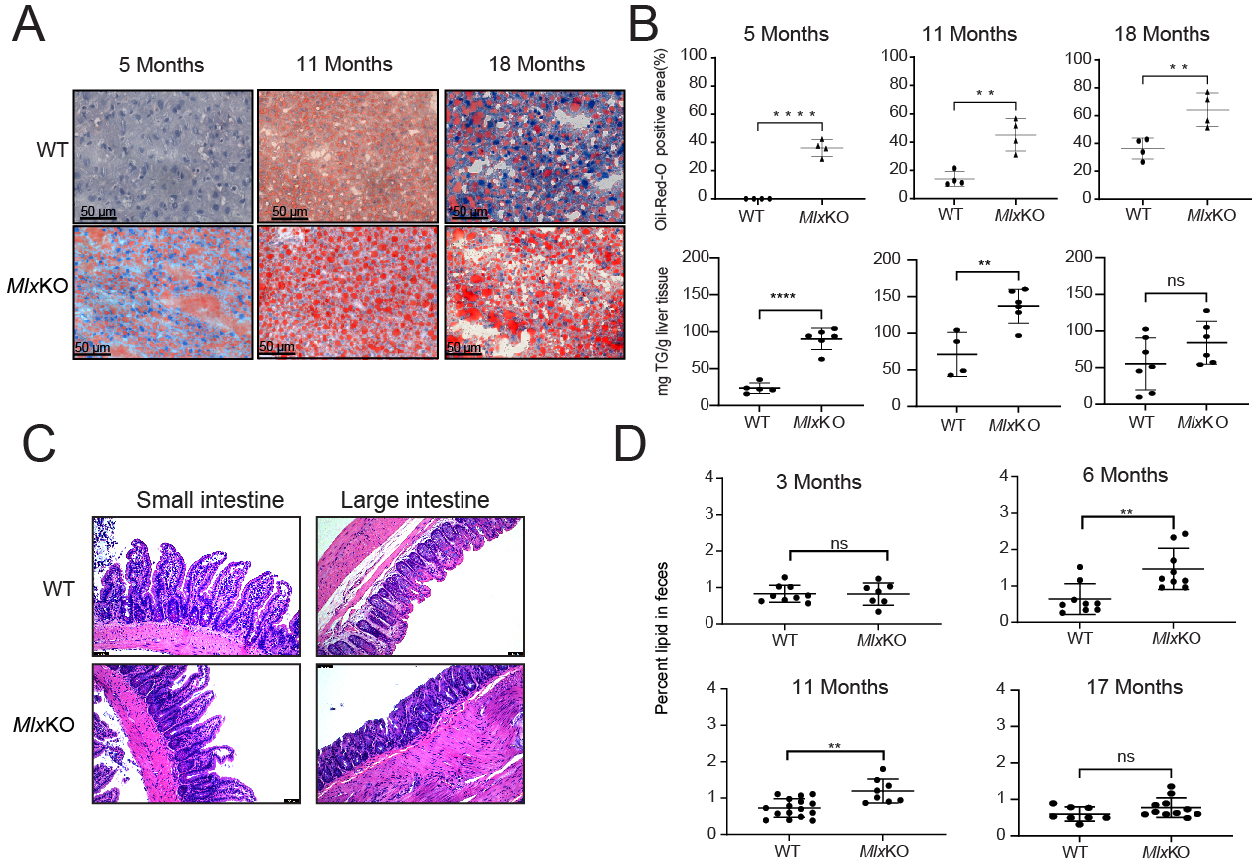
*Mlx*KO prematurely accumulate excessive hepatic lipids and show evidence of dietary fat malabsorption. **(A)** ORO staining of livers from WT and *Mlx*KO mice of the indicated ages **(B)** Triglyceride content in the livers of mice similar to those depicted in A **(C)** H&E-stained sections of small and large intestines of WT and *Mlx*KO mice of the indicated ages **(D)** Fecal fat content from WT and *Mlx*KO mice of the indicated ages. Fecal fat was collected over 48-72 hr from individually caged mice maintained on standard diets.

*Myc*KO mice and those with body-wide expression of the dominant negative inhibitor OmoMyc develop transient flattening of their intestinal mucosa that is not severe enough to cause malabsorption or impair growth (*23, 45*). In contrast, *Mlx*KO mice retained the normal architecture of both their small and large intestines but showed an increased fecal fat content that was noted as early as ∼5-6 months of age and persisted for longer than 11 months (Figure 2C and D). However, as with the above-mentioned reports (*23, 45*) this did not impact growth (Figure 1A).

### *Mlx*KO mice are overly reliant on fatty acid oxidation as an energy source while displaying metabolic inflexibility and abnormal mitochondrial function

Previous metabolic cage studies have shown *Myc*KO mice to be overly reliant on fatty acids as an energy source, particularly during nocturnal feeding when glucose availability is high and normally constitutes the primary energy-generating substrate (*23*). This excessive FAO dependency is also maintained during post-fasting re-feeding with either standard or high-fat diets (HFDs) (*23*). Two month old *Mlx*KO mice showed a similar nocturnal FAO dependency (Figure 3A). Unlike *Myc*KO mice, however, they were also more reliant on glucose during the day when feeding normally declines and fatty acids become alternative energy-generating substrates. The decreased amplitude of the diurnal fluctuation in respiratory exchange ratio (RER = [VCO_2_/VO_2_]) of *Mlx*KO mice pointed to them as being “metabolically inflexible”, a condition whereby excessive nutrient overload and over-competition for metabolic substrates can cause dysregulation in fuel choice and energy generation (*46*). The markedly lower RERs in both WT and *Mlx*KO cohorts in response to fasting indicated that they were similarly adaptable to a period of exogenous glucose deprivation that was rectified by switching their energy source to fatty acids. However, the *Myc*KO cohort’s over-dependence on FAO persisted when either standard or HFDs were re-introduced. The metabolic disparities between *Myc*KO and WT mice changed somewhat over life such that in older adults (ca. 20 months) the excessive reliance on FAO became more generalized across the entire day (Figure 3A).

**Figure 3.**
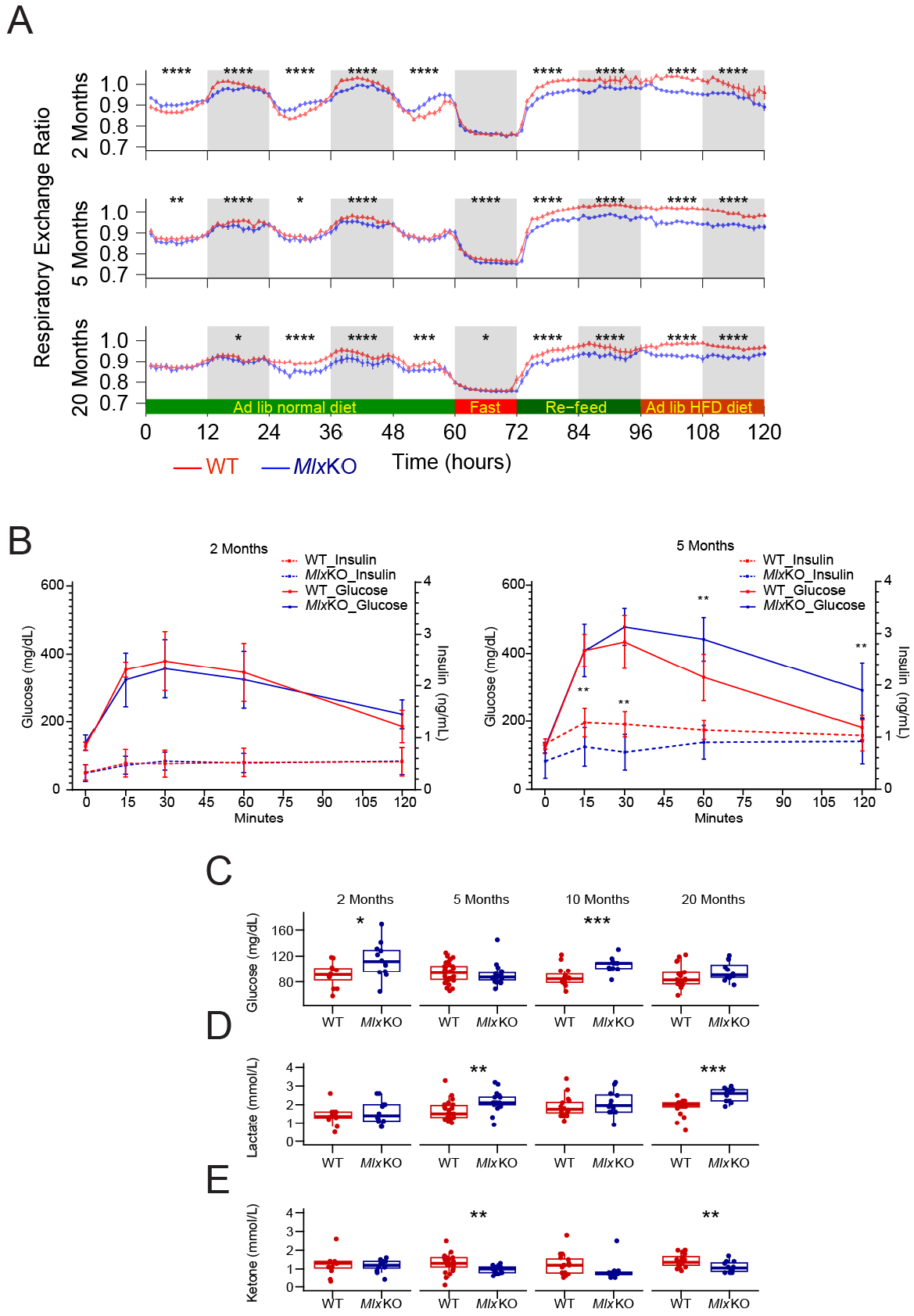
Metabolic alterations in *Mlx*KO mice. **(A)** Respiratory exchange ratios in WT and *Mlx*KO mice performed in metabolic cages at the indicated ages. N=11-12 animals/group. **(B)** GTTs performed on WT and *Mlx*KO mice at the indicated ages. Glucose was administered via i.p injection to fasting animals animals as previously described (*23*). N=5 animals/group. **(C,D,E)** Fasting serum glucose, lactate and ketone levels, respectively performed on WT and *Mlx*KO mice at the indicated ages.

To determine whether the exaggerated FAO dependency of *Mlx*KO mice was a consequence of Type 2 diabetes (T2D)-like insulin insensitivity as previously described in *Myc*KO mice (*23*), glucose tolerance tests (GTTs) were conducted with age-matched WT mice serving as controls. In 2 month old mice, no inter-group differences were observed in the peak glucose levels achieved after glucose administration or in the kinetics of the initial glucose response and its normalization (Figure 3B). Nor did the two groups differ in their insulin kinetics in response to the glucose challenge. By 5 months, however, *Mlx*KO mice demonstrated higher peripheral glucose levels and delayed normalization following the glucose challenge. While mimicking the response previously seen in *Myc*KO mice (*23*), the underlying basis for this transient hyperglycemia was quite different. Rather than the T2D-like exaggerated and prolonged hyperinsulinemia seen in *Myc*KO mice, the insulin response of *Mlx*KO mice was blunted and more akin to that associated with Type 1 diabetes (T1D). Further metabolic profiling showed that *Mlx*KO mice were intermittently prone to the development of fasting hyperglycemia and lactic acidemia (Figure 3C-E). Together with the previous metabolic cage experiments, these studies indicate that *Mlx*KO mice increase their reliance on FAO at least in part because they secrete insufficient amounts of insulin to allow for the adequate uptake of glucose.

Many genes whose encoded proteins contribute to mitochondrial structure and function are co-regulated by both the Myc and Mlx Network members (*13, 24, 29, 41*). Given the above-noted metabolic abnormalities in *Mlx*KO mice, we examined 3 tissues (liver, skeletal muscle and abdominal white adipose tissue) in which mitochondrial function is known to be compromised by *Myc* loss and/or aging (*15, 23, 24, 43, 47–49*). Respirometry studies performed on liver mitochondria from 5 month old WT and *Mlx*KO mice showed attenuated responses in the latter group to non-rate-limiting amounts of all tested substrates for Complex I (malate and pyruvate), Complex II (succinate), and fatty acid oxidation (palmitoyl-coenzyme A) (Figure 4A). In contrast, no differences were seen in the responses of skeletal muscle mitochondria, whereas in white adipose tissue mitochondria, both Complex I and Complex II defects were noted with the former being confined largely to the pyruvate response (Figure 4B and C). Mitochondrial function in tissues of older mice was also similar, with the only noticeable defect being a suppressed response to palmitoyl-coenzyme A in *Mlx*KO livers (Figure 4D-F). The Complex I and Complex II defects in younger *Mlx*KO mice thus resembled those identified previously in association with aging and/or Myc loss in various tissues with the exception of skeletal muscle, which tended to display little, if any, age-related declines in mitochondrial function (*15, 29, 50, 51*).

**Figure 4.**
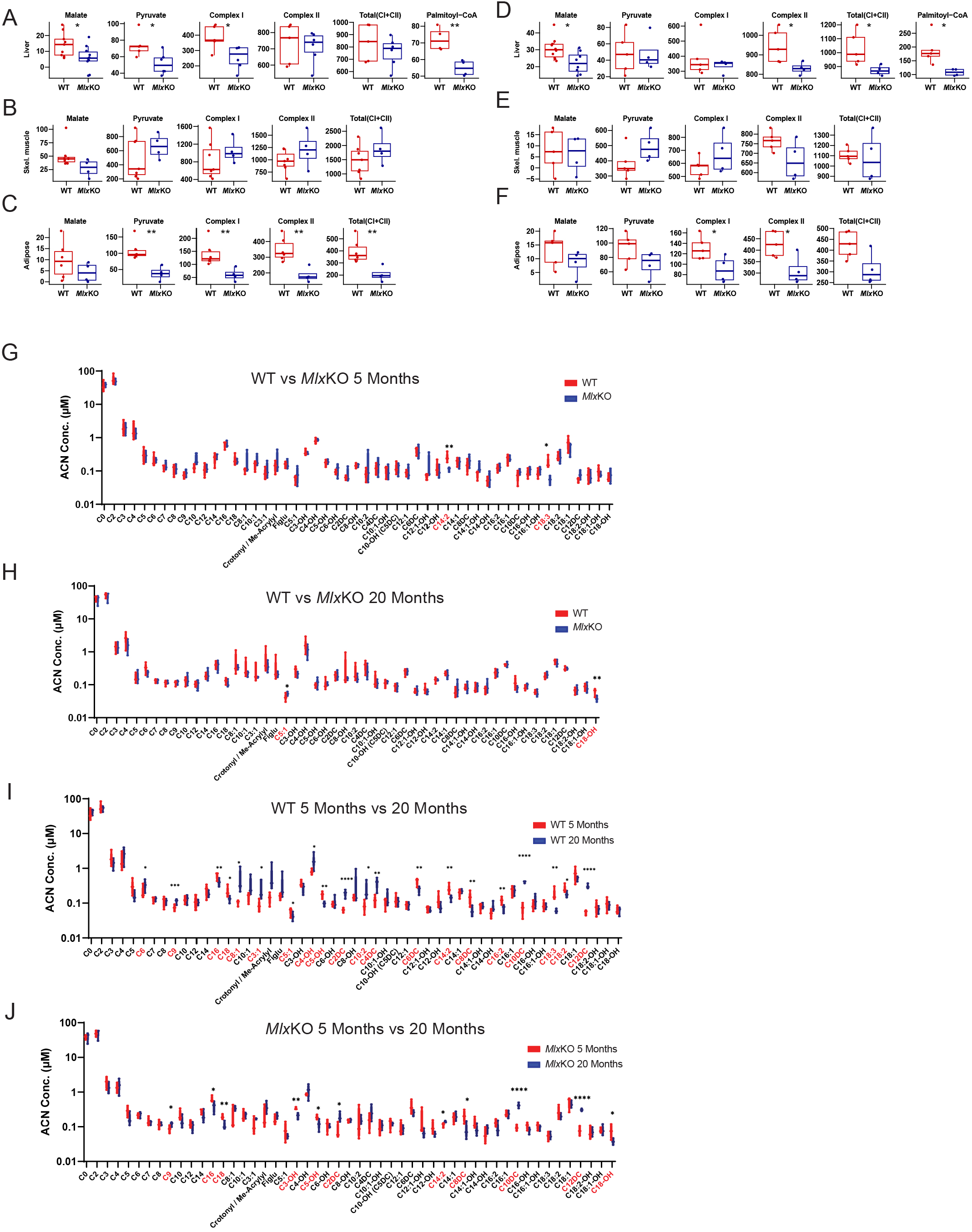
Aberrant Mitochondrial Function in *Mlx*KO tissues. **(A,B,C)** Respirometry studies on mitochondria from liver, adipose tissue and skeletal muscle, respectively of 5 month WT and *Mlx*KO mice in response to malate, pyruvate, palmitoyl-CoA and total Complex I and Complex II activities. **(D,E,F)** Respirometry studies as described in A-C performed on 20 month old mice **(G,H)** Serum acyl carnitine levels in 5 month old and 20 month old WT vs. *Mlx*KO mice, respectively. Significant differences between the two groups are highlighted in red along the horizontal axis. **(I,J)** Comparison of serum acyl carnitine levels in 5 month old vs. 20 month old WT and *Mlx*KO mice, respectively. Significant differences between the two groups are highlighted in red along the horizontal axis.

Serum acyl carnitine measurements can serve as surrogates for body-wide mitochondrial dysfunction and are useful for diagnosing Complex I defects (*52*). Mass spectrometry-based quantification of 51 serum carnitines revealed lower levels of C14:2 and C18:3 in 5 month old *Mlx*KO mice (Figure 4G and Supplementary Table 2). Rather than indicating mitochondrial dysfunction, this finding suggested a more rapid mitochondrial uptake and preferential utilization of these long-chain fatty acids for FAO as documented by metabolic cage profiling (Figure 3). The failure of these disparities to persist in older mice (Figure 4H) could indicate that the source of fatty acids changed over time, from those circulating in serum to those that had accumulated in tissues and thus were potentially more readily accessible (Figure 2A).

Age was more strongly associated with changes in serum acyl carnitines than was *Mlx* status and, in both cohorts, tended to reflect the loss of mitochondrial efficiency in general and Complex I specifically in older mice (Figure 4I and J and Supplementary Tables 2-5) (*47, 48, 51*). A tendency for age-related increases in serum dicarboxylic acids may have indicated a preferential shunting of serum fatty acids into non-energy-generating peroxisomal and lysosomal FAO pathways as seen in inborn errors of long chain fatty acid oxidation, while maintaining a reliance on tissue stores of fatty acids for mitochondrial utilization(*53*).

Compromised Myc and Mlx function can lead to structural and/or functional defects in the electron transport chain (ETC) along with altered mitochondrial size, number, ultrastructure and ATP generation (*15, 18, 24, 44*). Blue native gel electrophoresis (BNGE) and *in situ* enzymatic evaluations of ETC Complexes I, III, and IV from WT and *Mlx*KO livers of 5 month old mice revealed no overt differences between the two groups (Figure 5A). In contrast, the ATPase activity of Complex V was reduced nearly 4-fold in *Mlx*KO livers (Figure 5B). Similar studies on mitochondria from skeletal and cardiac muscle of these same animals revealed no differences thereby again indicating that the consequences of *Mlx* inactivation in mice are highly tissues-specific, as they are with *Myc* (Figure 5C-F) (*23*).

**Figure 5.**
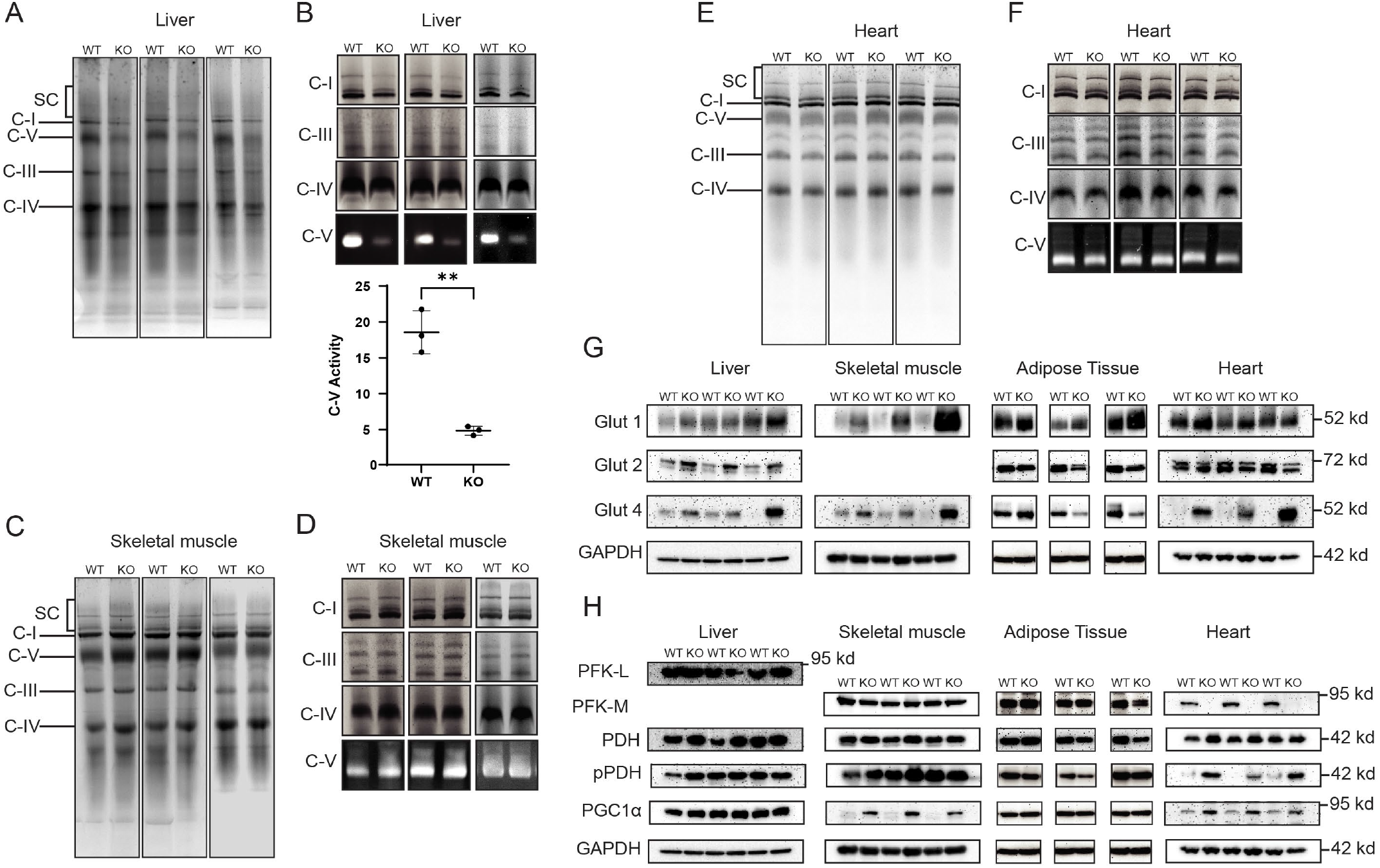
*Mlx*KO tissues are associated with selective changes in mitochondrial function. **(A)** BNGE of purified ETC Complexes I-V and ATP synthase (Complex V) from WT and *Mlx*KO livers of 5 month old mice. SC: supercomplexes. **(B)** *In situ* enzymatic activities of the Complexes shown in **A** (*18*). In the graph below the image, relative specific activities of Complex V were quantified by normalizing scans of enzymatic reactions to the Coomassie Blue-stained bands in **A**. **(C)** BNGE of purified ETC Complexes I-V and ATP synthase (Complex V) from WT and *Mlx*KO skeletal muscle of 5 month old mice. **(D)** *In situ* enzymatic activities of the Complexes shown in **C**. **(E)** BNGE of purified ETC Complexes I-V and ATP synthase (Complex V) from WT and *Mlx*KO hearts of 5 month old mice. **(F)** *In situ* enzymatic activities of the Complexes shown in **E**. **(G)** Immunoblots of select glucose transporters in the indicated tissues from 5 month old WT and *Mlx*KO mice. The absence of a Glut2 panel for skeletal muscle indicates that the protein was not detected in that tissue **(H)** Immunoblots of the indicated factors in WT and *Mlx*KO tissues.

Given the marked differences in fuel preferences, glucose kinetics and ETC substrate responses of *Mlx*KO mice and previously described *Mlx*KO MEFs (Figures 2A and B and 3A and B) (*24*), we evaluated several tissues in 5 month old mice for the expression of relevant glucose transporters, which are known to be differentially regulated by Myc (*17, 23, 24*). We focused on ubiquitously expressed Glut1/Slc2a1, which is particularly prominent in muscle and adipose tissue; Glut 2/Slc2a2, the major glucose transporter in liver; and Glut4/Slc2a4, which is insulin responsive and abundant in skeletal and cardiac muscle and adipose tissues (*54*). Each transporter showed unique changes in *Mlx*KO tissues (Figure 5G). For example, Glut1 was up-regulated in *Mlx*KO skeletal muscle tissue, Glut2 was up-regulated in *Mlx*KO liver and Glut4 was up-regulated *Mlx*KO liver, skeletal muscle and heart. We also examined the expression of several rate-limiting enzymes that control glucose’s fate and mitochondrial biogenesis (Figure 5H). Notably, the liver-specific isoform of phosphofructokinase, PFK-L, was unchanged in *Mlx*KO livers, whereas the muscle-specific isoform, PFK-M, was markedly down-regulated in *Mlx*KO hearts and unaltered in *Mlx*KO skeletal muscle and adipose tissue. Pyruvate dehydrogenase (PDH), which catalyzes the mitochondrial conversion of pyruvate to acetyl coenzyme A was unaltered across all tissues, irrespective of *Mlx* gene status although its activity, as judged by levels of inhibitory phosphorylation at Ser_293_ (pPDH), was decreased in cardiac muscle (Figure 5D). Peroxisome proliferator-activated receptor-gamma co-activator 1 alpha (PGC1α)/PPARGC1A), a transcriptional driver of mitochondrial biogenesis (*55*), was highly induced in *Mlx*KO skeletal muscle. Collectively, these findings broadly implicate *Mlx*KO in tissue-specific metabolic reprogramming involving both oxidative and non-oxidative arms of energy generation, particularly via the regulation of rate-limiting steps in glucose uptake, glycolysis, FAO and mitochondrial biogenesis.

### Transcriptional reprogramming of *Mlx*KO tissues involves subsets of Myc Network-regulated genes

Mlx Network members also bind to numerous Myc target genes (*13, 14, 25, 56*). Using RNA-seq, we compared the liver, abdominal adipose tissue and skeletal muscle transcriptomes from young (5 month old) and old (20 month old) *Mlx*KO and WT mice. Our recent studies on these tissues from *Myc*KO mice of the same ages also allowed us to compare and contrast gene expression profiles between the two KO cohorts (*23*).

We first used Gene Set Enrichment Analysis (GSEA) to verify that a previously authenticated 282-member Chrebp/MondoA/Mlx target gene set from the EnrichR database and a 157-member direct target set from the Qiagen IPA collection were dysregulated as expected in all 3 tissues from young *Mlx*KO mice (Supplementary Figure 2A and B) (*23*). While the magnitude of this dysregulation and the identities of individual affected genes varied among the tissues, the findings validated their use for all subsequent analyses. Consistent with previous findings that some Mlx Network target genes are also direct Myc Network targets (*23–25, 31, 56*) many of the transcripts from the above gene sets were also altered in the corresponding tissues from 5 month old *Myc*KO mice, albeit to a somewhat lesser degree (Supplementary Figure 2C and D). These results confirmed that *Mlx* inactivation was associated with the anticipated tissue-specific changes in the expression of its target genes. They also documented significant and complex cross-talk between the Mlx and Myc Networks, with some of the genes being responsive to Mlx only, others to Myc only and others being duly responsive to both factors (*13, 14, 23, 24*).

Three unbiased analytic methods (CLC Genomics, DeSeq and EdgeR) were used to capture gene expression differences between the above age-matched *Mlx*KO and WT tissues while ensuring the highest degree of confidence in their selection. Only transcripts identified as being differentially expressed by all 3 methods (q<0.05) were included in our final analysis. In 5 month old *Mlx*KO mice, 150 individual transcripts in livers met this criterion. Of these, 60 (40%) were previously identified as putative direct Mlx and/or MondoA targets based on a compilation of Chip-Seq results from 3 available human cell lines in the ENCODE database (HepG2 hepatoblastoma, K562 chronic myelogenous leukemia/lymphoid blast crisis and WTC11 induced pluripotent stem cells) (*57*). In adipose tissue, 2163 such differences were noted, 891 of which (41.2%) were direct Mlx/MondoA targets and in skeletal muscle, only 3 differences were noted, with 2 being direct targets. An initial analysis using Ingenuity Pathway Analysis (IPA) indicated that these 2316 genes were most commonly involved in oxidative phosphorylation (Oxphos), translation and responses to glucose and other nutrients as we and others have previously observed (*7, 13, 14, 24–26*).

GSEA was next used to expand these findings by identifying functionally related and coordinately dysregulated gene sets. Doing so in *Myc*KO tissues had previously revealed 7 such major categories, each comprised of multiple gene sets involved in ribosomal and mitochondrial structure and function, DNA damage response/repair, oxidative stress response, aging, senescence and mRNA splicing (*23, 24*). These categories were also identified in *Mlx*KO tissues, albeit with a different collection of gene sets and enrichment scores (Figure 6A, Supplementary Figures 3-9 and Supplementary File 1). A notable finding in the “DNA damage response/repair” category was that the enriched gene sets in *Mlx*KO tissues tended to be more restricted to those involving the recognition and repair of UV radiation-induced DNA damage whereas enriched gene sets in *Myc*KO tissues were more diverse, involving the repair of base-pair mis-matches, inter– and intra-strand cross-links, single– and double-stranded DNA breaks and shortened telomeres (*23, 24*).

**Figure 6.**
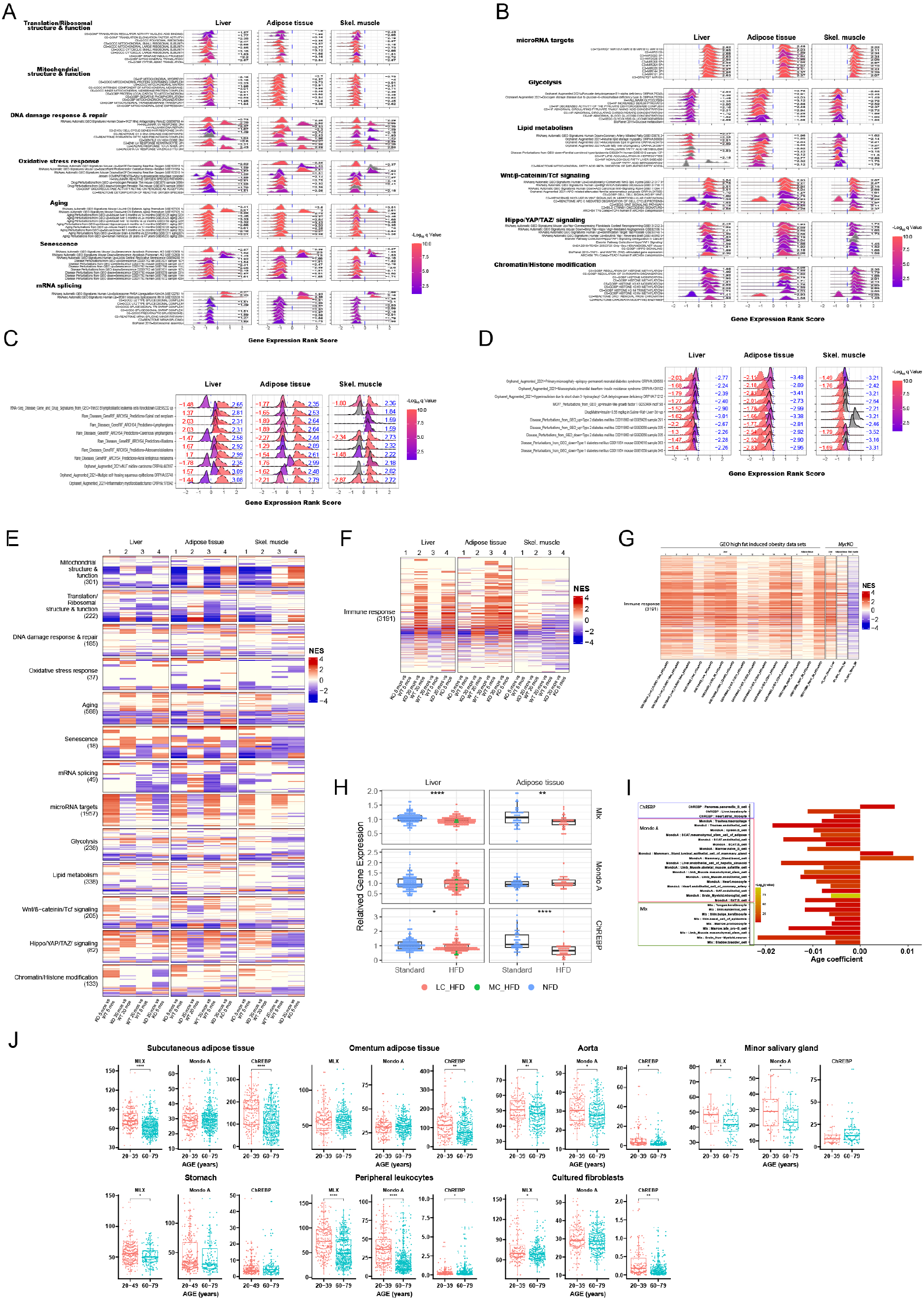
Transcriptional analysis of *Mlx*KO tissues identifies unique targets as well as those previously identified as being Myc-regulated. **(A)** The EnrichR and MSigDB databases were searched to identify gene sets in *Mlx*KO liver, abdominal adipose tissue and skeletal muscle that were selectively enriched relative to WT mice (N=5 tissues/group). Clusterprofiler and the ridgeline plot application tool (*80, 81*). were used to display representative examples of these differentially enriched gene sets from 7 functional categories whose component gene sets were enriched in the same tissues from 5 month old *Myc*KO mice. Numbers to the right of each profile indicate the normalized enrichment score (NES) for that gene set. Gray curves and the absence of an NES indicate gene sets that were not significantly enriched in that tissue but were enriched in at least one other. NESs >0 represent gene sets that were up-regulated relative to their corresponding WT tissue whereas NESs <0 indicate gene sets that were up-regulated. See Supplementary Figures 3-9 for actual GSEA profiles for these and other gene sets and Supplementary File 1 for a complete list of all significantly enriched gene sets identified from the above-mentioned tissues of *Mlx*KO mice. **(B)** Ridgeline plots of individual gene sets, comprising 6 functional categories, selectively enriched in the indicated *Mlx*KO tissues (See Supplementary Figures 10-15 for actual GSEA profiles for these and other gene sets contained within these categories and Supplementary File 2 for a complete list of all significantly enriched gene sets. **(C)** Ridgeline plots of individual gene sets related to cancer in each of the indicated *Mlx*KO and *Myc*KO tissues. Dotted curves, whose NESs are indicated in blue, represent the gene set profiles from *Mlx*KO tissues and solid curves, whose NESs are indicated in red, represent gene set profiles from *Myc*KO tissues. See Supplementary Figures 16 and 17 for actual GSEA profiles and Supplementary file 3 for a list of all cancer-related gene sets identified as being enriched in the above tissues. **(D)** Ridgeline plots for individual gene sets dysregulated in T1D and T2D represented as shown in **C**. **(E)** Heat map of gene sets from the categories shown in panels **A** and **B** along with a more extensive collection of gene sets with significant enrichment scores. The total number of gene sets in each category in indicated beneath its name and their identities are indicated in Supplementary File 4. **(F)** Heat map of 3191 gene sets pertaining to the immune response, inflammation and cytokine production that were significantly dysregulated in *Mlx*KO tissues from 20 month old mice versus matched WT tissues. **(G)** Enrichment of differentially expressed gene sets from **F** in liver, adipose tissue and skeletal muscle from adult mice maintained on standard or various types of HFDs for 10-24 weeks. The right-most 3 columns show GSEA differences between WT and *Myc*KO tissues (n=5 tissues/group) (*23*). **(H)** Mlx, MondoA and ChREBP transcript levels in the livers and adipose tissues from mice maintained on the standard diets or HFDs described in **G**. **(I)** scRNA-seq results for MondoA, ChREBP and Mlx expression from the Tabula Muris Consortium database (*61*) consisting of 90 sc populations from 23 different tissues of young (1-3 months) and old (18-30 months) mice. Results are expressed as q values that were calculated from correlation coefficients that compared transcript levels across aging populations. **(J)** Age-related declines in MondoA, ChREBP and Mlx transcripts in select human tissues and cultured primary fibroblasts. RNA seq results were divided into those obtained from younger (20-49 years) and older (60-79 years) individuals.

In addition to the above categories that were shared with *Myc*KO tissues, we identified 6 others that were more prominent in or unique to *Mlx*KO tissues. The gene sets comprising them encoded miRNA targets or proteins participating in glycolysis, lipid metabolism, Wnt/β-catenin/Tcf signaling, Hippo/YAP/TAZ signaling and chromatin/histone modification (Figure 6B and Supplementary Figures 10-15 and Supplementary File 2).

GSEA from the above tissues of *Myc*KO mice had previously revealed the suppression of numerous gene sets that are often collectively up-regulated in various cancers (*23*). Because many of the individual genes comprising these sets were also direct Myc targets, this finding was consistent with the 3.4-fold lower lifetime cancer incidence we have reported for these animals (*23*). In contrast, 5 month old *Mlx*KO mice tended to show an up-regulation of these gene sets (Figure 6C, Supplementary Figures 16 and17 and Supplementary File 3). Overall, these results were in agreement with previously proposed contrasting roles for Mlx in tumor suppression versus Myc’s role in tumor promotion (*7, 13, 14, 23, 24, 26*).

An additional category of gene sets previously identified as being enriched in *Myc*KO tissues was comprised of those that associated with either T2D or T1D and was consistent with the T2D-like GTTs and insulin kinetics of these animals (*23*). Conforming to the T1D-like profiles of *Mlx*KO mice (Figure 3B and C), this category of gene sets was dysregulated although the precise identities of the gene sets again differed from those of *Myc*KO mice (Figure 6D and Supplementary Figures 18 and19 and Supplementary File 4).

We next asked how gene expression profiles differed among tissues from 5 month old and 20 month old *Mlx*KO mice. To ensure that these results were again compiled in an unbiased and comprehensive manner, we searched the MSigDB and Enrichr databases for gene sets comprising each of the functional categories shown in Figure 6A-D as well as for novel categories of gene sets (Supplementary File 2). This broader search confirmed the overall directionality of the selected gene sets depicted in Figure 6A-D while revealing as many as 1957 additional gene sets in the “miRNA target” category to as few as 18 in the “Senescence” category (Figure 6E-column 1 and Supplementary File 5). In tissues of 20 month old WT and *Mlx*KO mice, a smaller total number of these enriched gene sets was noted and, in some cases, the direction of the dysregulation was the opposite of that seen at 5 months (Figure 6E-column 2). Comparing the gene sets of 5 month old and 20 month old mice showed many of the above gene sets to be both age– and Mlx-dependent (Figure 6E-columns 3 and 4).

We also determined whether tissues from 20 month WT and *Mlx*KO mice differed in the enrichment of functionally related gene sets that were either not detected during the initial analysis of 5 month old tissues or that comprised minor categories. Doing so identified 3191 dysregulated gene sets across all 3 *Mlx*KO tissues related to the immune response, inflammation and the production of or response to various inflammation/immune-related cytokines such as interferon gamma (IFNγ), tumor necrosis factor alpha (TNFγ), and interleukin-6 (IL-6) (Figure 6F and Supplementary File 6). These gene sets were enriched in highly tissue-specific patterns and were most prominent in liver where 1993 (62.5%) of them were significantly enriched. This was followed by adipose tissue and skeletal muscle where 999 (31.3%) and 649 (20.3%) of the sets were enriched, respectively. These were readily detectable by 5 month of age but were particularly pronounced in older *Mlx*KO mice, most notably in liver and adipose tissue (Figure 6F-column 4).

The above results did not distinguish between direct Mlx-dependent transcriptional targets and secondary ones whose dysregulation was a consequence of the metabolic dysfunction-associated steatotic liver disease (MASLD) associated with *Mlx*KO mice (Figure 2) (*58, 59*). We therefore asked whether any of the gene sets shown in Figure 6F were also enriched in tissues of otherwise normal mice maintained on HFDs. From the GEO database and our own previous study (*60*), we retrieved 12 sets of RNAseq results from livers of normal mice of various strains maintained on standard or HFDs, the latter of which varied in their composition (long-chain vs. medium chain) and/or duration (10-24 weeks). Four additional HFD data sets were analyzed in adipose deposits obtained from different anatomical locations and one HFD data set was obtained from skeletal muscle. Many of the differentially enriched gene sets in these tissues and *Mlx*KO tissues were identical, thereby indicating that the immune-related transcript differences of the latter were attributable to their higher neutral lipid content and/or its associated systemic pro-inflammatory state (Figure 6G and Supplementary File 6). Comparing these gene set profiles from older *Mlx*KO mice and *Myc*KO mice, both of which accumulated comparable degrees of hepatic lipid (*15, 29*), continued to show differences, most notably in liver. Collectively, this analysis, which sought to equalize the contributions made by HFDs, steatosis and system inflammation, indicated that, while a majority of immune-related gene expression differences in *Mlx*KO mice were attributable to these features, other differences were specifically related to the loss of Mlx.

Relative to control mice maintained on standard diets, those on HFDs showed no significant changes in liver-or adipose tissue-associated MondoA transcript levels whereas ChREBP and Mlx transcripts were significantly lower (Figure 6H). Together with our previous findings that mice with hepatocyte-specific loss of ChREBP also develop MASLD in the absence of obesity, these results suggest that the lipid accumulation seen in response to HFDs requires the down-regulation of ChREBP and/or Mlx, which play important roles in lipid and carbohydrate metabolism (*7, 13, 14, 26, 27, 32, 34, 56*).

Myc expression normally declines with age in a number of murine and human tissues (*23*). To determine whether this also occurs with any of the above Mlx Network members, we compared RNA-seq results from 90 single cell (sc) populations originating from 23 tissues from young (1-3 months) and old (18-30 months) mice (Figure 6I) (*61*). Significant age-related declines were noted in MondoA transcripts in 16 sc populations, declines in ChREBP transcripts were noted in 2 sc populations, and declines in Mlx transcripts were noted in 8 sc populations. Examination of total organ transcripts from young and old humans (ages 20-49 and 60-79, respectively) from the Broad Institute’s GTEx database (https://gtexportal.org/home/) also showed age-related declines in MondoA, ChREBP and Mlx expression in several tissues as well as in *in vitro* cultured primary skin fibroblasts (Figure 6J) (*62*). Thus, like Myc transcripts (*23, 40*), those of some Mlx Network members progressively decline in some normal aging tissues. In this setting, the co-regulation of many Myc target genes by Mlx Network members may explain why the previously observed loss of regulation of Myc target genes in response to aging was often more pronounced than was the decline of Myc itself (*23*).

## DISCUSSION

The Mlx Network target gene repertoire, originally thought to regulate glucose and lipid metabolism, has recently been expanded to include the maintenance of mitochondrial structure and function, ribosomal biogenesis and translation and other functions described here and elsewhere (Figure 6A-E) (*7, 14, 26, 32, 34, 40, 56, 63*). Numerous Chip-seq results have demonstrated that Myc and Mlx Network members can compete for the same E box and/or ChoREs in positively-regulated target gene promoters or bind to distinct elements in close physical proximity, with similar co-regulation extending to negatively-regulated target genes (*23, 24, 40, 57, 63*). Indeed, the loss of Mlx Network members may in some cases have a greater impact on the expression of previously identified direct Myc target genes than does the loss of Myc itself and *vice versa* (*23, 24, 28*). For these reasons, and because the target gene sets regulated by these two Networks are not identical, it is perhaps unsurprising that premature aging would be observed in *Mlx*KO mice with the individual phenotypes overlapping those described in *Myc*KO mice (*23*).

*Mlx*KO females achieved their maximal body weights earlier than WT mice while also attaining a higher fat:lean mass ratio although this was not nearly as pronounced as that seen in *Myc*KO mice (Figure 1A) (*23*). The fat content and body mass of these individuals also began their declines earlier than they did in WT mice. Although *Mlx*KO mice did not manifest the premature alopecia or achromotricia that typifies *Myc*KO mice, they did accumulate an overall ∼3-fold higher lifetime incidence of corneal opacities than either WT mice or *Myc*KO mice (Figure 1C) (*23*). Male *Mlx*KO mice also had less endurance when subjected to treadmill testing and both sexes showed reduced diurnal ambulatory activity, although this was again more prominent in females (Figure 1E and G). Together these findings show that *Mlx*KO and *Myc*KO mice develop both common and unique features that are consistent with premature aging.

Despite their various co-morbidities that would be expected to reduce lifespan, *Myc*KO mice actually live up to 20% longer than WT mice (*23*). We have attributed this to their 3.4-fold lower lifetime incidence of cancer, which is the major associated finding at the time of death in most inbred mouse strains and which often requires Myc to initiate and/or maximize neoplastic growth (*1, 13, 14, 45, 64*). In contrast, *Mlx*KO mice of both sexes had the same lifespans as WT mice despite a ∼1.5-1.7-fold higher cancer incidence (Figure 1H and I). This is consistent with previous observations that Mlx is a suppressor of both normal and neoplastic cell growth, as are other Mlx and Myc Network members such as Mxd1, Mxd4 or Mnt (*13, 14, 25, 65–67*). Thus, unlike *Myc*KO mice in which aging and cancer development can be dissociated so as to increase longevity, *Mlx*KO mice retain this link or even strengthen it, due presumably to both the retention of *Myc* and the loss of competition from the *Mlx* Network.

The increased fecal fat content of *Mlx*KO mice (Figure 2D) has not been previously observed in either WT or *Myc*KO mice at any age (*23*). Fat malabsorption in aging humans is common and is often caused by imbalances in the intestinal flora or other factors that alter the uptake of monoglycerides and fatty acids by intestinal enterocytes (*68*). Our inability to discern any consistent histologic changes in either the small or large intestines of *Mlx*KO mice (Figure 2C) indicates that these are unlikely to be causes of their steatorrhea, particularly since we have not observed it in *Myc*KO mice in which more pronounced structural defects of their intestinal vili are observed, albeit transiently following the loss of Myc (*23, 45*). Other non-mutually exclusive possibilities to explain our finding include an inability to synthesize or excrete bile acids and pancreatic exocrine insufficiency. Regardless of the cause, the observed malabsorption was not significant enough to impair normal growth (Figure 1A).

In both mice and humans, the incidence of MASLD normally increases with age and is exacerbated by resistance to or under-production of insulin, both of which force a switch from glucose to fatty acids as the preferred energy-generating substrate (*42, 43*). Age-related declines in Myc and mitochondrial efficiency may be at least partially responsible for this switch and the resulting lipid accumulation that exceeds the amount needed to generate sufficient energy (*15, 47, 50, 51, 69–71*). Normal and neoplastic tissues of non-hepatic origin are also driven to accumulate neutral lipid following the genetic or pharmacologic compromise of Myc (*44*). That MASLD also develops in response to the knockout of *Chrebp* or *Mlx* has been attributed to a similar deregulation of many of the same Myc target genes that support mitochondrial intergrity (*13, 14, 24, 25, 29*). In either case, the greater dependency on FAO as a more efficient means of energy generation may be accompanied by other compensatory changes such as increases in mitochondrial mass (*48, 70, 71*). Changes in circulating acyl carnitines and the selective utilization of fatty acids of different lengths likely reflect the integration of normal age-related changes, those arising as a consequence of Mlx’s absence, and whether the lipid used for FAO derives directly from immediately available dietary sources or pre-existing tissue depots (Figure 4I and J and Supplementary Table 2-5) (*23*).

Disparities in mitochondrial function and fuel selection between WT and *Mlx*KO mice were observed quite early and were reflected in RERs measured during metabolic cage studies (Figure 3A). As with *Myc*KO mice (*23*), this was manifested by a significantly greater reliance on FAO, particularly at night when feeding activity peaks and should provide sufficient glucose for it to be the fuel of choice. Whereas this fatty acid preference was somewhat age-dependent and erratic, a more consistent indicator of it was seen following overnight fasting when re-feeding either normal or HFDs was invariably associated with markedly lower RERs. While this reliance on FAO might have been partially attributable to the mild T1D-like responses of older animals following a glucose challenge, it did not explain the findings in younger mice who had normal GTTs and plasma insulin responses (Figure 3B and E). Thus, the abnormal RERs seen throughout life likely reflect both defects in mitochondrial function as well as the later development of inadequate insulin production. Another similarity between *Mlx*KO and *Myc*KO mice was seen in the youngest individuals (2 months of age) in which nocturnal RERs never exceeded 1 as they did in WT mice (Figure 3A). When not attributable to re-feeding, RERs >1 are a reliable indicator of the high rates of fatty acid synthesis and glucose utilization that are generally associated with rapid post-natal growth and which should be blunted in response to the functional inactivation of both MondoA and ChREBP that accompanies *Mlx* KO (*20, 26*). The failure to note this exaggerated RER in 2 month old *Mlx*KO mice might have indicated that it was offset by their excessive reliance on FAO and/or inefficient fatty acid synthesis (*26*). It may also have been a result of the premature aging of these mice such that this normally transient response had already disappeared by 2 months of age. The inability of *Mlx*KO RERs to ever exceed 1 during times of post-fasting re-feeding where this response was maintained in all but the oldest WT mice, suggested that the primary cause was the high levels of FAO.

GSEA of *Mlx*KO tissues revealed alterations in functionally related gene sets that were consistent with the described phenotypes and behaviors (Figure 6E). Not unexpectedly some of these were also enriched in *Myc*KO tissues and MEFs and *Chrebp*KO livers (*23, 24, 28*). The most highly enriched gene sets within the “DNA Damage Response/Repair” category in *Mlx*KO tissues involved UV-induced DNA damage whereas those pertaining to other forms of DNA damage were less highly enriched than they were in *Myc*KO tissues and MEFs (Figure 6E and Supplementary Figure 5). This was consistent with previous observations concerning the differential sensitivities of WT, *Myc*KO and *Mlx*KO MEFs to agents that induce different types of DNA damage (*24*). In these studies *Myc*KO MEFs were highly resistant to virtually all forms of DNA damage whereas in *Mlx*KO MEFs, the resistance was both less pronounced and more restricted to that involving the anti-metabolite 6-thioguanine and the DNA adduct-forming agent cis-platinum. The seemingly paradoxical chemo-resistance of *Myc*KO MEFs, in contrast to the marked sensitivities of cells with monogenic defects in DNA damage recognition/repair pathways, was attributed to the much more global dysregulation of these numerous pathways, their failure to cross-talk with one another and to communicate effectively with downstream apoptotic effector pathways (*23*).

Another category of gene sets deserving of mention was a large one pertaining to “Cancer” (Figure 6C, Supplementary Figure 16-17 and Supplementary File 3). In *Myc*KO tissues, these tended to be regulated in directions opposite to those in tumors whereas in *Mlx*KO tissues, they were regulated in the same direction (*23*). Both findings were in keeping with the 3.4-fold lower lifetime cancer incidence of the former group of mice and the 1.5-1.7-fold *higher* cancer incidence of the latter and with the Mlx Network’s putative role in tumor suppression (*23, 40, 65–67*). The altered transcriptomic landscapes of *Mlx*KO mice implies that they might be poised for cancer development by virtue of dysregulating important cancer-related genes (including *Myc* itself) and the probable need to acquire fewer subsequent oncogenic “hits” as a result (Figure 1I).

We recently identified age-related declines in Myc expression and an even more pronounced loss of Myc target gene regulation in numerous murine and human tissues and have suggested that they may be directly responsible for at least some aspects of normal aging (*23, 40*). This notion largely derives from the fact that Myc oversees many critical functions that deteriorate in response to and even drive aging including DNA replication, mitochondrial and ribosomal structure and function, protection against reactive oxygen species, DNA damage recognition and repair, rRNA splicing and the maintenance of proper metabolic balance (Figures 5 and 6A) (*17, 19, 23, 24, 42, 43, 48, 72–74*). By analogy, various combinations of Mlx, MondoA and ChREBP transcripts show similar age-related and tissue-specific declines in their expression (Figure 6H-J). Given that, at least in fibroblasts, over 60% of Mlx-responsive genes are also Myc-regulated (*24*), it is not surprising that many of the same cellular functions are affected by body-wide loss *Mlx* leading to similar premature aging phenotypes as well as others that reflect the specificities of each Network’s functions.

## Materials and Methods

### Mouse maintenance, breeding and body-wide deletion of the *Mlx* locus

All care, breeding, husbandry and procedures were approved by The University of Pittsburgh Department of Laboratory and Animal Resources (DLAR) and the Institutional Animal Care and Use Committee (IACUC). Unless stated otherwise, mice were maintained on standard animal chow diets and water, both provided *ad libitum*. C57Bl6 mice with LoxP sites flanking exons 3 and 6 of the *Mlx* locus were a generous gift of Dr. R.N. Eisenman (Supplementary Figure 1A-C)(*24*). They were crossed with the B6.129-*Gt(ROSA)26Sortm1(cre/ERT2)Tyj*/J strain (Jackson Labs. Bar Harbor, ME), which expresses the Cre recombinase-estrogen receptor (CreER) fusion transgene driven by the ubiquitously-expressed ROSA26 promoter (*23, 24*). At the time of weaning and upon reaching a weight of at least 15 g (∼4 wks of age), mice containing either one or 2 copies of the CreER transgene were subjected to daily i.p. injections of tamoxifen (75 mg/Kg) in mineral oil for 5 days as previously described (*23*). Two wks later, randomly chosen animals from each group were selected and tissues were examined to determine the efficiency and tissue distribution of *Mlx* locus excision (Supplementary Figure 1E). Subsequent evaluations were performed throughout life to ensure the maintenance of the knockout state (Supplementary Table 1). Tamoxifen-treated offspring of Mlx^fl/fl^ or Myc^fl/fl^ (*23*) mice with intact Mlx and Myc genes served as wild-type (WT) controls.

### Body fat and lean mass determinations

Beginning at the time of weaning, body weights were determined weekly until the age of 4-5 months and then every 2-3 weeks thereafter. In parallel with these studies, the fractional lean and fat content were determined using an Echo-MRI^TM^ scanner (EMR-055, version 160301, Echo Medical Systems, Inc. Houston, TX) as previously described (*23*).

### Strength, balance and endurance testing

A Grip Strength Meter (Harvard Apparatus, Holliston, MA) was used to assess strength at various ages as described previously (*23*). Balance was measured using a Rotarod apparatus (SPW Industrial, Laguna Hills, CA) and a modification of the procedure provided by Jackson Laboratories (Bar Harbor, ME) as described previously (*23*). Treadmill endurance was followed with a Columbus Instruments Exer 3/6 apparatus (Columbus, OH) as described previously (*23*).

### Glucose tolerance tests (GTT) and serum glucose, lactate and ketone quantification

These studies were performed as previously described (*23*). For GTTs, mice were fasted for 5 hour and then injected i.p. with 2g of dextrose/kg body mass. Serum insulin levels were quantified with an Ultra Sensitive Mouse Insulin ELISA Kit (Crystal Chem, Elk Grove Village, IL). Whole blood glucose lactate and ketone levels were also measured in fasted mice (Glucose AimStrip Plus, Germaine Laboratories, Inc. San Antonio, TX; Lactate Plus Analyzer, Sports Resource Group, Inc., Hawthorne NY; Keto-Mojo Ketone Meter, Keto-Check, Inc. Napa, CA).

### Mitochondrial respirometry

Oxygen consumption rates (OCRs) were determine on ∼50 mg of disrupted tissue immediately after sacrifice using an Oroboros Oxygraph 2k instrument (Oroboros Instruments, Inc., Innsbruck, Austria) as described elsewhere (*23, 29, 60*). Baseline OCR values were determined in 2 mL of Mir05 buffer following the addition of non-rate-limiting concentrations of cytochrome c (final concentration=10 μM), malate (2 mM), ADP (5 mM), pyruvate (5 mM), glutamate (10 mM) and palmitoyl-coenzyme A (3 μM). Collectively, these provided an estimate of the Complex I activity whereas the addition of succinate (10 mM final concentration) provided an estimate of Complex II (succinate dehydrogenase) activity. Rotenone (0.5 μM final concentration) was used to inhibit Complex I and to calculate the proportional contributions of Complexes I and II. To measure β-FAO, reactions were primed with ADP (5 mM) and malate (5 mM) and then supplemented with palmitoyl-CoA (3 μM) and L-carnitine (10 μM). Activities were normalized to total protein.

### Blue native gel electrophoresis (BNGE) and *in situ* enzymatic assays of ETC complexes and ATP synthase

These evaluations were performed as described previously (*15, 18*). Briefly, 50-100 mg of fresh liver was placed into 0.5 ml of an ice-cold solution containing 25 mM Tris-HCl, pH 7.5; 100 mM KCl; 0.4 M sucrose and supplemented with protease inhibitor cocktail (Sigma-Aldrich, Inc. St. Louis, MO). Tissues were then disrupted by homogenization (Isobiotec, Heidelberg, Germany) followed by centrifugation at 500 × *g* for 10 min at 4°C. The mitochondria-rich supernatant was further centrifuged at 14,000× *g* for 15 min. The pellet was washed twice with the above buffer and then re-suspended at a final protein concentration of 5 mg/ml. 8 mg of the suspension was disrupted with digitonin (MP Biomedicals, Solon OH) so as to provide a final ratio of protein:digitonin of 1:8 (*15, 18*). After incubating on ice for 20 min, Coomassie blue (5% Coomassie blue G250 in 750 mM 6-aminocaproic acid) was added (1/20 v/v), and the mixture was clarified by centrifugation at 14,000 x g for 20 min at 4°C. The supernatant was then electrophoresed on a 3-12% Native PAGE Novex Bis-Tris gel (Invitrogen, Inc. Waltham, MA) at 80 V for 4 hours at 4°C in the supplier’s buffer. Gels were then stained with Bio-Safe Coomassie G250 (Bio-Rad, Hercules, CA) for 30 min and exhaustively de-stained with water. Scanning and analysis for band densities were performed using an AlphaEaseFC 2200 scanner and AlphaEaseFC software. To quantify Complex I activity (NADH ubiquinone oxidoreductase), gels were incubated in 3–4 ml of 2 mM Tris-HCl, pH 7.4; 2.5 mg/ml nitrotertrazolinum blue chloride and 0.1 mg/ml NADH at 37°C for 1–2 hours before quantifying band intensities by densitometric analysis. An average value from 3-4 gels was calculated. Complex III (CIII) (decylubiquinol cytochrome c oxidoreductase) activity was assayed by incubating gels with CIII assay solution overnight with mild agitation (*18, 23*). Complex IV (CIV) (cytochrome c oxidase) activity was measured by incubating the gel in 1 nM catalase, 10 mg cytochrome c and 750 mg sucrose in CIII assay buffer with mild agitation for 30 min followed by a final overnight wash in water. Complex V was quantified by measuring ATPase activity. Gels were incubated in 3–4 ml of 34 mM Tris-glycine, pH 7.8; 14 mM MgSO_4_; 0.2% Pb(NO_3_)_2_ and 8 mM ATP for at least 3 hours at 37°C with any further incubation being performed at room temperature to further strengthen band intensities.

### SDS-polyacrylamide gel electrophoresis (SDS-PAGE) and immunoblotting

After sacrifice, tissues were removed from WT and *Mlx*KO mice, immediately placed on ice and then divided into small sections. They were then snap-frozen in liquid nitrogen and transferred to a –80C freezer for subsequent storage. For SDS-PAGE, frozen tissue fragments were disrupted in protease inhibitor-containing SDS-PAGE loading buffer with Bullet Blender as previously described (*23*). Protein concentrations were measured with the Pierce™ BCA Protein Assay Kit (23227, Thermo Fisher Scientific). Electrophoresis, blotting to PVDF membranes and protein detection was performed as previously described (*23*). Antibodies used for the detection of specific proteins were used largely to the directions of the suppliers and are shown in Supplementary Table 6.

### Oil Red O (ORO) staining and triglyceride content of liver samples

The neutral lipid content of livers was determined as previously described (*23*). Briefly ORO-stained liver sections were viewed with a Leica DFC7000T microscope and overlapping images were joined using the stitching plugin program of FIJI software (*23*). Color de-convolution (*75*)was also performed in FIJI where the colors were specified in advance from ROIs respectively corresponding to strongly stained and unstained tissue and background. Oil-Red-O quantification was performed as previously described (*23*). Tissue triglyceride assays were performed as previously described using a Triglyceride Quantification Colorimetric/Fluorimetric Kit (Cat. no. MAK266 Sigma-Aldrich, Inc.) (*23*).

### Measurement of fecal fat

Mice were individually caged and provided with standard chow diets containing 4.5% fat (Picolab 5053; LabDiet, St. Louis, MO, USA). Following a 24 hr period of acclimation, feces were collected over the course of the next 48-72 hours, combined, and stored at –20C until processing. Approximately 200 mg of dried sample was re-suspended in 1 ml of PBS. Total lipids were then extracted using chloroform:methanol, dried and quantified as described after adjusting to the original weight of the sample (*76*).

### Metabolic cage profiling and determination of respiratory exchange ratios (RERs)

These studies were performed as described previously (*23*). Briefly, approximately 12 mice from each group, comprised of equal numbers of both sexes, were maintained individually in metabolic cages (Columbus Instruments, Inc. Columbus, OH) while being provided *ad lib* access to water and the standard mouse chow containing 5% fat described above but in powdered form. After a 48 hr acclimation period, VO_2_ and VCO_2_ were recorded every 20 min over the ensuing 48 hr period as were food and water intake and total activity. At the conclusion of these measurements, mice were fasted overnight (12 hr) and then provided with a standard diet for 24 hr followed by a HFD (45%) for an additional 24 hr while again monitoring RERs. Data analyses were performed with the CalR Analysis online software package (https://calrapp.org/) (*77*).

### Nucleic acid isolation, qPCR and qRT-PCR procedures

Tissue DNAs and RNAs were purified using DNeasy and RNeasy tissue extraction kits, respectively, according to the protocols supplied by the vendor (Qiagen, Inc., Germantown, MD). Sample quantification and purity were determined using a NanoDrop Microvolume Spectrophotometer (Thermo Fisher Scientific, Pittsburgh, PA). A quantitative TaqMan-based qPCR assay to determine the efficiency of *Mlx* locus inactivation was performed using the strategy described in Supplementary Figure 1A-D. A TaqMan-based qRT-PCR strategy was also employed to quantify the ratio of exon 1 transcripts to exon 6 transcripts in WT and *Mlx*KO tissues and to further confirm the extent of *Mlx* locus inactivation (Supplementary Figure 1E). Reverse transcription was performed using a SuperScript IV 1^st^ Strand cDNA Synthesis Kit (Life Technologies, Inc.) under the conditions recommended by the vendor. All oligonucleotide primers and probes were synthesized by IDT, Inc. (Coralville, IA) based on sequences obtained from the *Mlx* GenBank DNA sequence: NC_000077.7 (100977538..100983033) (Supplementary Figure 1D). All PCR and RT-PCR reactions were performed on a CFX96 Touch^TM^ real-time PCR detection system (Bio-Rad Laboratories, Inc. Hercules, CA).

### RNAseq and transcriptomic analyses

RNAseq and transcriptomic analyses were conducted on liver, skeletal muscle, and abdominal white adipose tissue RNAs obtained from WT and *Mlx*KO mice of specified ages (5 per group). Total RNA purification was performed simultaneously for all samples using the QIAGEN RNeasy Mini Kit (QIAGEN, GmbH, Hilden, Germany), followed by DNase digestion utilizing the TURBO DNA-free™ Kit (Thermo Fisher Scientific Inc.) (*23, 29, 60*). RIN values were assessed using an Agilent 2100 Bioanalyzer (Agilent Technologies, Santa Clara, CA), and samples with RIN values ≥8.5 were selected for further processing. The sequencing procedures were carried out on a NovaSeq 600 instrument by Novagene, Inc. as previously described (*1, 23, 29, 60*), and the raw data were deposited in the National Center for Biotechnology Information (NCBI) Gene Expression database under the accession number GSE248073. This data is accessible through the Gene Expression Omnibus (GEO).

Differentially expressed transcripts were identified using three methods: CLC Genomic Workbench version 23.0 (QIAGEN), EdgeR, and DeSeq2, as previously described (*23, 24*). For the latter two methods, raw reads from FASTQ files were mapped against the GRCm38.p6 mouse reference genome using nf-core/rnaseq v3.12.0. FeatureCounts from this analysis served as input for EdgeR and DeSeq2 analyses.

Ingenuity Profiling Analysis (IPA) was utilized for classifying transcripts into pathways, with significance adjusted for false discovery using the Bonferroni–Hochberg correction. Gene Set Enrichment Analysis (GSEA) was employed to detect alterations in the collective expression of functionally-related sets of transcripts. This analysis was performed using the clusterprofiler package in R. These gene sets were obtained from the Enrichr Database (https://maayanlab.cloud/Enrichr/#libraries) and the Molecular Signatures Database (MSigDB) v.7.2 (http://www.gsea-msigdb.org/gsea/msigdb/index.jsp). Heatmaps were generated using the ComplexHeatmap R package. Additionally, RNAseq results obtained from liver, skeletal muscle, and adipose tissues of 5-month-old and 20-month-old MycKO mice were extracted from previously published work and are accessible through the GEO accession number GSE223676 (https://www.ncbi.nlm.nih.gov/geo/query/acc.cgi?acc=GSE223676) (*23*).

### Bioinformatics analyses and database searches

The mouse single-cell RNAseq data used to assess the expression of Mlx, MondoA (Mlxip), and ChREBP (Mlxipl) were obtained from https://figshare.com/ndownloader/files/27856758. These data were analyzed for gene and aging correlations (*78*). The aging correlation of Mlx, MondoA (Mlxip), and ChREBP (Mlxipl) was examined across various tissue cells. Significant tissues were identified and visualized using ggplot2.

Transcript levels for Mlx, MondoA (Mlxip), and ChREBP (Mlxipl) in tissues from young and elderly humans were collected from the GTEx Portal (GTEx Analysis V8 release: RNAseq gene TPMs by tissue) at https://gtexportal.org/home/datasets (dbGaP: phs000424.v8.p2).

Additionally, RNA-seq datasets for normal mice of different strains, maintained on either standard or high-fat diets (HFDs), were downloaded from GEO databases. Detailed information is available in Supplementary file 7. Signal-to-noise rank lists were generated from the normalized transcription levels using GSEA v4.3.2 (*79*) for gene set enrichment analysis.

### Statistical analyses

All statistical analysis used GraphPad Prism v9.00 (GraphPad Software Inc., USA) and R software v4.2.0 (R Foundation for Statistical Computing, Vienna, Austria) as previously described (*23*). ggplot2 and ComplexHeatmap packages were used for boxplot and heatmap visualizations and the survminer package was used to plot survival curves. A 2-tailed, unpaired t-test was used to assess significant differences between normally distributed populations, and a 2-tailed Mann-Whitney exact test was used for the analysis of non-normally distributed populations.

## Author contributions

The analysis of RNAseq and other genomics data was, in part, supported by the University of Pittsburgh Center for Research Computing. Specifically, this work utilized the HTC cluster, which is funded by NIH award number S10OD028483. Additionally, funding for this project was provided by NIH grant RO1 CA174713, The Rally Foundation (grant no. 22N42), and a Hyundai Hope on Wheels Scholar grant, all awarded to EVP.

## Data and materials availability

All data needed to evaluate the conclusions in the paper are present in the paper and/or the Supplementary Materials. All raw RNA-seq files have been deposited in the NCBI Gene Expression Omnibus and are accessible through GEO Series accession number GSE248073 and GSE223676.

