## Supplemental Table 1 to 6; Supplemental Figure 1 to 19 for "The Myc-Like Mlx Network Impacts Aging and Metabolism"

Science Advances
AAAS

### Supplementary Materials for

#### The Myc-Like Mlx Network Impacts Aging and Metabolism

Huabo Wang *et al.*

**This PDF file includes:**

Supplementary Tables 1 to 5
Supplementary Figures 1 to 19

**Other Supplementary Materials for this manuscript include the following:**

Supplementary Files 1 to 7

**Supplementary Table 1. Efficiency and Persistence of *Mlx* Gene Excision and Reduction of Expression Relative to Comparably Aged WT Tissues**

| Mouse ID | Age (months) | Tissue | qPCR (% knockout) | qRT-PCR (% expression vs. WT) |
|---|---|---|---|---|
| 1 | 2 | Liver | 93 | |
| 2 | 2 | Skeletal muscle | 71 | |
| 3 | 2 | Lung | 80 | |
| 5 | 2 | Kidney | 77 | |
| 6 | 2 | Stomach | 82 | |
| 7 | 2 | Small intestine | 91 | |
| 8 | 2 | Large intestine | 94 | |
| 10 | 2 | Pancreas | 92 | |
| 11 | 2 | Spleen | 95 | |
| 12 | 2 | Bone marrow | 100 | |
| 14 | 2 | Liver | 89 | |
| 15 | 2 | Skeletal muscle | 89 | |
| 16 | 2 | Lung | 80 | |
| 17 | 2 | Heart | 73 | |
| 18 | 2 | Kidney | 79 | |
| 19 | 2 | Stomach | 85 | |
| 20 | 2 | Small intestine | 95 | |
| 21 | 2 | Large intestine | 95 | |
| 23 | 2 | Pancreas | 91 | |
| 24 | 2 | Spleen | 97 | |
| 25 | 2 | Bone marrow | 100 | |
| 52 | 5 | Liver | | 2 |
| 53 | 5 | Skeletal Muscle | | 26 |

| | | | |
|---|---|---|---|
| 54 | 5 | Lung | 21 |
| 55 | 5 | Heart | 30 |
| 56 | 5 | Kidney | 11 |
| 57 | 5 | Stomach | 8 |
| 58 | 5 | Small Intestine | 2 |
| 59 | 5 | Large Intestine | 0.6 |
| 60 | 5 | Brain | 51 |
| 61 | 5 | Pancreas | 10 |
| 62 | 5 | Spleen | 8 |
| 63 | 5 | Bone Marrow | 3 |
| 64 | 5 | Adipose tissue | 2 |
| 65 | 5 | Liver | 1 |
| 66 | 5 | Skeletal Muscle | 29 |
| 67 | 5 | Lung | 12 |
| 68 | 5 | Heart | 17 |
| 69 | 5 | Kidney | 7 |
| 70 | 5 | Stomach | 3 |
| 53 | 5 | Small Intestine | 0.2 |
| 54 | 5 | Large Intestine | 2 |
| 55 | 5 | Brain | 44 |
| 56 | 5 | Pancreas | 2 |
| 57 | 5 | Spleen | 5 |
| 58 | 5 | Bone Marrow | 2 |
| 59 | 5 | Adipose tissue | 1.5 |
| 4299 | 5 | Skeletal Muscle | 40 |
| | | Liver | 1 |

| | | | | |
|---|---|---|---|---|
| | | Adipose tissue | | 11 |
| 5001 | 5 | Skeletal Muscle | | 23 |
| | | Liver | | 2 |
| | | Adipose tissue | | 5 |
| 5238 | 5 | Skeletal Muscle | | 48 |
| | | Liver | | 1 |
| | | Adipose tissue | | 6 |
| 5239 | 5 | Skeletal Muscle | | 30 |
| | | Liver | | 2 |
| | | Adipose tissue | | 9 |
| 27 | 5 | Liver | 84 | |
| 31 | 5 | Kidney | 86 | |
| 32 | 5 | Stomach | 71 | |
| 33 | 5 | Small intestine | 93 | |
| 34 | 5 | Large intestine | 99 | |
| 36 | 5 | Pancreas | 70 | |
| 37 | 5 | Spleen | 99 | |
| 38 | 5 | Bone marrow | 93 | |
| 42 | 5 | Lung | 79 | |
| 43 | 5 | Heart | 54 | |
| 46 | 5 | Small Intestine | 99 | |
| 47 | 5 | Large Intestine | 99 | |
| 49 | 5 | Pancreas | 58 | |
| 50 | 5 | Spleen | 99 | |
| 51 | 5 | Bone Marrow | 99 | |

| | | | | |
|---|---|---|---|---|
| 7236 | 18 | Adipose tissue | | 9 |
| 7237 | 18 | Adipose tissue | | 10 |
| 7238 | 18 | Adipose tissue | | 18 |
| 7239 | 18 | Adipose tissue | | 15 |
| 7240 | 18 | Adipose tissue | | 14 |
| 7116 | 18 | Liver | | 0 |
| 7117 | 18 | Liver | | 4 |
| 7118 | 18 | Liver | | 4 |
| 7119 | 18 | Liver | | 2 |
| 7174 | 18 | Liver | | 2 |
| 7356 | 18 | Skeletal muscle | | 61 |
| 7357 | 18 | Skeletal muscle | | 44 |
| 7358 | 18 | Skeletal muscle | | 41 |
| 7359 | 18 | Skeletal muscle | | 30 |
| 7360 | 18 | Skeletal muscle | | 52 |
| 5521 | 19 | Bone marrow | 95 | |
| | | Adipose tissue | 89 | |
| | | Large intestine | 88 | |
| | | Pancreas | 78 | |
| | | Small intestine | 91 | |
| | | Spleen | 82 | |
| 5538 | 19 | Adipose tissue | 90 | |
| | | Liver | 88 | |
| 5453 | 20 | Bone marrow | 91 | |

| | | | |
|---|---|---|---|
| | | Adipose tissue | 86 |
| | | Heart | 74 |
| | | Liver | 93 |
| 3868 | 21 | Heart | 82 |
| | | Kidney | 89 |
| | | Large intestine | 91 |
| | | Pancreas | 80 |
| | | Small intestine | 83 |
| | | Spleen | 89 |
| | | Stomach | 83 |
| 5311 | 22 | Adipose tissue | 89 |
| | | Liver | 85 |
| | | Spleen | 85 |
| 5593 | 25 | Small intestine | 89 |
| | | Liver | 92 |
| | | Large intestine | 88 |
| | | Spleen | 89 |
| | | Kidney | 61 |
| 5638 | 27 | Spleen | 98 |
| 5433 | 31 | Liver | 82 |
| | | Stomach | 94 |
| | | Kidney | 79 |
| | | Spleen | 95 |
| | | Large Intestine | 85 |
| 5532 | 31 | Liver | 96 |
| | | Pancreas | 64 |
| | | Stomach | 90 |

| | | |
|---|---|---|
| | Small Intestine | 91 |
| | Kidney | 70 |
| | Large intestine | 90 |

**Supplementary Table 2. Serum acylcarnitine levels: 5 month WT and MlxKO mice**

| | P value | Mean of WT 5mos | Mean of MlxKO 5 mos | Difference | SE of difference | t ratio |
|---|---|---|---|---|---|---|
| C0 | 0.548594 | 34.71 | 38.75 | -4.043 | 6.416 | 0.6302 |
| C2 | 0.549941 | 54.89 | 48.82 | 6.064 | 9.655 | 0.628 |
| C3 | 0.933289 | 1.936 | 1.984 | -0.04755 | 0.548 | 0.08676 |
| C4 | 0.427705 | 1.823 | 1.375 | 0.4478 | 0.5319 | 0.8418 |
| C5 | 0.949985 | 0.2986 | 0.2933 | 0.00535 | 0.0823 | 0.06501 |
| C6 | 0.845501 | 0.2048 | 0.2145 | -0.0097 | 0.04797 | 0.2022 |
| C7 | 0.660826 | 0.1328 | 0.1428 | -0.00995 | 0.02172 | 0.458 |
| C8 | 0.833693 | 0.113 | 0.1183 | -0.00525 | 0.02409 | 0.2179 |
| C9 | 0.281512 | 0.0712 | 0.08725 | -0.01605 | 0.01376 | 1.167 |
| C10 | 0.064004 | 0.1304 | 0.2163 | -0.08585 | 0.03907 | 2.197 |
| C12 | 0.637483 | 0.1066 | 0.1188 | -0.01215 | 0.02467 | 0.4924 |
| C14 | 0.276868 | 0.2424 | 0.298 | -0.0556 | 0.04715 | 1.179 |
| C16 | 0.74747 | 0.652 | 0.6203 | 0.03175 | 0.09479 | 0.335 |
| C18 | 0.699576 | 0.2104 | 0.1938 | 0.01665 | 0.0414 | 0.4021 |
| C8:1 | 0.289567 | 0.1066 | 0.1888 | -0.08215 | 0.0717 | 1.146 |
| C10:1 | 0.236804 | 0.1496 | 0.2053 | -0.05565 | 0.04301 | 1.294 |
| C3:1 | 0.393799 | 0.083 | 0.1118 | -0.02875 | 0.03164 | 0.9085 |
| Crotonyl / Me | 0.435421 | 0.1394 | 0.2008 | -0.06135 | 0.07417 | 0.8272 |
| Figlu | 0.995623 | 0.1574 | 0.1573 | 0.00015 | 0.02639 | 0.005685 |
| C5:1 | 0.342821 | 0.0608 | 0.08175 | -0.02095 | 0.02059 | 1.017 |
| C3-OH | 0.435213 | 0.3656 | 0.3328 | 0.03285 | 0.03969 | 0.8276 |
| C4-OH | 0.311007 | 0.7836 | 0.8593 | -0.07565 | 0.06928 | 1.092 |
| C5-OH | 0.441613 | 0.1724 | 0.1898 | -0.01735 | 0.02127 | 0.8156 |
| C6-OH | 0.262673 | 0.0924 | 0.117 | -0.0246 | 0.0202 | 1.218 |
| C2DC | 0.492812 | 0.0632 | 0.07675 | -0.01355 | 0.01873 | 0.7235 |
| C8-OH | 0.606769 | 0.1462 | 0.154 | -0.0078 | 0.01448 | 0.5387 |
| C10:2 | 0.352861 | 0.0828 | 0.1625 | -0.0797 | 0.0801 | 0.995 |
| C4DC | 0.342363 | 0.1168 | 0.156 | -0.0392 | 0.03849 | 1.018 |
| C10:1-OH | 0.599678 | 0.1106 | 0.0965 | 0.0141 | 0.02565 | 0.5496 |
| C10-OH (C5D | 0.699714 | 0.1208 | 0.1328 | -0.01195 | 0.02973 | 0.4019 |
| C12:1 | 0.864531 | 0.0876 | 0.09125 | -0.00365 | 0.02062 | 0.177 |
| C6DC | 0.690628 | 0.426 | 0.39 | 0.036 | 0.08677 | 0.4149 |
| C12:1-OH | 0.264378 | 0.0722 | 0.149 | -0.0768 | 0.0633 | 1.213 |
| C12-OH | 0.545436 | 0.1208 | 0.099 | 0.0218 | 0.03431 | 0.6353 |
| C14:2 | 0.00927 | 0.2566 | 0.1153 | 0.1414 | 0.03975 | 3.556 |
| C14:1 | 0.798863 | 0.1952 | 0.1865 | 0.0087 | 0.03287 | 0.2647 |
| C8DC | 0.184811 | 0.1434 | 0.2098 | -0.06635 | 0.04511 | 1.471 |
| C14:1-OH | 0.082436 | 0.0806 | 0.1268 | -0.04615 | 0.02278 | 2.026 |
| C14-OH | 0.578705 | 0.0514 | 0.05975 | -0.00835 | 0.01434 | 0.5822 |
| C16:2 | 0.559678 | 0.12 | 0.131 | -0.011 | 0.01796 | 0.6123 |
| C16:1 | 0.89597 | 0.229 | 0.2345 | -0.0055 | 0.04057 | 0.1356 |
| C10DC | 0.172962 | 0.072 | 0.09425 | -0.02225 | 0.01466 | 1.517 |
| C16-OH | 0.187956 | 0.0854 | 0.1083 | -0.02285 | 0.01566 | 1.459 |
| C16:1-OH | 0.187956 | 0.0854 | 0.1083 | -0.02285 | 0.01566 | 1.459 |
| C18:3 | 0.012975 | 0.1778 | 0.055 | 0.1228 | 0.03712 | 3.308 |
| C18:2 | 0.813765 | 0.26 | 0.2745 | -0.0145 | 0.05928 | 0.2446 |
| C18:1 | 0.392628 | 0.6884 | 0.536 | 0.1524 | 0.1673 | 0.9109 |
| C12DC | 0.057416 | 0.0538 | 0.078 | -0.0242 | 0.01066 | 2.271 |
| C18:2-OH | 0.635748 | 0.078 | 0.06825 | 0.00975 | 0.0197 | 0.495 |
| C18:1-OH | 0.331346 | 0.0978 | 0.0785 | 0.0193 | 0.01849 | 1.044 |
| C18-OH | 0.41041 | 0.063 | 0.07775 | -0.01475 | 0.01685 | 0.8754 |

**Supplementary Table 3. Serum acylcarnitine levels: 20 month WT and *Mlx* KO mice**

| | P value | Mean of WT 20mos | Mean of MlxKO 20mos | Difference | SE of difference | t ratio |
|---|---|---|---|---|---|---|
| C0 | 0.625899 | 43.63 | 41.14 | 2.489 | 4.931 | 0.5047 |
| C2 | 0.888484 | 53.57 | 52.7 | 0.8707 | 6.036 | 0.1442 |
| C3 | 0.796305 | 1.505 | 1.423 | 0.0819 | 0.308 | 0.2659 |
| C4 | 0.074544 | 2.666 | 1.615 | 1.05 | 0.5208 | 2.016 |
| C5 | 0.869218 | 0.1652 | 0.171 | -0.005833 | 0.03443 | 0.1694 |
| C6 | 0.050208 | 0.338 | 0.2328 | 0.1052 | 0.04656 | 2.26 |
| C7 | 0.994275 | 0.1333 | 0.1334 | -6.67E-05 | 0.009037 | 0.007377 |
| C8 | 0.718718 | 0.1138 | 0.1174 | -0.003567 | 0.009596 | 0.3717 |
| C9 | 0.528966 | 0.116 | 0.1216 | -0.0056 | 0.008552 | 0.6548 |
| C10 | 0.798725 | 0.1375 | 0.144 | -0.0065 | 0.02475 | 0.2627 |
| C12 | 0.956378 | 0.1052 | 0.106 | -0.0008333 | 0.01482 | 0.05624 |
| C14 | 0.565468 | 0.1833 | 0.203 | -0.01967 | 0.03296 | 0.5966 |
| C16 | 0.852866 | 0.4013 | 0.389 | 0.01233 | 0.06462 | 0.1909 |
| C18 | 0.135671 | 0.1325 | 0.1094 | 0.0231 | 0.0141 | 1.639 |
| C8:1 | 0.358068 | 0.5018 | 0.3408 | 0.161 | 0.1663 | 0.9686 |
| C10:1 | 0.261019 | 0.3303 | 0.223 | 0.1073 | 0.08949 | 1.199 |
| C3:1 | 0.228638 | 0.2602 | 0.1662 | 0.09397 | 0.07275 | 1.292 |
| Crotonyl / Me-Acrylyl | 0.37111 | 0.5775 | 0.3714 | 0.2061 | 0.2189 | 0.9413 |
| Figlu | 0.349148 | 0.321 | 0.2072 | 0.1138 | 0.1152 | 0.9876 |
| C5:1 | 0.045734 | 0.042 | 0.0538 | -0.0118 | 0.005093 | 2.317 |
| C3-OH | 0.109183 | 0.2785 | 0.2094 | 0.0691 | 0.03887 | 1.778 |
| C4-OH | 0.188395 | 1.662 | 1.146 | 0.5164 | 0.3628 | 1.423 |
| C5-OH | 0.289939 | 0.1005 | 0.1184 | -0.0179 | 0.01592 | 1.124 |
| C6-OH | 0.96035 | 0.1013 | 0.1022 | -0.0008667 | 0.01696 | 0.05112 |
| C2DC | 0.94062 | 0.193 | 0.1908 | 0.0022 | 0.02872 | 0.0766 |
| C8-OH | 0.292902 | 0.3257 | 0.1532 | 0.1725 | 0.1544 | 1.117 |
| C10:2 | 0.16925 | 0.2447 | 0.1532 | 0.09147 | 0.0612 | 1.495 |
| C4DC | 0.148943 | 0.3838 | 0.2674 | 0.1164 | 0.07377 | 1.578 |
| C10:1-OH | 0.405108 | 0.1365 | 0.1086 | 0.0279 | 0.03194 | 0.8735 |
| C10-OH (C5DC) | 0.469388 | 0.1117 | 0.1222 | -0.01053 | 0.01395 | 0.7553 |
| C12:1 | 0.846614 | 0.08933 | 0.0874 | 0.001933 | 0.009711 | 0.1991 |
| C6DC | 0.921003 | 0.2507 | 0.2538 | -0.003133 | 0.03072 | 0.102 |
| C12:1-OH | 0.828888 | 0.063 | 0.0646 | -0.0016 | 0.007191 | 0.2225 |
| C12-OH | 0.182724 | 0.07483 | 0.0622 | 0.01263 | 0.008751 | 1.444 |
| C14:2 | 0.475765 | 0.1437 | 0.136 | 0.007667 | 0.0103 | 0.7441 |
| C14:1 | 0.492798 | 0.2193 | 0.2028 | 0.01653 | 0.02313 | 0.7149 |
| C8DC | 0.294664 | 0.06017 | 0.0798 | -0.01963 | 0.01764 | 1.113 |
| C14:1-OH | 0.854228 | 0.08083 | 0.0832 | -0.002367 | 0.01252 | 0.1891 |
| C14-OH | 0.343857 | 0.07433 | 0.0874 | -0.01307 | 0.01308 | 0.9991 |
| C16:2 | 0.373093 | 0.07133 | 0.0868 | -0.01547 | 0.0165 | 0.9373 |
| C16:1 | 0.831389 | 0.2312 | 0.238 | -0.006833 | 0.03117 | 0.2192 |
| C10DC | 0.915181 | 0.4188 | 0.4158 | 0.003033 | 0.02769 | 0.1095 |
| C16-OH | 0.099289 | 0.1192 | 0.0804 | 0.03877 | 0.0211 | 1.838 |
| C16:1-OH | 0.126941 | 0.08233 | 0.0958 | -0.01347 | 0.008008 | 1.682 |
| C18:3 | 0.414354 | 0.058 | 0.0524 | 0.0056 | 0.006544 | 0.8557 |
| C18:2 | 0.194067 | 0.1787 | 0.2058 | -0.02713 | 0.01934 | 1.403 |
| C18:1 | 0.288622 | 0.526 | 0.4694 | 0.0566 | 0.05019 | 1.128 |
| C12DC | 0.710472 | 0.3005 | 0.3076 | -0.0071 | 0.01853 | 0.3832 |
| C18:2-OH | 0.629184 | 0.06733 | 0.0716 | -0.004267 | 0.008536 | 0.4998 |
| C18:1-OH | 0.560507 | 0.08667 | 0.0792 | 0.007467 | 0.01235 | 0.6044 |
| C18-OH | 0.002285 | 0.0645 | 0.0388 | 0.0257 | 0.00611 | 4.206 |

**Supplementary Table 4. Serum acylcarnitine levels: 5 month vs. 20 month WT mice**

| | P value | Mean of WT 20mos | Mean of WT 5mos | Difference | SE of difference | t ratio |
|---|---|---|---|---|---|---|
| C0 | 0.137324 | 43.63 | 34.71 | 8.923 | 5.471 | 1.631 |
| C2 | 0.867123 | 53.57 | 54.89 | -1.316 | 7.645 | 0.1722 |
| C3 | 0.338557 | 1.505 | 1.936 | -0.4317 | 0.4271 | 1.011 |
| C4 | 0.198952 | 2.666 | 1.823 | 0.8429 | 0.6079 | 1.387 |
| C5 | 0.054415 | 0.1652 | 0.2986 | -0.1334 | 0.06037 | 2.21 |
| C6 | 0.045689 | 0.338 | 0.2048 | 0.1332 | 0.05748 | 2.317 |
| C7 | 0.963978 | 0.1333 | 0.1328 | 0.0005333 | 0.01149 | 0.04643 |
| C8 | 0.962316 | 0.1138 | 0.113 | 0.0008333 | 0.01715 | 0.04858 |
| C9 | 0.000936 | 0.116 | 0.0712 | 0.0448 | 0.009278 | 4.829 |
| C10 | 0.696317 | 0.1375 | 0.1304 | 0.0071 | 0.01762 | 0.4031 |
| C12 | 0.940283 | 0.1052 | 0.1066 | -0.001433 | 0.01861 | 0.07703 |
| C14 | 0.134662 | 0.1833 | 0.2424 | -0.05907 | 0.03594 | 1.644 |
| C16 | 0.006241 | 0.4013 | 0.652 | -0.2507 | 0.07066 | 3.547 |
| C18 | 0.041676 | 0.1325 | 0.2104 | -0.0779 | 0.03282 | 2.373 |
| C8:1 | 0.040676 | 0.5018 | 0.1066 | 0.3952 | 0.1655 | 2.388 |
| C10:1 | 0.076358 | 0.3303 | 0.1496 | 0.1807 | 0.0903 | 2.002 |
| C3:1 | 0.039652 | 0.2602 | 0.083 | 0.1772 | 0.0737 | 2.404 |
| Crotonyl / Me-Acrylyl | 0.071318 | 0.5775 | 0.1394 | 0.4381 | 0.2143 | 2.044 |
| Figlu | 0.187693 | 0.321 | 0.1574 | 0.1636 | 0.1147 | 1.426 |
| C5:1 | 0.022373 | 0.042 | 0.0608 | -0.0188 | 0.006829 | 2.753 |
| C3-OH | 0.093491 | 0.2785 | 0.3656 | -0.0871 | 0.04644 | 1.875 |
| C4-OH | 0.023059 | 1.662 | 0.7836 | 0.8786 | 0.3213 | 2.734 |
| C5-OH | 0.00117 | 0.1005 | 0.1724 | -0.0719 | 0.0154 | 4.669 |
| C6-OH | 0.614363 | 0.1013 | 0.0924 | 0.008933 | 0.01712 | 0.5219 |
| C2DC | 0.00004 | 0.193 | 0.0632 | 0.1298 | 0.01746 | 7.433 |
| C8-OH | 0.275325 | 0.3257 | 0.1462 | 0.1795 | 0.1545 | 1.161 |
| C10:2 | 0.02499 | 0.2447 | 0.0828 | 0.1619 | 0.06028 | 2.685 |
| C4DC | 0.001343 | 0.3838 | 0.1168 | 0.267 | 0.05841 | 4.572 |
| C10:1-OH | 0.452006 | 0.1365 | 0.1106 | 0.0259 | 0.03295 | 0.7861 |
| C10-OH (C5DC) | 0.675613 | 0.1117 | 0.1208 | -0.009133 | 0.02112 | 0.4324 |
| C12:1 | 0.846046 | 0.08933 | 0.0876 | 0.001733 | 0.008673 | 0.1998 |
| C6DC | 0.002847 | 0.2507 | 0.426 | -0.1753 | 0.0432 | 4.059 |
| C12:1-OH | 0.12923 | 0.063 | 0.0722 | -0.0092 | 0.005509 | 1.67 |
| C12-OH | 0.097276 | 0.07483 | 0.1208 | -0.04597 | 0.02484 | 1.85 |
| C14:2 | 0.007265 | 0.1437 | 0.2566 | -0.1129 | 0.03273 | 3.451 |
| C14:1 | 0.270265 | 0.2193 | 0.1952 | 0.02413 | 0.02054 | 1.175 |
| C8DC | 0.008132 | 0.06017 | 0.1434 | -0.08323 | 0.02463 | 3.38 |
| C14:1-OH | 0.98218 | 0.08083 | 0.0806 | 0.0002333 | 0.01016 | 0.02296 |
| C14-OH | 0.066759 | 0.07433 | 0.0514 | 0.02293 | 0.011 | 2.085 |
| C16:2 | 0.002912 | 0.07133 | 0.12 | -0.04867 | 0.01204 | 4.044 |
| C16:1 | 0.95346 | 0.2312 | 0.229 | 0.002167 | 0.03611 | 0.06001 |
| C10DC | <0.000001 | 0.4188 | 0.072 | 0.3468 | 0.01715 | 20.22 |
| C16-OH | 0.169558 | 0.1192 | 0.0854 | 0.03377 | 0.02261 | 1.493 |
| C16:1-OH | 0.791838 | 0.08233 | 0.0854 | -0.003067 | 0.01128 | 0.2719 |
| C18:3 | 0.00276 | 0.058 | 0.1778 | -0.1198 | 0.02937 | 4.08 |
| C18:2 | 0.021146 | 0.1787 | 0.26 | -0.08133 | 0.02918 | 2.787 |
| C18:1 | 0.211098 | 0.526 | 0.6884 | -0.1624 | 0.1206 | 1.346 |
| C12DC | <0.000001 | 0.3005 | 0.0538 | 0.2467 | 0.01729 | 14.27 |
| C18:2-OH | 0.442574 | 0.06733 | 0.078 | -0.01067 | 0.01328 | 0.8032 |
| C18:1-OH | 0.503325 | 0.08667 | 0.0978 | -0.01113 | 0.01597 | 0.6971 |
| C18-OH | 0.859542 | 0.0645 | 0.063 | 0.0015 | 0.008237 | 0.1821 |

**Supplementary Table 5. Serum acylcarnitine levels: 5 month vs. 20 month *Mlx* KO mice**

| | P value | Mean of MlxKO 20mos | Mean of MlxKO 5 mos | Difference | SE of difference | t ratio |
|---|---|---|---|---|---|---|
| C0 | 0.686521 | 41.14 | 38.75 | 2.392 | 5.683 | 0.4208 |
| C2 | 0.627807 | 52.7 | 48.82 | 3.877 | 7.648 | 0.5069 |
| C3 | 0.206123 | 1.423 | 1.984 | -0.5612 | 0.4027 | 1.393 |
| C4 | 0.523088 | 1.615 | 1.375 | 0.2404 | 0.3577 | 0.6721 |
| C5 | 0.05724 | 0.171 | 0.2933 | -0.1223 | 0.05379 | 2.273 |
| C6 | 0.443384 | 0.2328 | 0.2145 | 0.0183 | 0.02253 | 0.8123 |
| C7 | 0.651386 | 0.1334 | 0.1428 | -0.00935 | 0.01982 | 0.4719 |
| C8 | 0.959529 | 0.1174 | 0.1183 | -0.00085 | 0.01616 | 0.05259 |
| C9 | 0.03324 | 0.1216 | 0.08725 | 0.03435 | 0.01299 | 2.644 |
| C10 | 0.150525 | 0.144 | 0.2163 | -0.07225 | 0.04476 | 1.614 |
| C12 | 0.548378 | 0.106 | 0.1188 | -0.01275 | 0.02022 | 0.6305 |
| C14 | 0.06571 | 0.203 | 0.298 | -0.095 | 0.04359 | 2.179 |
| C16 | 0.033604 | 0.389 | 0.6203 | -0.2313 | 0.08772 | 2.636 |
| C18 | 0.002319 | 0.1094 | 0.1938 | -0.08435 | 0.01811 | 4.658 |
| C8:1 | 0.080433 | 0.3408 | 0.1888 | 0.1521 | 0.07445 | 2.042 |
| C10:1 | 0.672792 | 0.223 | 0.2053 | 0.01775 | 0.04029 | 0.4406 |
| C3:1 | 0.092199 | 0.1662 | 0.1118 | 0.05445 | 0.02793 | 1.95 |
| Crotonyl / Me-Acrylyl | 0.10904 | 0.3714 | 0.2008 | 0.1707 | 0.09296 | 1.836 |
| Figlu | 0.134249 | 0.2072 | 0.1573 | 0.04995 | 0.0295 | 1.693 |
| C5:1 | 0.200578 | 0.0538 | 0.08175 | -0.02795 | 0.01978 | 1.413 |
| C3-OH | 0.001213 | 0.2094 | 0.3328 | -0.1234 | 0.02359 | 5.23 |
| C4-OH | 0.23915 | 1.146 | 0.8593 | 0.2866 | 0.2227 | 1.287 |
| C5-OH | 0.013816 | 0.1184 | 0.1898 | -0.07135 | 0.02187 | 3.263 |
| C6-OH | 0.482919 | 0.1022 | 0.117 | -0.0148 | 0.01998 | 0.7408 |
| C2DC | 0.012548 | 0.1908 | 0.07675 | 0.1141 | 0.03422 | 3.332 |
| C8-OH | 0.94974 | 0.1532 | 0.154 | -0.0008 | 0.01225 | 0.06533 |
| C10:2 | 0.912024 | 0.1532 | 0.1625 | -0.0093 | 0.08119 | 0.1145 |
| C4DC | 0.147661 | 0.2674 | 0.156 | 0.1114 | 0.06845 | 1.627 |
| C10:1-OH | 0.623387 | 0.1086 | 0.0965 | 0.0121 | 0.02356 | 0.5135 |
| C10-OH (C5DC) | 0.647129 | 0.1222 | 0.1328 | -0.01055 | 0.02207 | 0.4781 |
| C12:1 | 0.861941 | 0.0874 | 0.09125 | -0.00385 | 0.02134 | 0.1804 |
| C6DC | 0.124009 | 0.2538 | 0.39 | -0.1362 | 0.07793 | 1.748 |
| C12:1-OH | 0.225922 | 0.0646 | 0.149 | -0.0844 | 0.06357 | 1.328 |
| C12-OH | 0.080444 | 0.0622 | 0.099 | -0.0368 | 0.01802 | 2.042 |
| C14:2 | 0.029216 | 0.136 | 0.1153 | 0.02075 | 0.007593 | 2.733 |
| C14:1 | 0.659784 | 0.2028 | 0.1865 | 0.0163 | 0.03547 | 0.4595 |
| C8DC | 0.013478 | 0.0798 | 0.2098 | -0.13 | 0.03961 | 3.28 |
| C14:1-OH | 0.119517 | 0.0832 | 0.1268 | -0.04355 | 0.02456 | 1.773 |
| C14-OH | 0.145256 | 0.0874 | 0.05975 | 0.02765 | 0.01687 | 1.639 |
| C16:2 | 0.09476 | 0.0868 | 0.131 | -0.0442 | 0.02289 | 1.931 |
| C16:1 | 0.919716 | 0.238 | 0.2345 | 0.0035 | 0.0335 | 0.1045 |
| C10DC | 0.000017 | 0.4158 | 0.09425 | 0.3216 | 0.031 | 10.37 |
| C16-OH | 0.051298 | 0.0804 | 0.1083 | -0.02785 | 0.01187 | 2.347 |
| C16:1-OH | 0.336763 | 0.0958 | 0.1083 | -0.01245 | 0.01207 | 1.031 |
| C18:3 | 0.785854 | 0.0524 | 0.055 | -0.0026 | 0.009209 | 0.2823 |
| C18:2 | 0.232281 | 0.2058 | 0.2745 | -0.0687 | 0.05253 | 1.308 |
| C18:1 | 0.505545 | 0.4694 | 0.536 | -0.0666 | 0.09492 | 0.7017 |
| C12DC | <0.000001 | 0.3076 | 0.078 | 0.2296 | 0.01355 | 16.95 |
| C18:2-OH | 0.829509 | 0.0716 | 0.06825 | 0.00335 | 0.01499 | 0.2235 |
| C18:1-OH | 0.959889 | 0.0792 | 0.0785 | 0.0007 | 0.01343 | 0.05212 |
| C18-OH | 0.038853 | 0.0388 | 0.07775 | -0.03895 | 0.01535 | 2.537 |

**Supplementary Table 6. Antibodies used in the current study.**

| Name of Antibody | Vendor | Catalog no. | Dilution used |
|---|---|---|---|
| GAPDH | Sigma | G8795 | 1:10000 |
| GLUT1 | Abcam | AB115730 | 1:2000 |
| GLUT2 | ProteinTech | 20436-1-AP | 1:300 |
| GLUT4 | Cell Signaling Technology | 2213 | 1:1000 |
| PDH | Cell Signaling Technology | 3205 | 1:1000 |
| PFKL | Aviva System Biology | ARP45774 | 1:1000 |
| PFKM | R&D Systems | MAB7687 | 1:3000 |
| PGC1 | Novus Biological | NBP1-04676 | 1:1000 |
| pPDH-E1α(pSer293) | Millipore (Calbiochem) | AP1062 | 1:1000 |

A

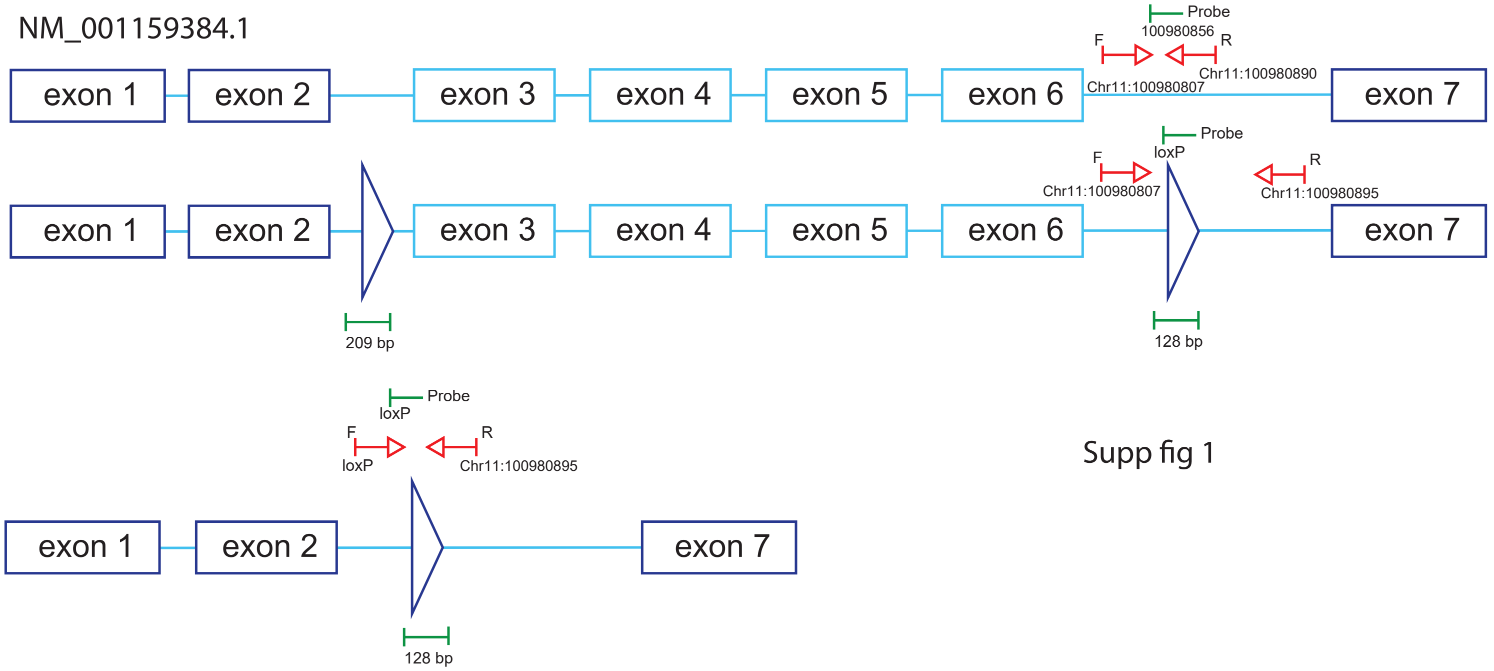

Supp fig 1

D

| | | Start bp | End bp | Sequence(5'-3') |
|---|---|---|---|---|
| MLX EXON 1 (NC_000077.7) | Fwd | 100978116 | 100978132 | AGTCCGCTGGCTTGTTT |
| | Rev | 100978192 | 100978176 | TTGACCCAAGGGTCCTC |
| | Probe | 100978134 | 100978157 | /56-FAM/CGGTTCGGT/ZEN/AGGTTCACGATGACG/3IABkFQ/ |
| MLX EXON 7 (NC_000077.7) | Fwd | 100980596 | 100980617 | CTCCCTCTTCCAGTCTTTCAAC |
| | Rev | 100980691 | 100980672 | GGCTCACAGTGTTCCTCAAT |
| | Probe | 100980655 | 100980669 | /56-FAM/AGCTCTCAG/ZEN/CTTGTGTCTTCAGCT/3IABkFQ/ |
| WT (NC_000077.7) | Fwd | 100980807 | 100980825 | TAGCCCAGTGAAGGTCTCA |
| | Rev | 100980911 | 100980890 | AGGAGTAGACAGGGTAGCTAAT |
| | Probe | 100980856 | 100980879 | /56-FAM/CAGGTCCAGCTTTAGCCCATGTCA/3IABkFQ/ |
| mlx $^{loxP/loxP}$ (NC_000077.7) | Fwd | 100980807 | 100980825 | TAGCCCAGTGAAGGTCTCA |
| | Rev | 100980914 | 100980895 | CCAAGGAGTAGACAGGGTAGAA |
| | Probe | loxP | loxP | /56-FAM/ACTAGTTCTAGAGCGGCCAATTCGC/3IABkFQ/ |
| mlx -/ - (KO) (NC_000077.7) | Fwd | loxP | loxP | CAGGTCGAGGGACCTAATAACT |
| | Rev | 100980914 | 100980895 | CCAAGGAGTAGACAGGGTAGAA |
| | Probe | loxP | loxP | /56-FAM/ACTAGTTCTAGAGCGGCCAATTCGC/3IABkFQ/ |

E

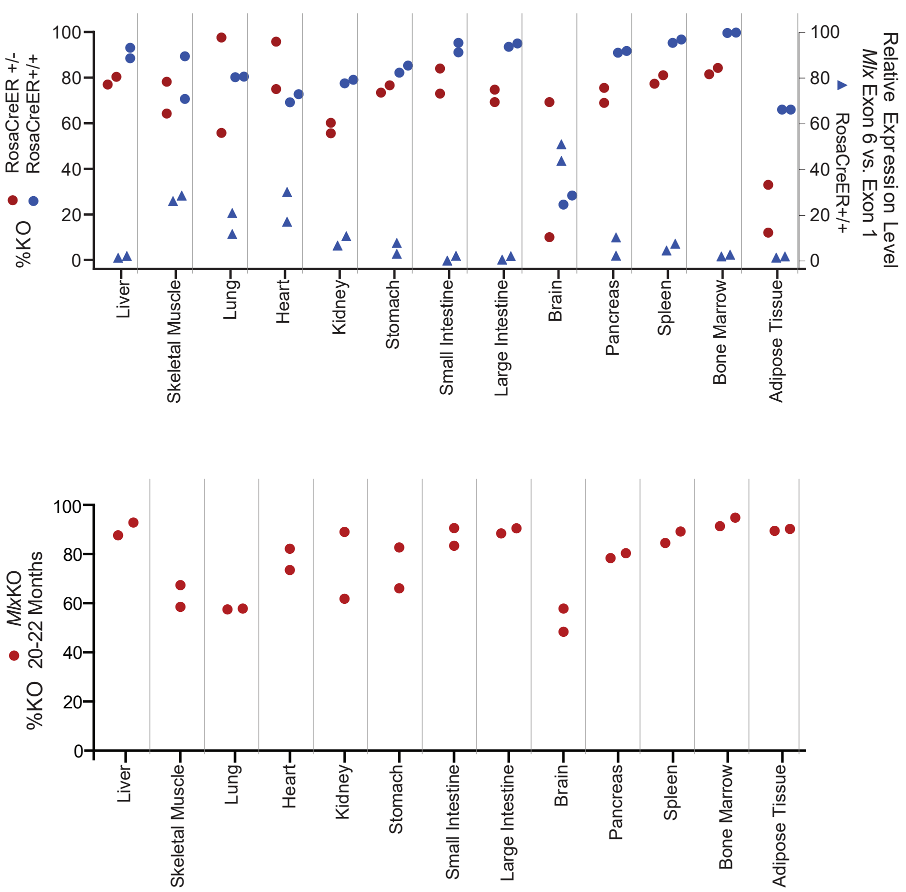

F

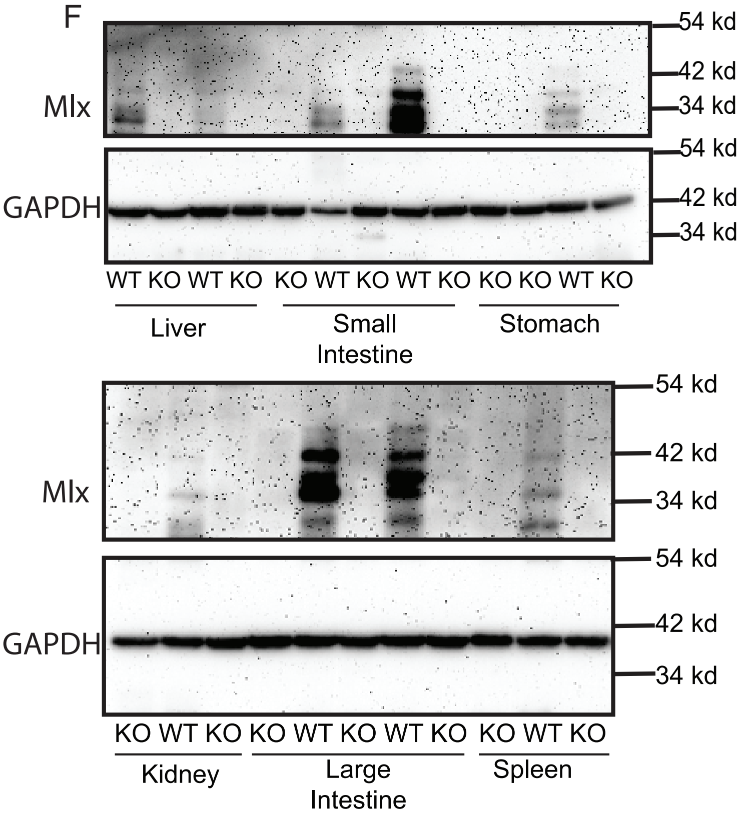

**Supplementary Figure 1. Strategy for Taq-Man-based qPCR and qRT-PCR assays for quantifying "floxed" (WT) and excised *Mlx* loci.**
**(A)** The WT *Mlx* locus and the unexcised $Mlx^{loxP/loxP}$ locus.
**(B)** LoxP sites flanking exons 3 and 6 are indicated by the red triangles. Red arrows in A indicate the primer set used for identifying WT alleles. The TaqMan probe is shown in green. Numbering of primers and probes is based on their correspondence to the sequence of the Mlx genomic locus (NC_000077.7 (100977538..100983033)). Black arrows indicate the primer set used to uniquely identify $Mlx^{loxP/loxP}$ alleles. The unique TaqMan probe used to quantify the PCR products is depicted in green.
**(C)** The *Mlx*KO locus following Cre-recombinase mediated recombination. Blue arrows indicate primers used to amplify KO (*Mlx*-/-) alleles.
**(D)** Sequences, genomic locations, and dyes for primers and probes shown in **A**-**C** and for quantification of WT transcripts by qRT-PCR.
**(E)** Excisional efficiency of the *Mlx* locus in tissues of randomly chosen *Mlx*KO mice 2 months after tamoxifen treatment. The indicated tissues were sampled from mice bearing either one or 2 copies of the CreER transgene. qRT-PCR results in each *Mlx*KO tissue were compared to the results obtained in the identical tissues from WT mice.
**(F)** Absence of Mlx protein in *Mlx*KO tissues. Total cell lysates from the indicated, age-matched WT and *Myc*KO tissues were examined by immuno-blotting for the expression of *Mlx*. GAPDH was used as a loading control.

Supp. fig 2

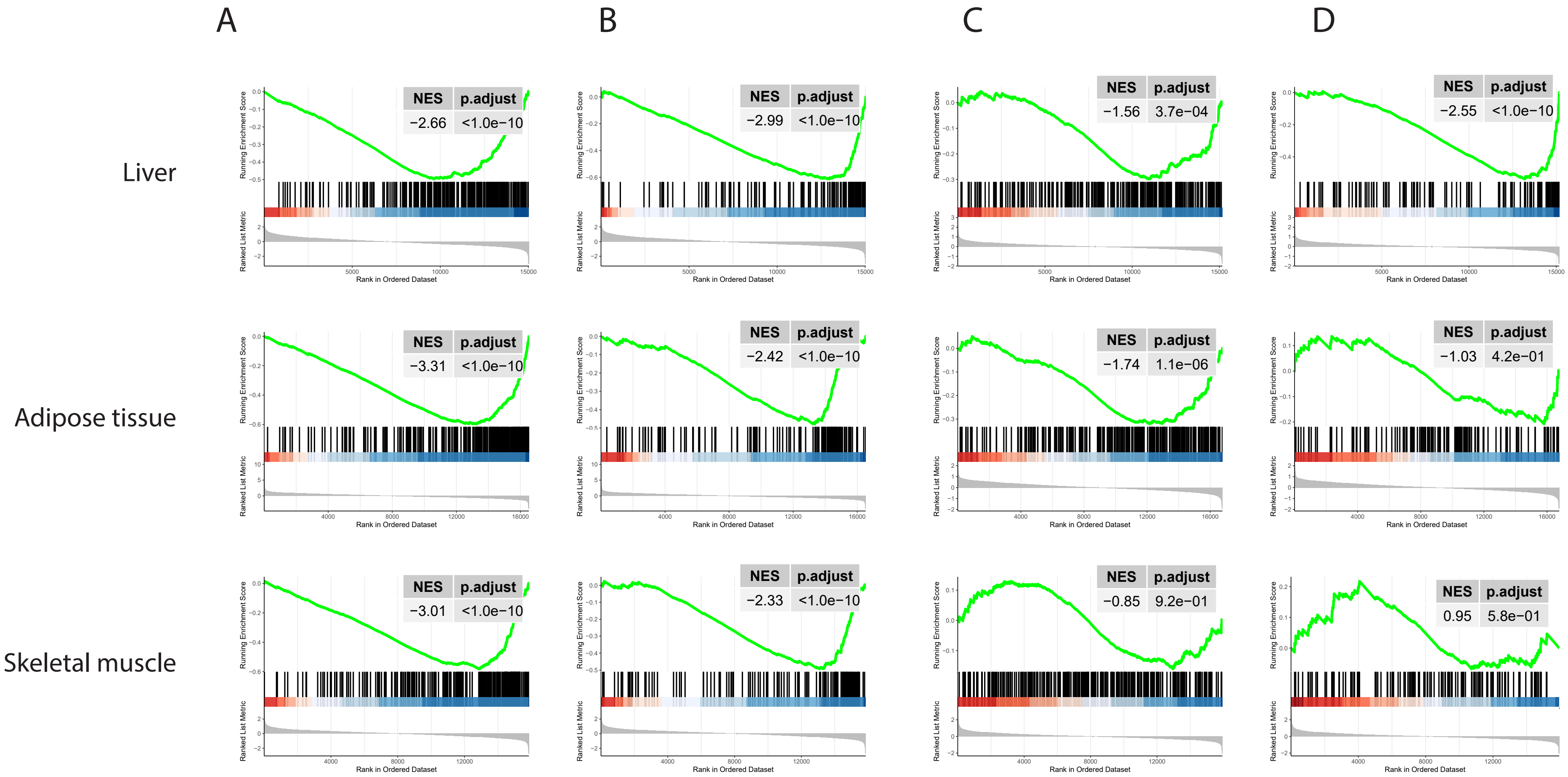
A
B
C
D
Liver
Adipose tissue
Skeletal muscle
NES p.adjust
−2.66 <1.0e−10
NES p.adjust
−2.99 <1.0e−10
NES p.adjust
−1.56 3.7e−04
NES p.adjust
−2.55 <1.0e−10
NES p.adjust
−3.31 <1.0e−10
NES p.adjust
−2.42 <1.0e−10
NES p.adjust
−1.74 1.1e−06
NES p.adjust
−1.03 4.2e−01
NES p.adjust
−3.01 <1.0e−10
NES p.adjust
−2.33 <1.0e−10
NES p.adjust
−0.85 9.2e−01
NES p.adjust
0.95 5.8e−01
Running Enrichment Score
Ranked List Metric
Rank in Ordered Dataset

**Supplementary Figure 2. GSEA with validated ChREBP, MondoA and/or Mlx target genes**

(A). GSEA of 264 ChREBP/MondoA/Mlx target genes from the Enrichr datbase gene set ARCHS4 in the indicated tissues of 5 month old *Mlx*KO versus WT mice.

(B). GSEA of a 181 member ChREBP/MondoA/Mlx direct target gene set from the Qiagen database in the indicated tissues of 5 month old *Mlx*KO versus WT mice.

(C). GSEA of the gene set from **A** performed on tissues from 5 month old *Myc*KO mice.

(D). GSEA of the gene set from **B** performed on tissues from 5 month old *Myc*KO mice.

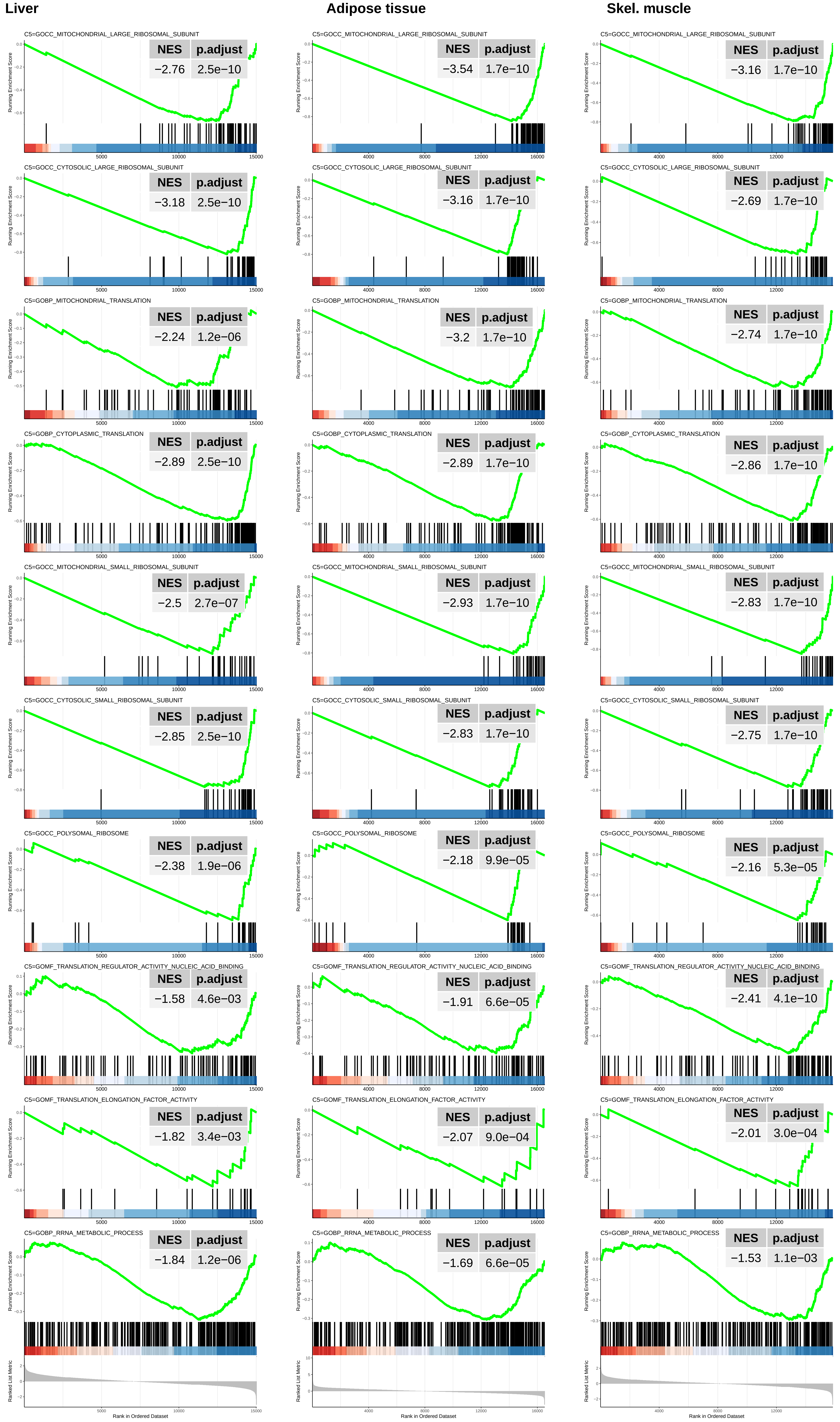

Liver
Adipose tissue
Skel. muscle
C5=GOCC_MITOCHONDRIAL_LARGE_RIBOSOMAL_SUBUNIT
NES −2.76 p.adjust 2.5e−10
C5=GOCC_MITOCHONDRIAL_LARGE_RIBOSOMAL_SUBUNIT
NES −3.54 p.adjust 1.7e−10
C5=GOCC_MITOCHONDRIAL_LARGE_RIBOSOMAL_SUBUNIT
NES −3.16 p.adjust 1.7e−10
C5=GOCC_CYTOSOLIC_LARGE_RIBOSOMAL_SUBUNIT
NES −3.18 p.adjust 2.5e−10
C5=GOCC_CYTOSOLIC_LARGE_RIBOSOMAL_SUBUNIT
NES −3.16 p.adjust 1.7e−10
C5=GOCC_CYTOSOLIC_LARGE_RIBOSOMAL_SUBUNIT
NES −2.69 p.adjust 1.7e−10
C5=GOBP_MITOCHONDRIAL_TRANSLATION
NES −2.24 p.adjust 1.2e−06
C5=GOBP_MITOCHONDRIAL_TRANSLATION
NES −3.2 p.adjust 1.7e−10
C5=GOBP_MITOCHONDRIAL_TRANSLATION
NES −2.74 p.adjust 1.7e−10
C5=GOBP_CYTOPLASMIC_TRANSLATION
NES −2.89 p.adjust 2.5e−10
C5=GOBP_CYTOPLASMIC_TRANSLATION
NES −2.89 p.adjust 1.7e−10
C5=GOBP_CYTOPLASMIC_TRANSLATION
NES −2.86 p.adjust 1.7e−10
C5=GOCC_MITOCHONDRIAL_SMALL_RIBOSOMAL_SUBUNIT
NES −2.5 p.adjust 2.7e−07
C5=GOCC_MITOCHONDRIAL_SMALL_RIBOSOMAL_SUBUNIT
NES −2.93 p.adjust 1.7e−10
C5=GOCC_MITOCHONDRIAL_SMALL_RIBOSOMAL_SUBUNIT
NES −2.83 p.adjust 1.7e−10
C5=GOCC_CYTOSOLIC_SMALL_RIBOSOMAL_SUBUNIT
NES −2.85 p.adjust 2.5e−10
C5=GOCC_CYTOSOLIC_SMALL_RIBOSOMAL_SUBUNIT
NES −2.83 p.adjust 1.7e−10
C5=GOCC_CYTOSOLIC_SMALL_RIBOSOMAL_SUBUNIT
NES −2.75 p.adjust 1.7e−10
C5=GOCC_POLYSOMAL_RIBOSOME
NES −2.38 p.adjust 1.9e−06
C5=GOCC_POLYSOMAL_RIBOSOME
NES −2.18 p.adjust 9.9e−05
C5=GOCC_POLYSOMAL_RIBOSOME
NES −2.16 p.adjust 5.3e−05
C5=GOMF_TRANSLATION_REGULATOR_ACTIVITY_NUCLEIC_ACID_BINDING
NES −1.58 p.adjust 4.6e−03
C5=GOMF_TRANSLATION_REGULATOR_ACTIVITY_NUCLEIC_ACID_BINDING
NES −1.91 p.adjust 6.6e−05
C5=GOMF_TRANSLATION_REGULATOR_ACTIVITY_NUCLEIC_ACID_BINDING
NES −2.41 p.adjust 4.1e−10
C5=GOMF_TRANSLATION_ELONGATION_FACTOR_ACTIVITY
NES −1.82 p.adjust 3.4e−03
C5=GOMF_TRANSLATION_ELONGATION_FACTOR_ACTIVITY
NES −2.07 p.adjust 9.0e−04
C5=GOMF_TRANSLATION_ELONGATION_FACTOR_ACTIVITY
NES −2.01 p.adjust 3.0e−04
C5=GOBP_RRNA_METABOLIC_PROCESS
NES −1.84 p.adjust 1.2e−06
C5=GOBP_RRNA_METABOLIC_PROCESS
NES −1.69 p.adjust 6.6e−05
C5=GOBP_RRNA_METABOLIC_PROCESS
NES −1.53 p.adjust 1.1e−03
Running Enrichment Score
Ranked List Metric
Rank in Ordered Dataset

**Supplementary Figure 3. "Translation/ribosomal structure & function" GSEA profiles in liver, adipose tissue and skeletal muscle from 5 month old *Mlx*KO mice relative to WT controls**

Normalized enrichment scores and q values are indicated in the upper right corner of each profile. Data used to generate the ridge plots shown in Figure 6A are re-graphed and included here along with additional representative profiles. See Supplementary File 1 for a complete list of all gene sets of significance included in this category.

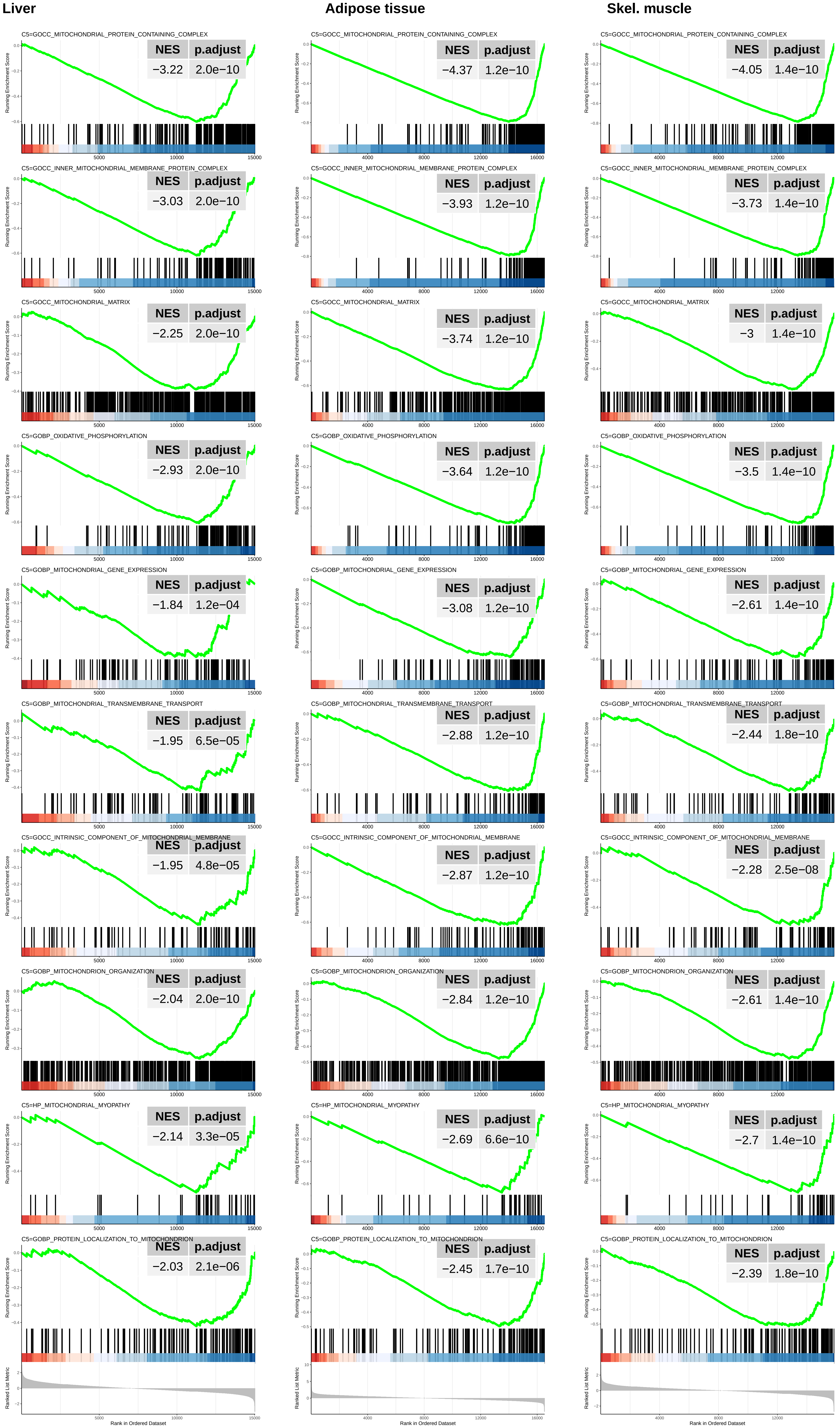
Liver
Adipose tissue
Skel. muscle
C5=GOCC_MITOCHONDRIAL_PROTEIN_CONTAINING_COMPLEX
NES p.adjust
−3.22 2.0e−10
−4.37 1.2e−10
−4.05 1.4e−10
C5=GOCC_INNER_MITOCHONDRIAL_MEMBRANE_PROTEIN_COMPLEX
−3.03 2.0e−10
−3.93 1.2e−10
−3.73 1.4e−10
C5=GOCC_MITOCHONDRIAL_MATRIX
−2.25 2.0e−10
−3.74 1.2e−10
−3 1.4e−10
C5=GOBP_OXIDATIVE_PHOSPHORYLATION
−2.93 2.0e−10
−3.64 1.2e−10
−3.5 1.4e−10
C5=GOBP_MITOCHONDRIAL_GENE_EXPRESSION
−1.84 1.2e−04
−3.08 1.2e−10
−2.61 1.4e−10
C5=GOBP_MITOCHONDRIAL_TRANSMEMBRANE_TRANSPORT
−1.95 6.5e−05
−2.88 1.2e−10
−2.44 1.8e−10
C5=GOCC_INTRINSIC_COMPONENT_OF_MITOCHONDRIAL_MEMBRANE
−1.95 4.8e−05
−2.87 1.2e−10
−2.28 2.5e−08
C5=GOBP_MITOCHONDRION_ORGANIZATION
−2.04 2.0e−10
−2.84 1.2e−10
−2.61 1.4e−10
C5=HP_MITOCHONDRIAL_MYOPATHY
−2.14 3.3e−05
−2.69 6.6e−10
−2.7 1.4e−10
C5=GOBP_PROTEIN_LOCALIZATION_TO_MITOCHONDRION
−2.03 2.1e−06
−2.45 1.7e−10
−2.39 1.8e−10
Running Enrichment Score
Ranked List Metric
Rank in Ordered Dataset

**Supplementary Figure 4. "Mitochondrial structure & function" GSEA profiles in liver, adipose tissue and skeletal muscle from 5 month old *Mlx*KO mice relative to WT controls**

Normalized enrichment scores and q values are indicated in the upper right corner of each profile. Data used to generate the ridge plots shown in Figure 6A are re-graphed and included here along with additional representative profiles. See Supplementary File 1 for a complete list of all gene sets of significance included in this category.

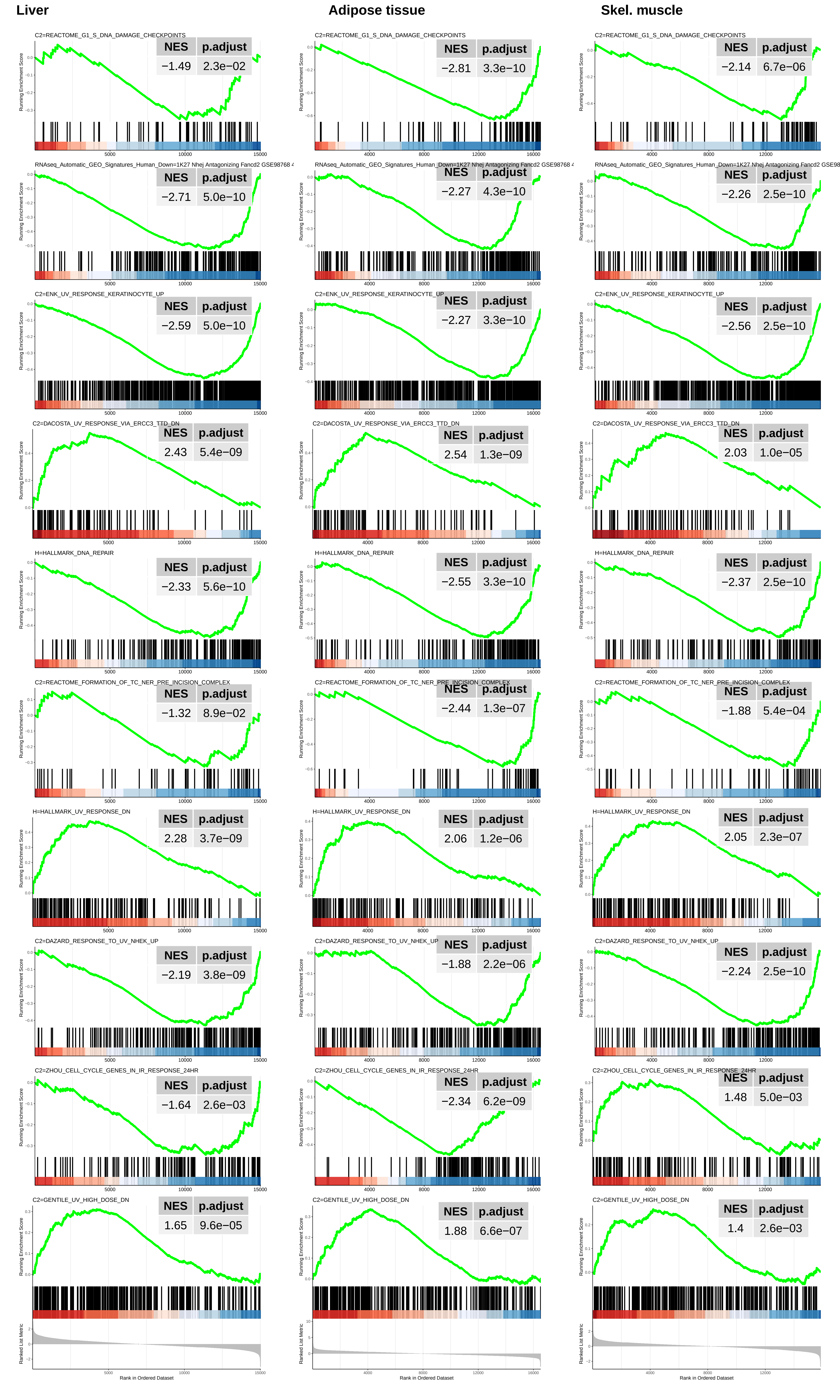
Liver
Adipose tissue
Skel. muscle
C2=REACTOME_G1_S_DNA_DAMAGE_CHECKPOINTS
NES p.adjust
−1.49 2.3e−02
−2.81 3.3e−10
−2.14 6.7e−06
RNAseq_Automatic_GEO_Signatures_Human_Down=1K27 Nhej Antagonizing Fancd2 GSE98768
NES p.adjust
−2.71 5.0e−10
−2.27 4.3e−10
−2.26 2.5e−10
C2=ENK_UV_RESPONSE_KERATINOCYTE_UP
NES p.adjust
−2.59 5.0e−10
−2.27 3.3e−10
−2.56 2.5e−10
C2=DACOSTA_UV_RESPONSE_VIA_ERCC3_TTD_DN
NES p.adjust
2.43 5.4e−09
2.54 1.3e−09
2.03 1.0e−05
H=HALLMARK_DNA_REPAIR
NES p.adjust
−2.33 5.6e−10
−2.55 3.3e−10
−2.37 2.5e−10
C2=REACTOME_FORMATION_OF_TC_NER_PRE_INCISION_COMPLEX
NES p.adjust
−1.32 8.9e−02
−2.44 1.3e−07
−1.88 5.4e−04
H=HALLMARK_UV_RESPONSE_DN
NES p.adjust
2.28 3.7e−09
2.06 1.2e−06
2.05 2.3e−07
C2=DAZARD_RESPONSE_TO_UV_NHEK_UP
NES p.adjust
−2.19 3.8e−09
−1.88 2.2e−06
−2.24 2.5e−10
C2=ZHOU_CELL_CYCLE_GENES_IN_IR_RESPONSE_24HR
NES p.adjust
−1.64 2.6e−03
−2.34 6.2e−09
1.48 5.0e−03
C2=GENTILE_UV_HIGH_DOSE_DN
NES p.adjust
1.65 9.6e−05
1.88 6.6e−07
1.4 2.6e−03
Running Enrichment Score
Ranked List Metric
Rank in Ordered Dataset

**Supplementary Figure 5. "DNA damage response and repair" GSEA profiles in liver, adipose tissue and skeletal muscle from 5 month old *Mlx*KO mice relative to WT controls**

Normalized enrichment scores and q values are indicated in the upper right corner of each profile. Data used to generate the ridge plots shown in Figure 6A are re-graphed and included here along with additional representative profiles. See Supplementary File 1 for a complete list of all gene sets of significance included in this category.

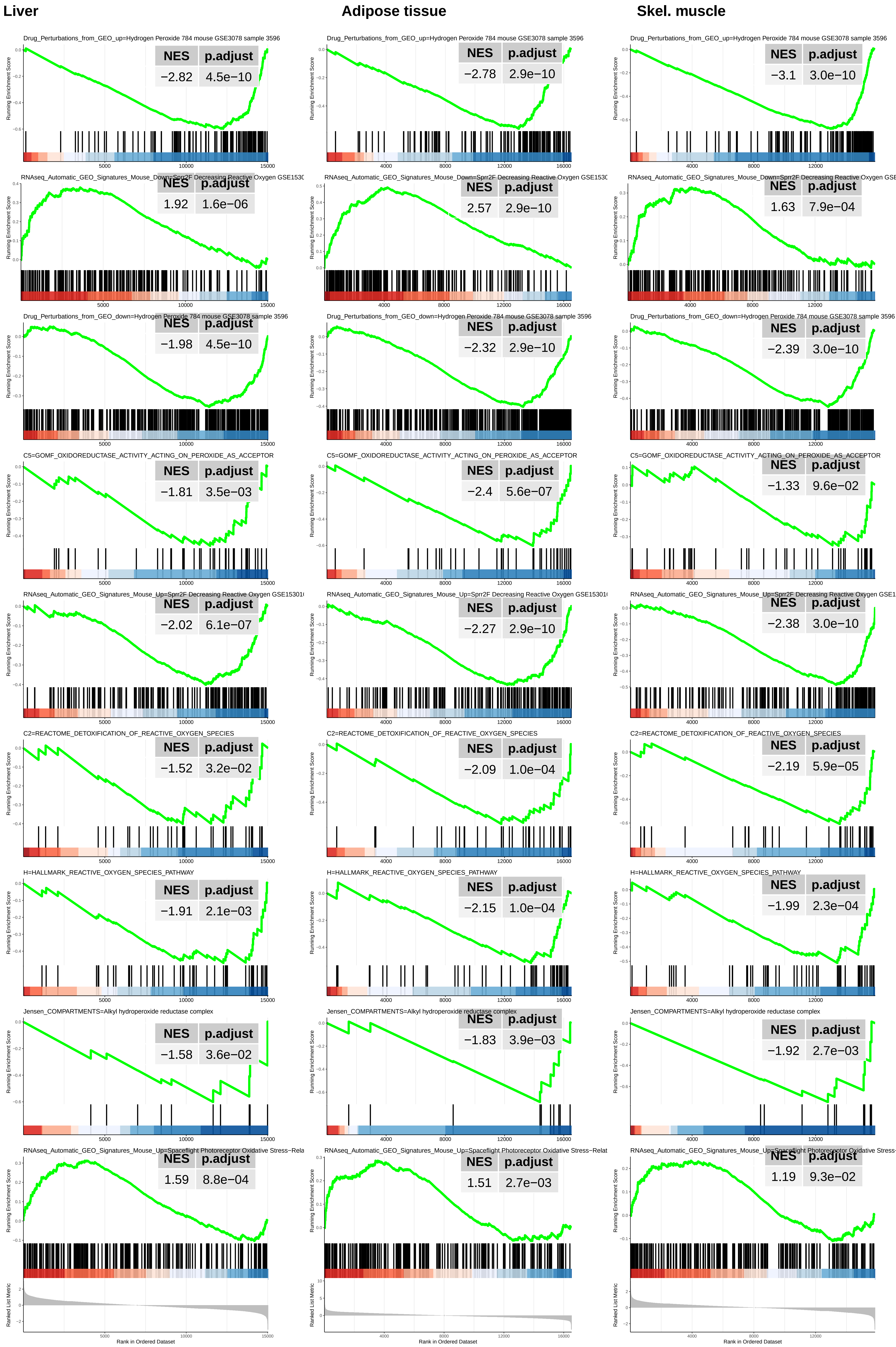

Liver
Adipose tissue
Skel. muscle
Drug_Perturbations_from_GEO_up=Hydrogen Peroxide 784 mouse GSE3078 sample 3596
NES p.adjust
−2.82 4.5e−10
−2.78 2.9e−10
−3.1 3.0e−10
RNAseq_Automatic_GEO_Signatures_Mouse_Down=Sprr2F Decreasing Reactive Oxygen GSE1530
1.92 1.6e−06
2.57 2.9e−10
1.63 7.9e−04
Drug_Perturbations_from_GEO_down=Hydrogen Peroxide 784 mouse GSE3078 sample 3596
−1.98 4.5e−10
−2.32 2.9e−10
−2.39 3.0e−10
C5=GOMF_OXIDOREDUCTASE_ACTIVITY_ACTING_ON_PEROXIDE_AS_ACCEPTOR
−1.81 3.5e−03
−2.4 5.6e−07
−1.33 9.6e−02
RNAseq_Automatic_GEO_Signatures_Mouse_Up=Sprr2F Decreasing Reactive Oxygen GSE15301
−2.02 6.1e−07
−2.27 2.9e−10
−2.38 3.0e−10
C2=REACTOME_DETOXIFICATION_OF_REACTIVE_OXYGEN_SPECIES
−1.52 3.2e−02
−2.09 1.0e−04
−2.19 5.9e−05
H=HALLMARK_REACTIVE_OXYGEN_SPECIES_PATHWAY
−1.91 2.1e−03
−2.15 1.0e−04
−1.99 2.3e−04
Jensen_COMPARTMENTS=Alkyl hydroperoxide reductase complex
−1.58 3.6e−02
−1.83 3.9e−03
−1.92 2.7e−03
RNAseq_Automatic_GEO_Signatures_Mouse_Up=Spaceflight Photoreceptor Oxidative Stress−Rela
1.59 8.8e−04
1.51 2.7e−03
1.19 9.3e−02
Running Enrichment Score
Ranked List Metric
Rank in Ordered Dataset

**Supplementary Figure 6. "Oxidative stress response" GSEA profiles in liver, adipose tissue and skeletal muscle from 5 month old *Mlx*KO mice relative to WT controls**

Normalized enrichment scores and q values are indicated in the upper right corner of each profile. Data used to generate the ridge plots shown in Figure 6A are re-graphed and included here along with additional representative profiles. See Supplementary File 1 for a complete list of all gene sets of significance included in this category.

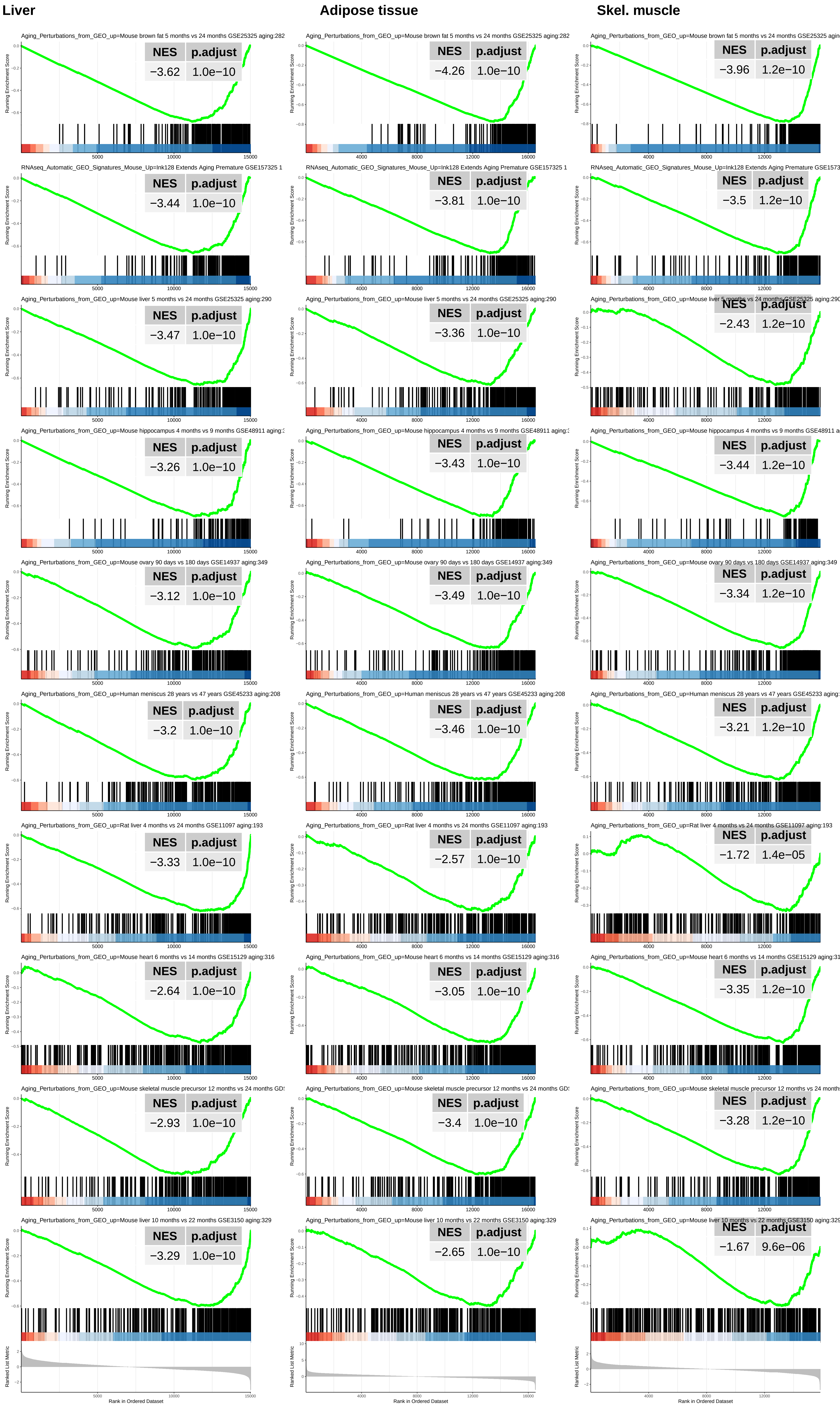

Liver
Adipose tissue
Skel. muscle
Aging_Perturbations_from_GEO_up=Mouse brown fat 5 months vs 24 months GSE25325 aging:282
NES p.adjust
−3.62 1.0e−10
−4.26 1.0e−10
−3.96 1.2e−10
RNAseq_Automatic_GEO_Signatures_Mouse_Up=Ink128 Extends Aging Premature GSE157325 1
−3.44 1.0e−10
−3.81 1.0e−10
−3.5 1.2e−10
Aging_Perturbations_from_GEO_up=Mouse liver 5 months vs 24 months GSE25325 aging:290
−3.47 1.0e−10
−3.36 1.0e−10
−2.43 1.2e−10
Aging_Perturbations_from_GEO_up=Mouse hippocampus 4 months vs 9 months GSE48911 aging:
−3.26 1.0e−10
−3.43 1.0e−10
−3.44 1.2e−10
Aging_Perturbations_from_GEO_up=Mouse ovary 90 days vs 180 days GSE14937 aging:349
−3.12 1.0e−10
−3.49 1.0e−10
−3.34 1.2e−10
Aging_Perturbations_from_GEO_up=Human meniscus 28 years vs 47 years GSE45233 aging:208
−3.2 1.0e−10
−3.46 1.0e−10
−3.21 1.2e−10
Aging_Perturbations_from_GEO_up=Rat liver 4 months vs 24 months GSE11097 aging:193
−3.33 1.0e−10
−2.57 1.0e−10
−1.72 1.4e−05
Aging_Perturbations_from_GEO_up=Mouse heart 6 months vs 14 months GSE15129 aging:316
−2.64 1.0e−10
−3.05 1.0e−10
−3.35 1.2e−10
Aging_Perturbations_from_GEO_up=Mouse skeletal muscle precursor 12 months vs 24 months GD
−2.93 1.0e−10
−3.4 1.0e−10
−3.28 1.2e−10
Aging_Perturbations_from_GEO_up=Mouse liver 10 months vs 22 months GSE3150 aging:329
−3.29 1.0e−10
−2.65 1.0e−10
−1.67 9.6e−06
Running Enrichment Score
Ranked List Metric
Rank in Ordered Dataset

**Supplementary Figure 7. "Aging" GSEA profiles in liver, adipose tissue and skeletal muscle from 5 month old *Mlx*KO mice relative to WT controls**

Normalized enrichment scores and q values are indicated in the upper right corner of each profile. Data used to generate the ridge plots shown in Figure 6A are re-graphed and included here along with additional representative profiles. See Supplementary File 1 for a complete list of all gene sets of significance included in this category.

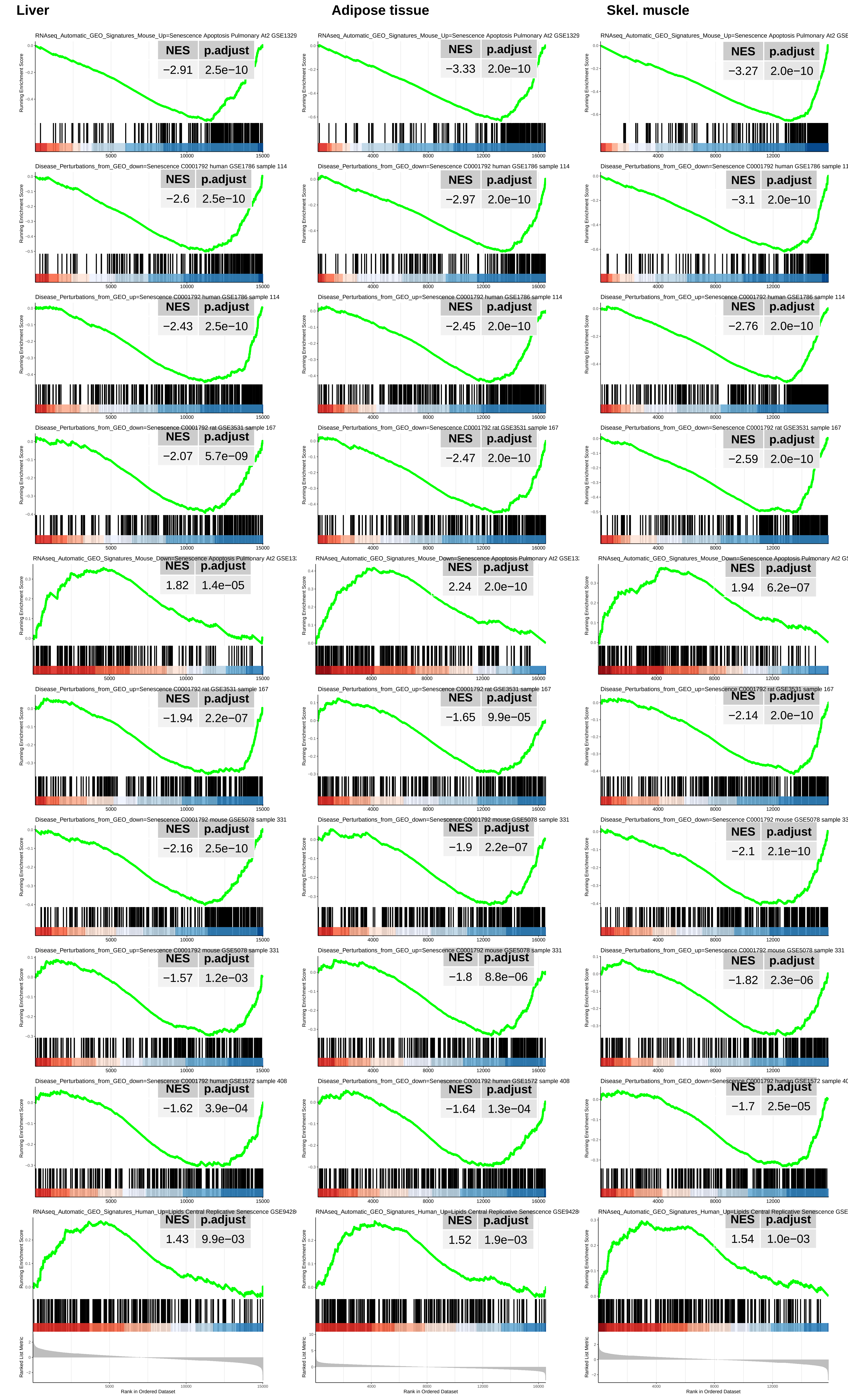

Liver
Adipose tissue
Skel. muscle
RNAseq_Automatic_GEO_Signatures_Mouse_Up=Senescence Apoptosis Pulmonary At2 GSE1329
NES p.adjust −2.91 2.5e−10
NES p.adjust −3.33 2.0e−10
NES p.adjust −3.27 2.0e−10
Disease_Perturbations_from_GEO_down=Senescence C0001792 human GSE1786 sample 114
NES p.adjust −2.6 2.5e−10
NES p.adjust −2.97 2.0e−10
NES p.adjust −3.1 2.0e−10
Disease_Perturbations_from_GEO_up=Senescence C0001792 human GSE1786 sample 114
NES p.adjust −2.43 2.5e−10
NES p.adjust −2.45 2.0e−10
NES p.adjust −2.76 2.0e−10
Disease_Perturbations_from_GEO_down=Senescence C0001792 rat GSE3531 sample 167
NES p.adjust −2.07 5.7e−09
NES p.adjust −2.47 2.0e−10
NES p.adjust −2.59 2.0e−10
RNAseq_Automatic_GEO_Signatures_Mouse_Down=Senescence Apoptosis Pulmonary At2 GSE13
NES p.adjust 1.82 1.4e−05
NES p.adjust 2.24 2.0e−10
NES p.adjust 1.94 6.2e−07
Disease_Perturbations_from_GEO_up=Senescence C0001792 rat GSE3531 sample 167
NES p.adjust −1.94 2.2e−07
NES p.adjust −1.65 9.9e−05
NES p.adjust −2.14 2.0e−10
Disease_Perturbations_from_GEO_down=Senescence C0001792 mouse GSE5078 sample 331
NES p.adjust −2.16 2.5e−10
NES p.adjust −1.9 2.2e−07
NES p.adjust −2.1 2.1e−10
Disease_Perturbations_from_GEO_up=Senescence C0001792 mouse GSE5078 sample 331
NES p.adjust −1.57 1.2e−03
NES p.adjust −1.8 8.8e−06
NES p.adjust −1.82 2.3e−06
Disease_Perturbations_from_GEO_down=Senescence C0001792 human GSE1572 sample 408
NES p.adjust −1.62 3.9e−04
NES p.adjust −1.64 1.3e−04
NES p.adjust −1.7 2.5e−05
RNAseq_Automatic_GEO_Signatures_Human_Up=Lipids Central Replicative Senescence GSE9428
NES p.adjust 1.43 9.9e−03
NES p.adjust 1.52 1.9e−03
NES p.adjust 1.54 1.0e−03
Running Enrichment Score
Ranked List Metric
Rank in Ordered Dataset

**Supplementary Figure 8. “Senescence” GSEA profiles in liver, adipose tissue and skeletal muscle from 5 month old *Mlx*KO mice relative to WT controls**

Normalized enrichment scores and q values are indicated in the upper right corner of each profile. Data used to generate the ridge plots shown in Figure 6A are re-graphed and included here along with additional representative profiles. See Supplementary File 1 for a complete list of all gene sets of significance included in this category.

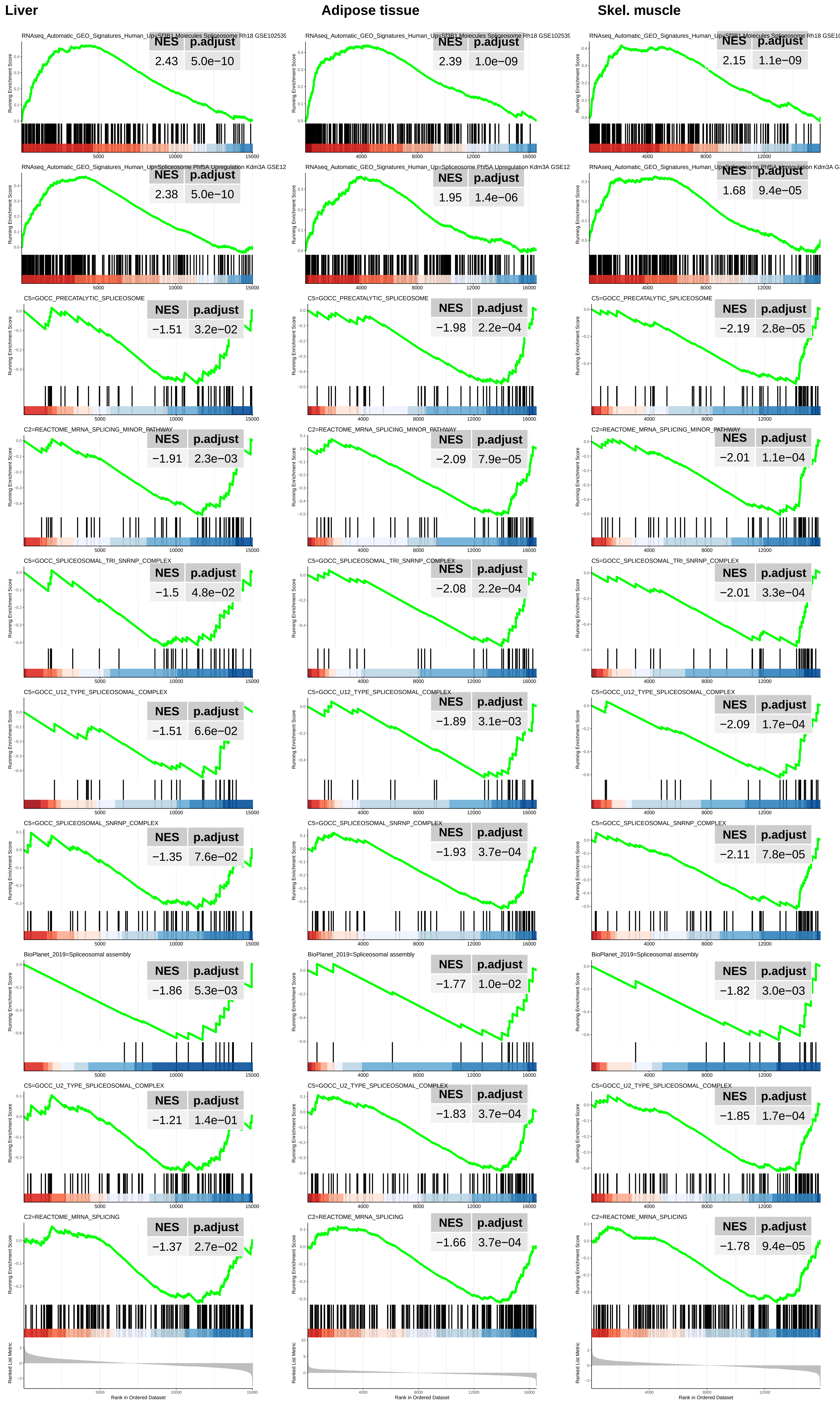
Liver
Adipose tissue
Skel. muscle
RNAseq_Automatic_GEO_Signatures_Human_Up=Sf3B1 Molecules Spliceosome Rh18 GSE10253
NES 2.43 p.adjust 5.0e−10
NES 2.39 p.adjust 1.0e−09
NES 2.15 p.adjust 1.1e−09
RNAseq_Automatic_GEO_Signatures_Human_Up=Spliceosome Phf5A Upregulation Kdm3A GSE12
NES 2.38 p.adjust 5.0e−10
NES 1.95 p.adjust 1.4e−06
NES 1.68 p.adjust 9.4e−05
C5=GOCC_PRECATALYTIC_SPLICEOSOME
NES −1.51 p.adjust 3.2e−02
NES −1.98 p.adjust 2.2e−04
NES −2.19 p.adjust 2.8e−05
C2=REACTOME_MRNA_SPLICING_MINOR_PATHWAY
NES −1.91 p.adjust 2.3e−03
NES −2.09 p.adjust 7.9e−05
NES −2.01 p.adjust 1.1e−04
C5=GOCC_SPLICEOSOMAL_TRI_SNRNP_COMPLEX
NES −1.5 p.adjust 4.8e−02
NES −2.08 p.adjust 2.2e−04
NES −2.01 p.adjust 3.3e−04
C5=GOCC_U12_TYPE_SPLICEOSOMAL_COMPLEX
NES −1.51 p.adjust 6.6e−02
NES −1.89 p.adjust 3.1e−03
NES −2.09 p.adjust 1.7e−04
C5=GOCC_SPLICEOSOMAL_SNRNP_COMPLEX
NES −1.35 p.adjust 7.6e−02
NES −1.93 p.adjust 3.7e−04
NES −2.11 p.adjust 7.8e−05
BioPlanet_2019=Spliceosomal assembly
NES −1.86 p.adjust 5.3e−03
NES −1.77 p.adjust 1.0e−02
NES −1.82 p.adjust 3.0e−03
C5=GOCC_U2_TYPE_SPLICEOSOMAL_COMPLEX
NES −1.21 p.adjust 1.4e−01
NES −1.83 p.adjust 3.7e−04
NES −1.85 p.adjust 1.7e−04
C2=REACTOME_MRNA_SPLICING
NES −1.37 p.adjust 2.7e−02
NES −1.66 p.adjust 3.7e−04
NES −1.78 p.adjust 9.4e−05
Running Enrichment Score
Ranked List Metric
Rank in Ordered Dataset

**Supplementary Figure 9. "mRNA splicing" GSEA profiles in liver, adipose tissue and skeletal muscle from 5 month old *Mlx*KO mice relative to WT controls**

Normalized enrichment scores and q values are indicated in the upper right corner of each profile. Data used to generate the ridge plots shown in Figure 6A are re-graphed and included here along with additional representative profiles. See Supplementary File 1 for a complete list of all gene sets of significance included in this category.

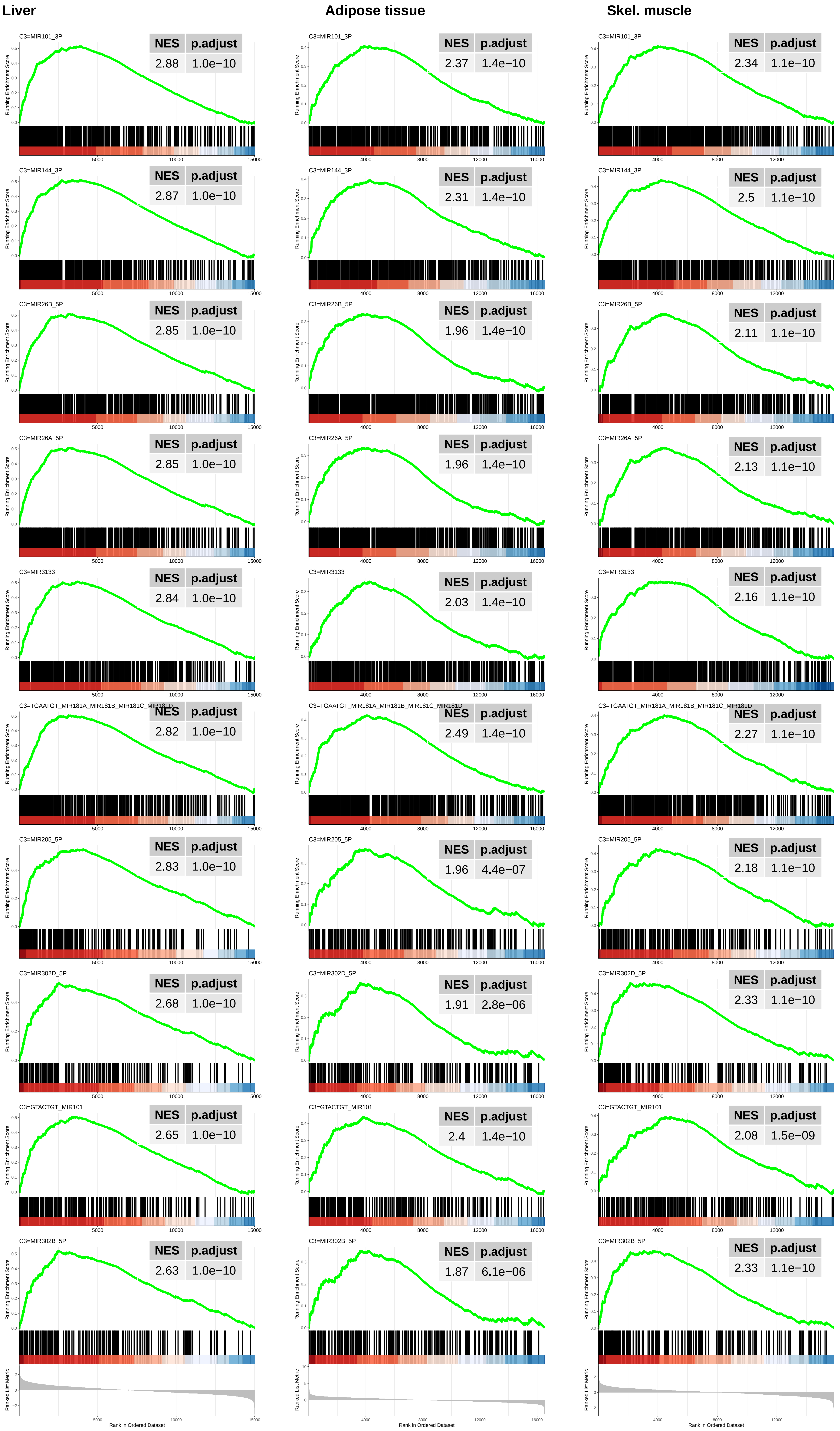

Liver
Adipose tissue
Skel. muscle
C3=MIR101_3P
NES p.adjust 2.88 1.0e−10
NES p.adjust 2.37 1.4e−10
NES p.adjust 2.34 1.1e−10
C3=MIR144_3P
NES p.adjust 2.87 1.0e−10
NES p.adjust 2.31 1.4e−10
NES p.adjust 2.5 1.1e−10
C3=MIR26B_5P
NES p.adjust 2.85 1.0e−10
NES p.adjust 1.96 1.4e−10
NES p.adjust 2.11 1.1e−10
C3=MIR26A_5P
NES p.adjust 2.85 1.0e−10
NES p.adjust 1.96 1.4e−10
NES p.adjust 2.13 1.1e−10
C3=MIR3133
NES p.adjust 2.84 1.0e−10
NES p.adjust 2.03 1.4e−10
NES p.adjust 2.16 1.1e−10
C3=TGAATGT_MIR181A_MIR181B_MIR181C_MIR181D
NES p.adjust 2.82 1.0e−10
NES p.adjust 2.49 1.4e−10
NES p.adjust 2.27 1.1e−10
C3=MIR205_5P
NES p.adjust 2.83 1.0e−10
NES p.adjust 1.96 4.4e−07
NES p.adjust 2.18 1.1e−10
C3=MIR302D_5P
NES p.adjust 2.68 1.0e−10
NES p.adjust 1.91 2.8e−06
NES p.adjust 2.33 1.1e−10
C3=GTACTGT_MIR101
NES p.adjust 2.65 1.0e−10
NES p.adjust 2.4 1.4e−10
NES p.adjust 2.08 1.5e−09
C3=MIR302B_5P
NES p.adjust 2.63 1.0e−10
NES p.adjust 1.87 6.1e−06
NES p.adjust 2.33 1.1e−10
Running Enrichment Score
Ranked List Metric
Rank in Ordered Dataset

**Supplementary Figure 10. "miRNA Targets" GSEA profiles in liver, adipose tissue and skeletal muscle from 5 month old *Mlx*KO mice relative to WT controls.**

Normalized enrichment scores (NESs) and q values are indicated in the upper right corner of each profile. Data used to generate the ridgeline plots shown in Figure 6B are re-graphed and included here along with additional representative GSEA profiles. See Supplementary File 2 for a complete list of all gene sets of significance included in this category.

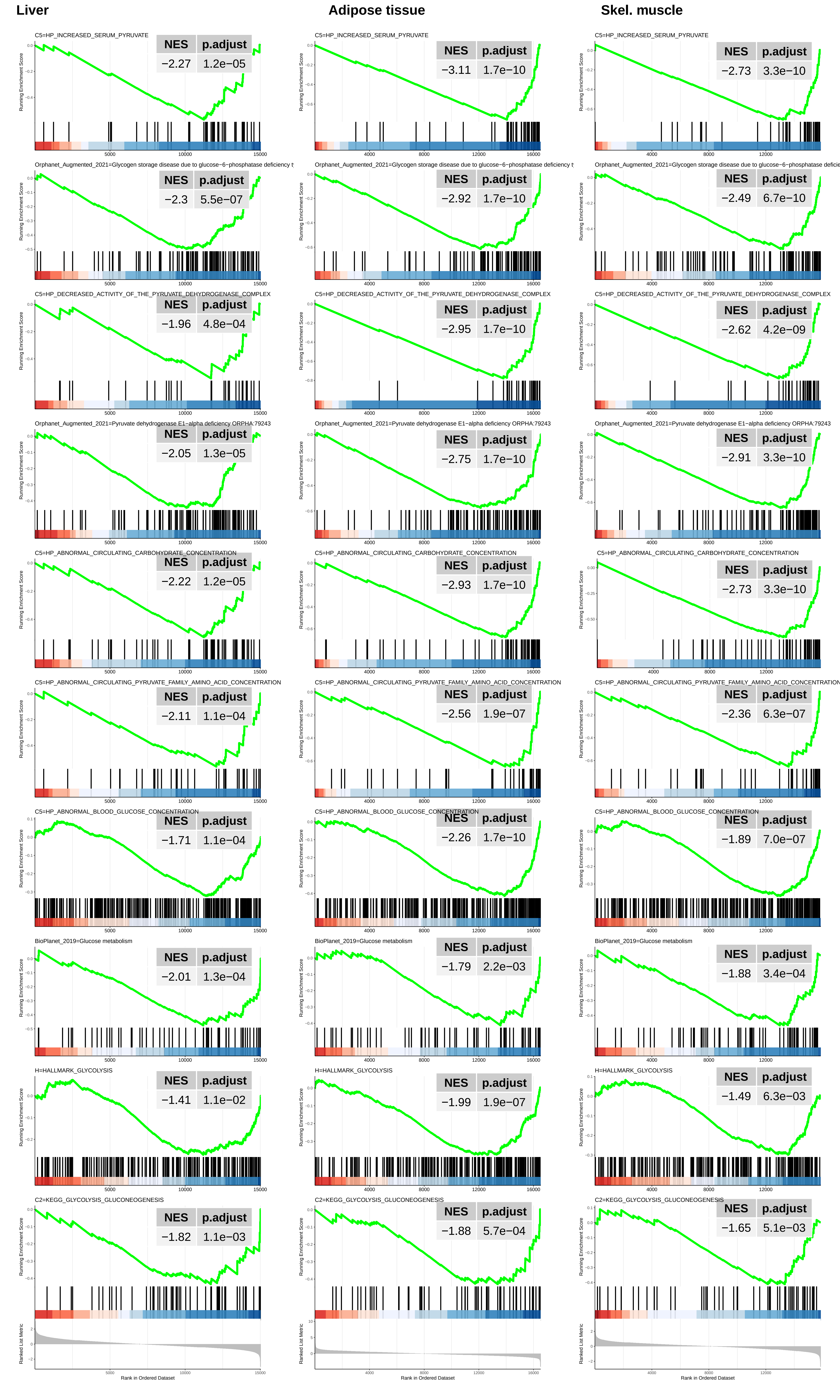
Liver
Adipose tissue
Skel. muscle
C5=HP_INCREASED_SERUM_PYRUVATE
NES p.adjust
−2.27 1.2e−05
C5=HP_INCREASED_SERUM_PYRUVATE
NES p.adjust
−3.11 1.7e−10
C5=HP_INCREASED_SERUM_PYRUVATE
NES p.adjust
−2.73 3.3e−10
Orphanet_Augmented_2021=Glycogen storage disease due to glucose−6−phosphatase deficiency t
NES p.adjust
−2.3 5.5e−07
Orphanet_Augmented_2021=Glycogen storage disease due to glucose−6−phosphatase deficiency t
NES p.adjust
−2.92 1.7e−10
Orphanet_Augmented_2021=Glycogen storage disease due to glucose−6−phosphatase deficie
NES p.adjust
−2.49 6.7e−10
C5=HP_DECREASED_ACTIVITY_OF_THE_PYRUVATE_DEHYDROGENASE_COMPLEX
NES p.adjust
−1.96 4.8e−04
C5=HP_DECREASED_ACTIVITY_OF_THE_PYRUVATE_DEHYDROGENASE_COMPLEX
NES p.adjust
−2.95 1.7e−10
C5=HP_DECREASED_ACTIVITY_OF_THE_PYRUVATE_DEHYDROGENASE_COMPLEX
NES p.adjust
−2.62 4.2e−09
Orphanet_Augmented_2021=Pyruvate dehydrogenase E1−alpha deficiency ORPHA:79243
NES p.adjust
−2.05 1.3e−05
Orphanet_Augmented_2021=Pyruvate dehydrogenase E1−alpha deficiency ORPHA:79243
NES p.adjust
−2.75 1.7e−10
Orphanet_Augmented_2021=Pyruvate dehydrogenase E1−alpha deficiency ORPHA:79243
NES p.adjust
−2.91 3.3e−10
C5=HP_ABNORMAL_CIRCULATING_CARBOHYDRATE_CONCENTRATION
NES p.adjust
−2.22 1.2e−05
C5=HP_ABNORMAL_CIRCULATING_CARBOHYDRATE_CONCENTRATION
NES p.adjust
−2.93 1.7e−10
C5=HP_ABNORMAL_CIRCULATING_CARBOHYDRATE_CONCENTRATION
NES p.adjust
−2.73 3.3e−10
C5=HP_ABNORMAL_CIRCULATING_PYRUVATE_FAMILY_AMINO_ACID_CONCENTRATION
NES p.adjust
−2.11 1.1e−04
C5=HP_ABNORMAL_CIRCULATING_PYRUVATE_FAMILY_AMINO_ACID_CONCENTRATION
NES p.adjust
−2.56 1.9e−07
C5=HP_ABNORMAL_CIRCULATING_PYRUVATE_FAMILY_AMINO_ACID_CONCENTRATION
NES p.adjust
−2.36 6.3e−07
C5=HP_ABNORMAL_BLOOD_GLUCOSE_CONCENTRATION
NES p.adjust
−1.71 1.1e−04
C5=HP_ABNORMAL_BLOOD_GLUCOSE_CONCENTRATION
NES p.adjust
−2.26 1.7e−10
C5=HP_ABNORMAL_BLOOD_GLUCOSE_CONCENTRATION
NES p.adjust
−1.89 7.0e−07
BioPlanet_2019=Glucose metabolism
NES p.adjust
−2.01 1.3e−04
BioPlanet_2019=Glucose metabolism
NES p.adjust
−1.79 2.2e−03
BioPlanet_2019=Glucose metabolism
NES p.adjust
−1.88 3.4e−04
H=HALLMARK_GLYCOLYSIS
NES p.adjust
−1.41 1.1e−02
H=HALLMARK_GLYCOLYSIS
NES p.adjust
−1.99 1.9e−07
H=HALLMARK_GLYCOLYSIS
NES p.adjust
−1.49 6.3e−03
C2=KEGG_GLYCOLYSIS_GLUCONEOGENESIS
NES p.adjust
−1.82 1.1e−03
C2=KEGG_GLYCOLYSIS_GLUCONEOGENESIS
NES p.adjust
−1.88 5.7e−04
C2=KEGG_GLYCOLYSIS_GLUCONEOGENESIS
NES p.adjust
−1.65 5.1e−03
Running Enrichment Score
Ranked List Metric
Rank in Ordered Dataset

**Supplementary Figure 11. "Glycolysis-related" GSEA profiles in liver, adipose tissue and skeletal muscle from 5 month old *Mlx*KO mice relative to WT controls.**

Normalized enrichment scores (NESs) and q values are indicated in the upper right corner of each profile. Data used to generate the ridgeline plots shown in Figure 6B are re-graphed and included here along with additional representative GSEA profiles. See Supplementary File 2 for a complete list of all gene sets of significance included in this category.

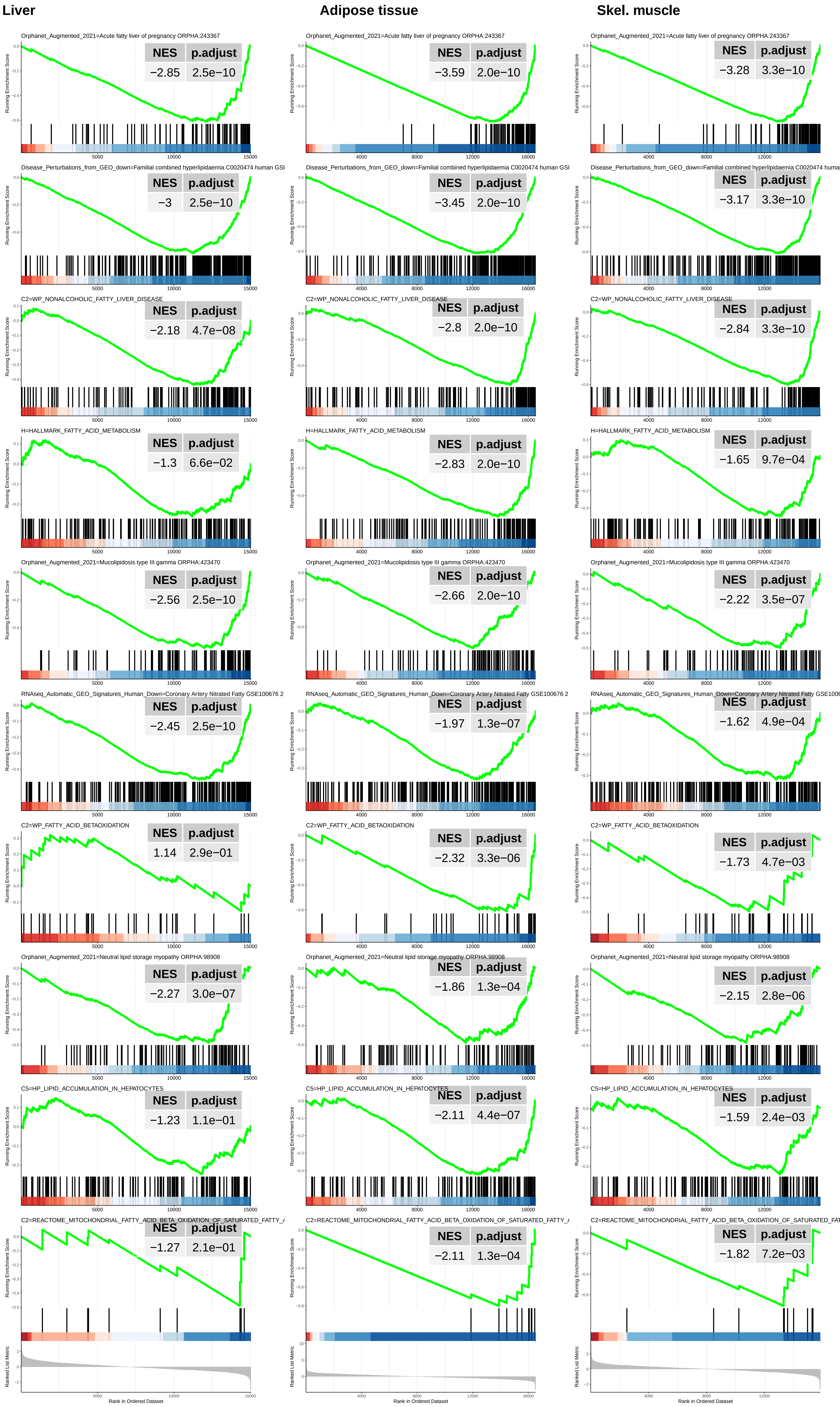

Liver
Adipose tissue
Skel. muscle
Orphanet_Augmented_2021=Acute fatty liver of pregnancy ORPHA:243367
NES p.adjust
−2.85 2.5e−10
−3.59 2.0e−10
−3.28 3.3e−10
Disease_Perturbations_from_GEO_down=Familial combined hyperlipidaemia C0020474 human GSI
NES p.adjust
−3 2.5e−10
−3.45 2.0e−10
−3.17 3.3e−10
C2=WP_NONALCOHOLIC_FATTY_LIVER_DISEASE
NES p.adjust
−2.18 4.7e−08
−2.8 2.0e−10
−2.84 3.3e−10
H=HALLMARK_FATTY_ACID_METABOLISM
NES p.adjust
−1.3 6.6e−02
−2.83 2.0e−10
−1.65 9.7e−04
Orphanet_Augmented_2021=Mucolipidosis type III gamma ORPHA:423470
NES p.adjust
−2.56 2.5e−10
−2.66 2.0e−10
−2.22 3.5e−07
RNAseq_Automatic_GEO_Signatures_Human_Down=Coronary Artery Nitrated Fatty GSE100676 2
NES p.adjust
−2.45 2.5e−10
−1.97 1.3e−07
−1.62 4.9e−04
C2=WP_FATTY_ACID_BETAOXIDATION
NES p.adjust
1.14 2.9e−01
−2.32 3.3e−06
−1.73 4.7e−03
Orphanet_Augmented_2021=Neutral lipid storage myopathy ORPHA:98908
NES p.adjust
−2.27 3.0e−07
−1.86 1.3e−04
−2.15 2.8e−06
C5=HP_LIPID_ACCUMULATION_IN_HEPATOCYTES
NES p.adjust
−1.23 1.1e−01
−2.11 4.4e−07
−1.59 2.4e−03
C2=REACTOME_MITOCHONDRIAL_FATTY_ACID_BETA_OXIDATION_OF_SATURATED_FATTY_
NES p.adjust
−1.27 2.1e−01
−2.11 1.3e−04
−1.82 7.2e−03
Running Enrichment Score
Ranked List Metric
Rank in Ordered Dataset

**Supplementary Figure 12. "Lipid metabolism" GSEA profiles in liver, adipose tissue and skeletal muscle from 5 month old *Mlx*KO mice relative to WT controls.**

Normalized enrichment scores (NESs) and q values are indicated in the upper right corner of each profile. Data used to generate the ridgeline plots shown in Figure 6B are re-graphed and included here along with additional representative GSEA profiles. See Supplementary File 2 for a complete list of all gene sets of significance included in this category.

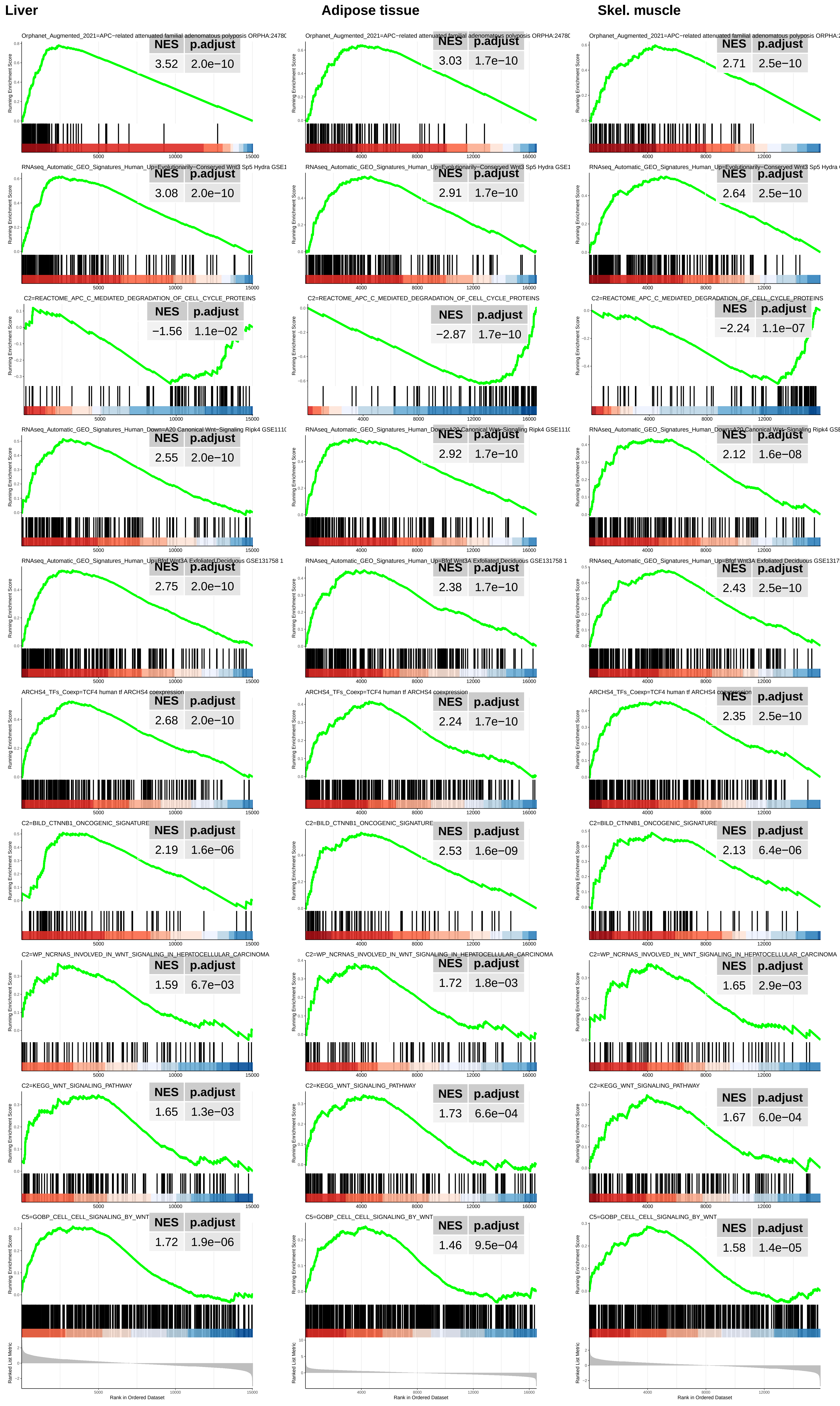

Liver
Adipose tissue
Skel. muscle
Orphanet_Augmented_2021=APC−related attenuated familial adenomatous polyposis ORPHA:24780
NES p.adjust
3.52 2.0e−10
3.03 1.7e−10
2.71 2.5e−10
RNAseq_Automatic_GEO_Signatures_Human_Up=Evolutionarily−Conserved Wnt3 Sp5 Hydra GSE1
3.08 2.0e−10
2.91 1.7e−10
2.64 2.5e−10
C2=REACTOME_APC_C_MEDIATED_DEGRADATION_OF_CELL_CYCLE_PROTEINS
−1.56 1.1e−02
−2.87 1.7e−10
−2.24 1.1e−07
RNAseq_Automatic_GEO_Signatures_Human_Down=A20 Canonical Wnt−Signaling Ripk4 GSE1110
2.55 2.0e−10
2.92 1.7e−10
2.12 1.6e−08
RNAseq_Automatic_GEO_Signatures_Human_Up=Bfgf Wnt3A Exfoliated Deciduous GSE131758 1
2.75 2.0e−10
2.38 1.7e−10
2.43 2.5e−10
ARCHS4_TFs_Coexp=TCF4 human tf ARCHS4 coexpression
2.68 2.0e−10
2.24 1.7e−10
2.35 2.5e−10
C2=BILD_CTNNB1_ONCOGENIC_SIGNATURE
2.19 1.6e−06
2.53 1.6e−09
2.13 6.4e−06
C2=WP_NCRNAS_INVOLVED_IN_WNT_SIGNALING_IN_HEPATOCELLULAR_CARCINOMA
1.59 6.7e−03
1.72 1.8e−03
1.65 2.9e−03
C2=KEGG_WNT_SIGNALING_PATHWAY
1.65 1.3e−03
1.73 6.6e−04
1.67 6.0e−04
C5=GOBP_CELL_CELL_SIGNALING_BY_WNT
1.72 1.9e−06
1.46 9.5e−04
1.58 1.4e−05
Running Enrichment Score
Ranked List Metric
Rank in Ordered Dataset

**Supplementary Figure 13. "Wnt/β-catenin/Tcf signaling" GSEA profiles in liver, adipose tissue and skeletal muscle from 5 month old *Mlx*KO mice relative to WT controls.**

Normalized enrichment scores (NESs) and q values are indicated in the upper right corner of each profile. Data used to generate the ridgeline plots shown in Figure 6B are re-graphed and included here along with additional representative GSEA profiles. See Supplementary File 2 for a complete list of all gene sets of significance included in this category.

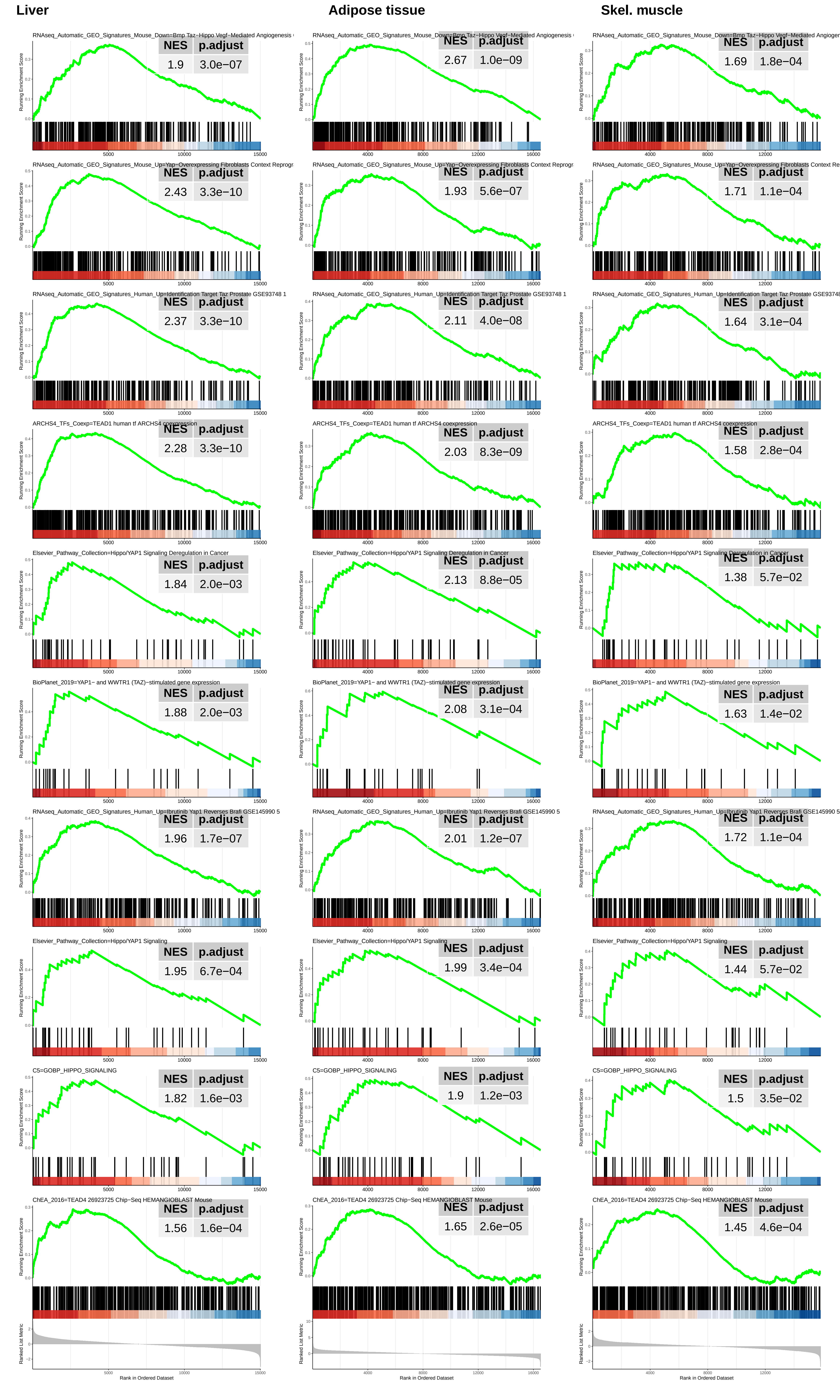
Liver
Adipose tissue
Skel. muscle
RNAseq_Automatic_GEO_Signatures_Mouse_Down=Bmp Taz−Hippo Vegf−Mediated Angiogenesis
NES p.adjust
1.9 3.0e−07
2.67 1.0e−09
1.69 1.8e−04
RNAseq_Automatic_GEO_Signatures_Mouse_Up=Yap−Overexpressing Fibroblasts Context Reprogr
2.43 3.3e−10
1.93 5.6e−07
1.71 1.1e−04
RNAseq_Automatic_GEO_Signatures_Human_Up=Identification Target Taz Prostate GSE93748 1
2.37 3.3e−10
2.11 4.0e−08
1.64 3.1e−04
ARCHS4_TFs_Coexp=TEAD1 human tf ARCHS4 coexpression
2.28 3.3e−10
2.03 8.3e−09
1.58 2.8e−04
Elsevier_Pathway_Collection=Hippo/YAP1 Signaling Deregulation in Cancer
1.84 2.0e−03
2.13 8.8e−05
1.38 5.7e−02
BioPlanet_2019=YAP1− and WWTR1 (TAZ)−stimulated gene expression
1.88 2.0e−03
2.08 3.1e−04
1.63 1.4e−02
RNAseq_Automatic_GEO_Signatures_Human_Up=Ibrutinib Yap1 Reverses Brafi GSE145990 5
1.96 1.7e−07
2.01 1.2e−07
1.72 1.1e−04
Elsevier_Pathway_Collection=Hippo/YAP1 Signaling
1.95 6.7e−04
1.99 3.4e−04
1.44 5.7e−02
C5=GOBP_HIPPO_SIGNALING
1.82 1.6e−03
1.9 1.2e−03
1.5 3.5e−02
ChEA_2016=TEAD4 26923725 Chip−Seq HEMANGIOBLAST Mouse
1.56 1.6e−04
1.65 2.6e−05
1.45 4.6e−04
Running Enrichment Score
Ranked List Metric
Rank in Ordered Dataset

**Supplementary Figure 14. “Hippo/YAP/TAZ signaling” GSEA profiles in liver, adipose tissue and skeletal muscle from 5 month old *Mlx*KO mice relative to WT controls.**

Normalized enrichment scores (NESs) and q values are indicated in the upper right corner of each profile. Data used to generate the ridgeline plots shown in Figure 6B are re-graphed and included here along with additional representative GSEA profiles. See Supplementary File 2 for a complete list of all gene sets of significance included in this category.

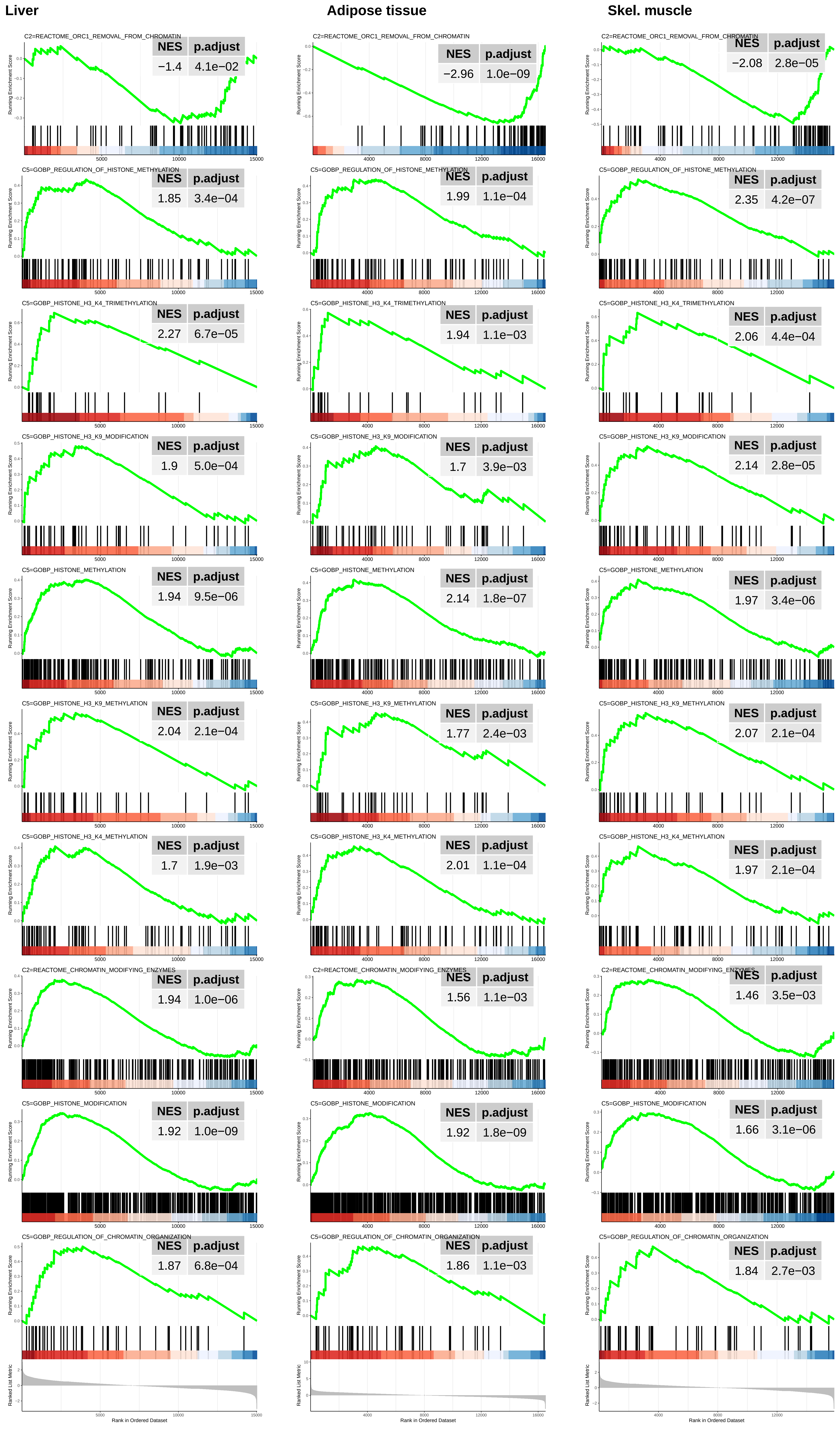
Liver
Adipose tissue
Skel. muscle
C2=REACTOME_ORC1_REMOVAL_FROM_CHROMATIN
NES p.adjust
−1.4 4.1e−02
−2.96 1.0e−09
−2.08 2.8e−05
C5=GOBP_REGULATION_OF_HISTONE_METHYLATION
1.85 3.4e−04
1.99 1.1e−04
2.35 4.2e−07
C5=GOBP_HISTONE_H3_K4_TRIMETHYLATION
2.27 6.7e−05
1.94 1.1e−03
2.06 4.4e−04
C5=GOBP_HISTONE_H3_K9_MODIFICATION
1.9 5.0e−04
1.7 3.9e−03
2.14 2.8e−05
C5=GOBP_HISTONE_METHYLATION
1.94 9.5e−06
2.14 1.8e−07
1.97 3.4e−06
C5=GOBP_HISTONE_H3_K9_METHYLATION
2.04 2.1e−04
1.77 2.4e−03
2.07 2.1e−04
C5=GOBP_HISTONE_H3_K4_METHYLATION
1.7 1.9e−03
2.01 1.1e−04
1.97 2.1e−04
C2=REACTOME_CHROMATIN_MODIFYING_ENZYMES
1.94 1.0e−06
1.56 1.1e−03
1.46 3.5e−03
C5=GOBP_HISTONE_MODIFICATION
1.92 1.0e−09
1.92 1.8e−09
1.66 3.1e−06
C5=GOBP_REGULATION_OF_CHROMATIN_ORGANIZATION
1.87 6.8e−04
1.86 1.1e−03
1.84 2.7e−03
Running Enrichment Score
Ranked List Metric
Rank in Ordered Dataset

**Supplementary Figure 15. "Chromatin/histone modification" GSEA profiles in liver, adipose tissue and skeletal muscle from 5 month old *Mlx*KO mice relative to WT controls.**

Normalized enrichment scores (NESs) and q values are indicated in the upper right corner of each profile. Data used to generate the ridgeline plots shown in Figure 6B are re-graphed and included here along with additional representative GSEA profiles. See Supplementary File 2 for a complete list of all gene sets of significance included in this category.

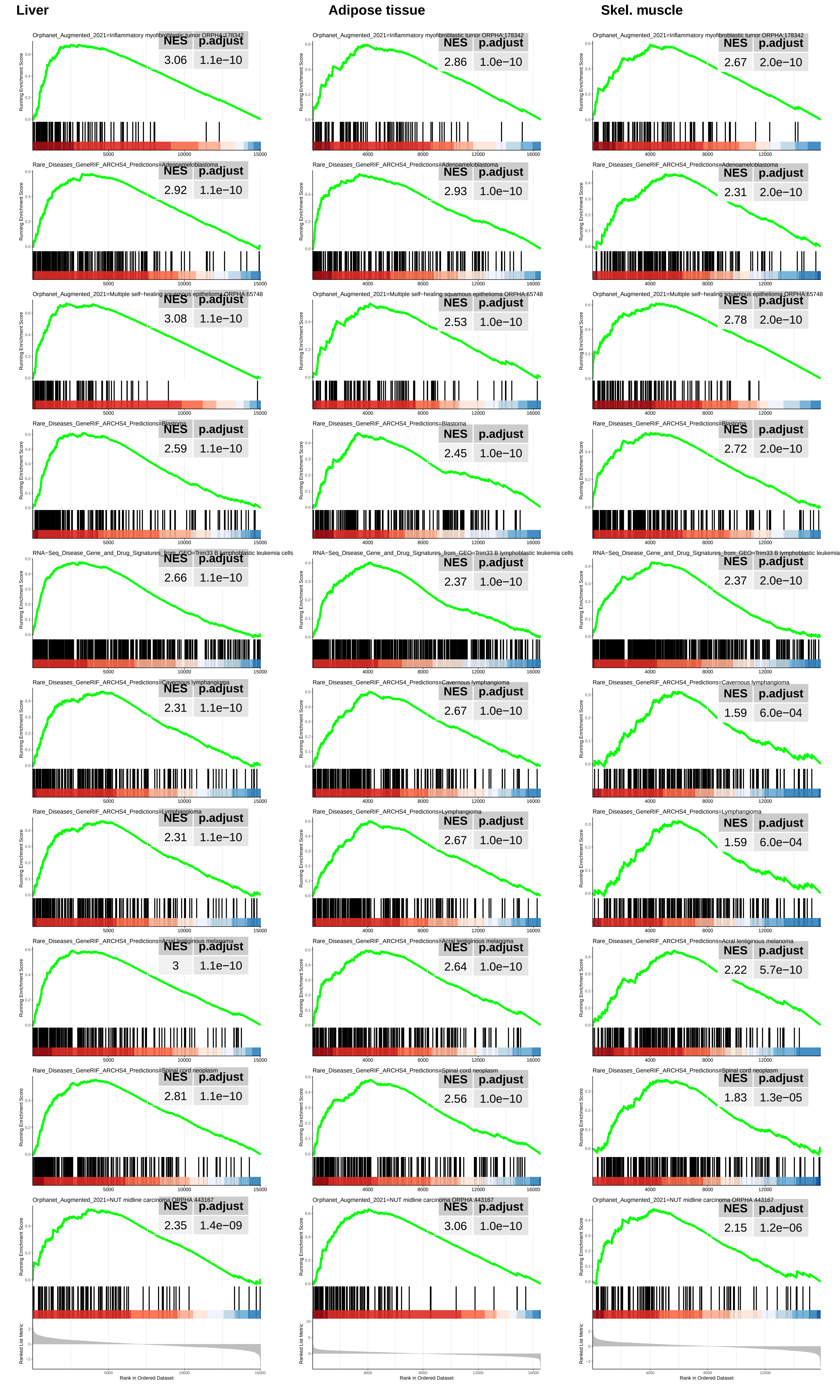

Liver
Adipose tissue
Skel. muscle
Orphanet_Augmented_2021=Inflammatory myofibroblastic tumor ORPHA:178342
NES 3.06 p.adjust 1.1e−10
NES 2.86 p.adjust 1.0e−10
NES 2.67 p.adjust 2.0e−10
Rare_Diseases_GeneRIF_ARCHS4_Predictions=Adenoameloblastoma
NES 2.92 p.adjust 1.1e−10
NES 2.93 p.adjust 1.0e−10
NES 2.31 p.adjust 2.0e−10
Orphanet_Augmented_2021=Multiple self−healing squamous epithelioma ORPHA:65748
NES 3.08 p.adjust 1.1e−10
NES 2.53 p.adjust 1.0e−10
NES 2.78 p.adjust 2.0e−10
Rare_Diseases_GeneRIF_ARCHS4_Predictions=Blastoma
NES 2.59 p.adjust 1.1e−10
NES 2.45 p.adjust 1.0e−10
NES 2.72 p.adjust 2.0e−10
RNA−Seq_Disease_Gene_and_Drug_Signatures_from_GEO=Trim33 B lymphoblastic leukemia cells
NES 2.66 p.adjust 1.1e−10
NES 2.37 p.adjust 1.0e−10
NES 2.37 p.adjust 2.0e−10
Rare_Diseases_GeneRIF_ARCHS4_Predictions=Cavernous lymphangioma
NES 2.31 p.adjust 1.1e−10
NES 2.67 p.adjust 1.0e−10
NES 1.59 p.adjust 6.0e−04
Rare_Diseases_GeneRIF_ARCHS4_Predictions=Lymphangioma
NES 2.31 p.adjust 1.1e−10
NES 2.67 p.adjust 1.0e−10
NES 1.59 p.adjust 6.0e−04
Rare_Diseases_GeneRIF_ARCHS4_Predictions=Acral lentiginous melanoma
NES 3 p.adjust 1.1e−10
NES 2.64 p.adjust 1.0e−10
NES 2.22 p.adjust 5.7e−10
Rare_Diseases_GeneRIF_ARCHS4_Predictions=Spinal cord neoplasm
NES 2.81 p.adjust 1.1e−10
NES 2.56 p.adjust 1.0e−10
NES 1.83 p.adjust 1.3e−05
Orphanet_Augmented_2021=NUT midline carcinoma ORPHA:443167
NES 2.35 p.adjust 1.4e−09
NES 3.06 p.adjust 1.0e−10
NES 2.15 p.adjust 1.2e−06
Running Enrichment Score
Ranked List Metric
Rank in Ordered Dataset

**Supplementary Figure 16.** "Cancer-related" GSEA profiles in liver, adipose tissue and skeletal muscle from 5 month old *Mlx*KO mice relative to WT controls

Normalized enrichment scores and q values are indicated in the upper right corner of each profile. Data used to generate the ridge plots shown in Figure 6C are re-graphed and included here along with additional representative profiles. See Supplementary File 3 for a complete list of all gene sets of significance included in this category.

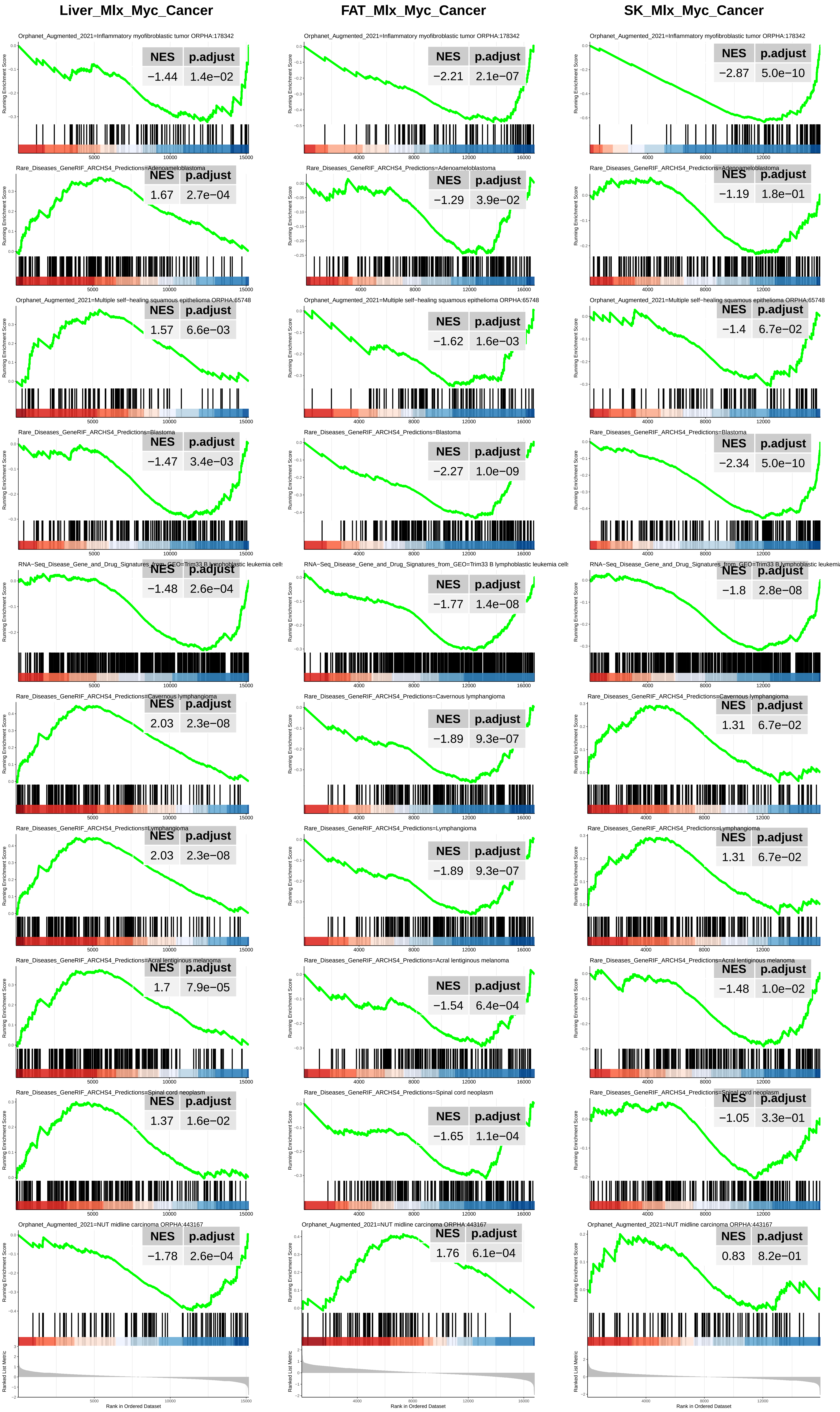

Liver_Mlx_Myc_Cancer
FAT_Mlx_Myc_Cancer
SK_Mlx_Myc_Cancer
Orphanet_Augmented_2021=Inflammatory myofibroblastic tumor ORPHA:178342
NES p.adjust
−1.44 1.4e−02
−2.21 2.1e−07
−2.87 5.0e−10
Rare_Diseases_GeneRIF_ARCHS4_Predictions=Adenoameloblastoma
1.67 2.7e−04
−1.29 3.9e−02
−1.19 1.8e−01
Orphanet_Augmented_2021=Multiple self−healing squamous epithelioma ORPHA:65748
1.57 6.6e−03
−1.62 1.6e−03
−1.4 6.7e−02
Rare_Diseases_GeneRIF_ARCHS4_Predictions=Blastoma
−1.47 3.4e−03
−2.27 1.0e−09
−2.34 5.0e−10
RNA−Seq_Disease_Gene_and_Drug_Signatures_from_GEO=Trim33 B lymphoblastic leukemia cells
−1.48 2.6e−04
−1.77 1.4e−08
−1.8 2.8e−08
Rare_Diseases_GeneRIF_ARCHS4_Predictions=Cavernous lymphangioma
2.03 2.3e−08
−1.89 9.3e−07
1.31 6.7e−02
Rare_Diseases_GeneRIF_ARCHS4_Predictions=Lymphangioma
2.03 2.3e−08
−1.89 9.3e−07
1.31 6.7e−02
Rare_Diseases_GeneRIF_ARCHS4_Predictions=Acral lentiginous melanoma
1.7 7.9e−05
−1.54 6.4e−04
−1.48 1.0e−02
Rare_Diseases_GeneRIF_ARCHS4_Predictions=Spinal cord neoplasm
1.37 1.6e−02
−1.65 1.1e−04
−1.05 3.3e−01
Orphanet_Augmented_2021=NUT midline carcinoma ORPHA:443167
−1.78 2.6e−04
1.76 6.1e−04
0.83 8.2e−01
Running Enrichment Score
Ranked List Metric
Rank in Ordered Dataset

**Supplementary Figure 17. “Cancer-related” GSEA profiles in liver, adipose tissue and skeletal muscle from 5 month old *Myc*KO mice relative to WT controls.**

Normalized enrichment scores and q values are indicated in the upper right corner of each profile. Data used to generate the ridge plots shown in Figure 6C are re-graphed and included here along with additional representative profiles. See Supplementary File 3 for a complete list of all gene sets of significance included in this category.

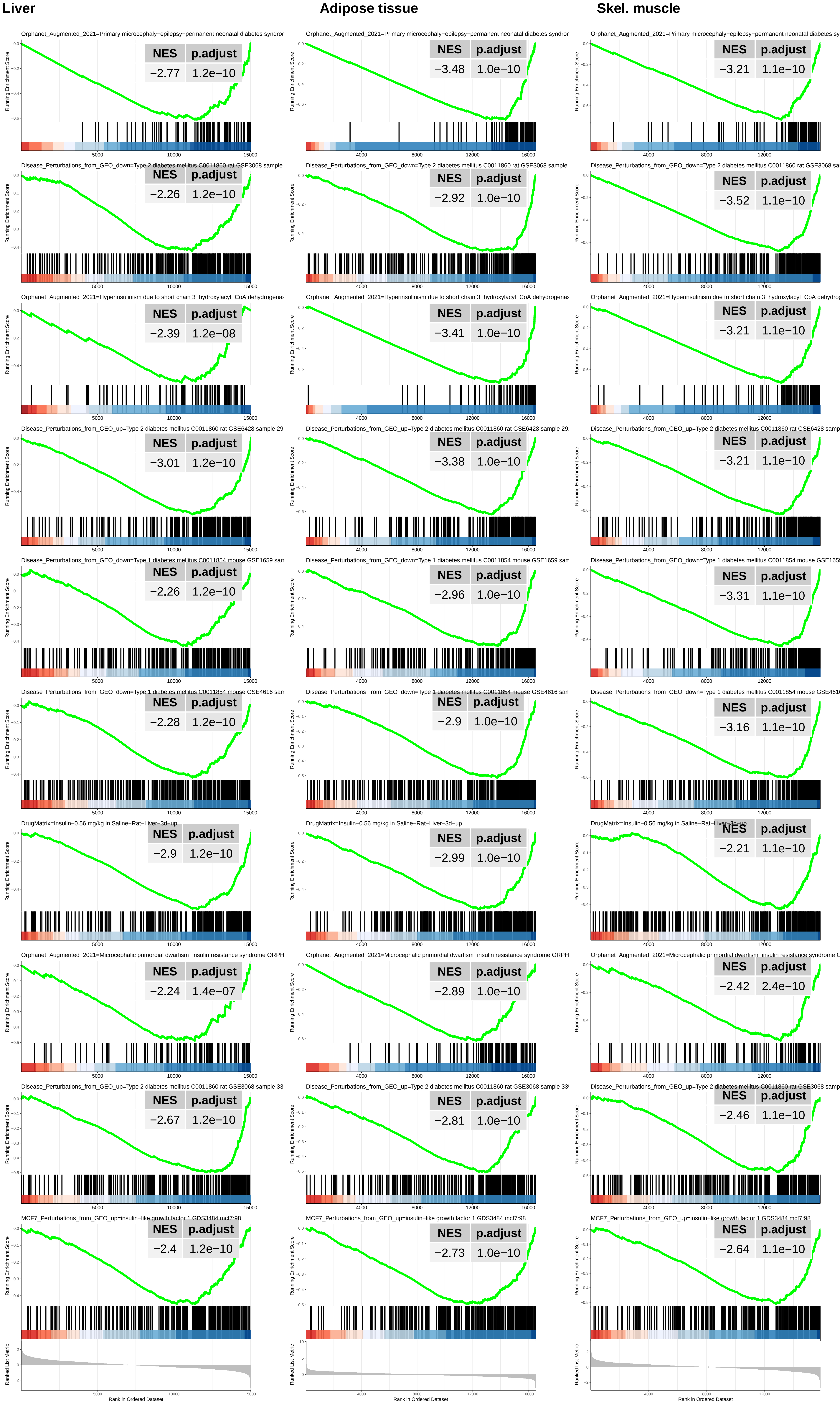

Liver
Adipose tissue
Skel. muscle
Orphanet_Augmented_2021=Primary microcephaly−epilepsy−permanent neonatal diabetes syndron
NES p.adjust
−2.77 1.2e−10
−3.48 1.0e−10
−3.21 1.1e−10
Disease_Perturbations_from_GEO_down=Type 2 diabetes mellitus C0011860 rat GSE3068 sample
NES p.adjust
−2.26 1.2e−10
−2.92 1.0e−10
−3.52 1.1e−10
Orphanet_Augmented_2021=Hyperinsulinism due to short chain 3−hydroxylacyl−CoA dehydrogenas
NES p.adjust
−2.39 1.2e−08
−3.41 1.0e−10
−3.21 1.1e−10
Disease_Perturbations_from_GEO_up=Type 2 diabetes mellitus C0011860 rat GSE6428 sample 29
NES p.adjust
−3.01 1.2e−10
−3.38 1.0e−10
−3.21 1.1e−10
Disease_Perturbations_from_GEO_down=Type 1 diabetes mellitus C0011854 mouse GSE1659 sam
NES p.adjust
−2.26 1.2e−10
−2.96 1.0e−10
−3.31 1.1e−10
Disease_Perturbations_from_GEO_down=Type 1 diabetes mellitus C0011854 mouse GSE4616 sam
NES p.adjust
−2.28 1.2e−10
−2.9 1.0e−10
−3.16 1.1e−10
DrugMatrix=Insulin−0.56 mg/kg in Saline−Rat−Liver−3d−up
NES p.adjust
−2.9 1.2e−10
−2.99 1.0e−10
−2.21 1.1e−10
Orphanet_Augmented_2021=Microcephalic primordial dwarfism−insulin resistance syndrome ORPH
NES p.adjust
−2.24 1.4e−07
−2.89 1.0e−10
−2.42 2.4e−10
Disease_Perturbations_from_GEO_up=Type 2 diabetes mellitus C0011860 rat GSE3068 sample 33
NES p.adjust
−2.67 1.2e−10
−2.81 1.0e−10
−2.46 1.1e−10
MCF7_Perturbations_from_GEO_up=insulin−like growth factor 1 GDS3484 mcf7:98
NES p.adjust
−2.4 1.2e−10
−2.73 1.0e−10
−2.64 1.1e−10
Running Enrichment Score
Ranked List Metric
Rank in Ordered Dataset

**Supplementary Figure 18. "T1D- and T2D-related" GSEA profiles in liver, adipose tissue and skeletal muscle from 5 month old *Mlx*KO mice relative to WT controls**

Normalized enrichment scores and q values are indicated in the upper right corner of each profile. Data used to generate the ridge plots shown in Figure 6D are re-graphed and included here along with additional representative profiles. See Supplementary File 4 for a complete list of all gene sets of significance included in this category.

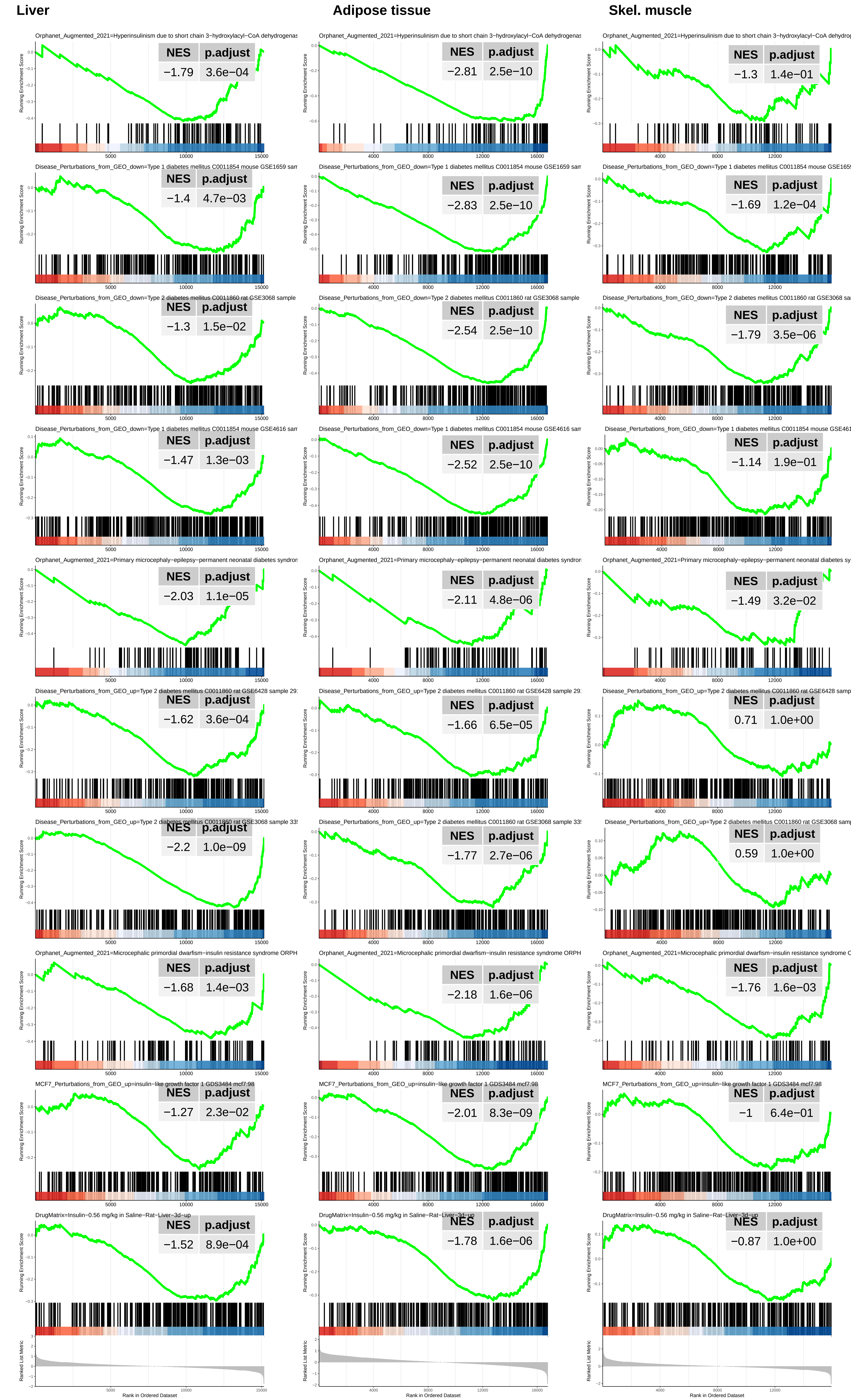
Liver
Adipose tissue
Skel. muscle
Orphanet_Augmented_2021=Hyperinsulinism due to short chain 3−hydroxylacyl−CoA dehydrogenas
NES p.adjust −1.79 3.6e−04
NES p.adjust −2.81 2.5e−10
NES p.adjust −1.3 1.4e−01
Disease_Perturbations_from_GEO_down=Type 1 diabetes mellitus C0011854 mouse GSE1659 sam
NES p.adjust −1.4 4.7e−03
NES p.adjust −2.83 2.5e−10
NES p.adjust −1.69 1.2e−04
Disease_Perturbations_from_GEO_down=Type 2 diabetes mellitus C0011860 rat GSE3068 sample
NES p.adjust −1.3 1.5e−02
NES p.adjust −2.54 2.5e−10
NES p.adjust −1.79 3.5e−06
Disease_Perturbations_from_GEO_down=Type 1 diabetes mellitus C0011854 mouse GSE4616 sam
NES p.adjust −1.47 1.3e−03
NES p.adjust −2.52 2.5e−10
NES p.adjust −1.14 1.9e−01
Orphanet_Augmented_2021=Primary microcephaly−epilepsy−permanent neonatal diabetes syndron
NES p.adjust −2.03 1.1e−05
NES p.adjust −2.11 4.8e−06
NES p.adjust −1.49 3.2e−02
Disease_Perturbations_from_GEO_up=Type 2 diabetes mellitus C0011860 rat GSE6428 sample 29
NES p.adjust −1.62 3.6e−04
NES p.adjust −1.66 6.5e−05
NES p.adjust 0.71 1.0e+00
Disease_Perturbations_from_GEO_up=Type 2 diabetes mellitus C0011860 rat GSE3068 sample 33
NES p.adjust −2.2 1.0e−09
NES p.adjust −1.77 2.7e−06
NES p.adjust 0.59 1.0e+00
Orphanet_Augmented_2021=Microcephalic primordial dwarfism−insulin resistance syndrome ORPH
NES p.adjust −1.68 1.4e−03
NES p.adjust −2.18 1.6e−06
NES p.adjust −1.76 1.6e−03
MCF7_Perturbations_from_GEO_up=insulin−like growth factor 1 GDS3484 mcf7:98
NES p.adjust −1.27 2.3e−02
NES p.adjust −2.01 8.3e−09
NES p.adjust −1 6.4e−01
DrugMatrix=Insulin−0.56 mg/kg in Saline−Rat−Liver−3d−up
NES p.adjust −1.52 8.9e−04
NES p.adjust −1.78 1.6e−06
NES p.adjust −0.87 1.0e+00
Running Enrichment Score
Ranked List Metric
Rank in Ordered Dataset

**Supplementary Figure 19. "T1D- and T2D-related" GSEA profiles in liver, adipose tissue and skeletal muscle from 5 month old *Myc*KO mice relative to WT controls *(23)*.**

Normalized enrichment scores and q values are indicated in the upper right corner of each profile. Data used to generate the ridge plots shown in Figure 6D are re-graphed and included here along with additional representative profiles. See Supplementary File 4 for a complete list of all gene sets of significance included in this category.

**Supplementary File 1. Complete list of all gene sets of significance from which the GSEA profiles depicted in Figure 6A and Supplementary Figures 3-9 are drawn.**

**Supplementary File 2. Complete list of gene sets identified as belonging to the categories indicated in Supplementary Figures10-15.**

**Supplementary File 3**. **Complete list of all "Cancer-related" gene sets of significance.** Within this category are those sets depicted in Figure 6 and Supplementary Figure 16-17.

**Supplementary File 4. Complete list of all T1D- and T2D-related gene sets of significance.** Within this category are those sets depicted in Figure 6D and Supplementary Figure 18-19.

**Supplementary File 5. Complete list of all gene sets used to generate the heat map shown in Figure 6E.**

**Supplementary File 6. Identities of the 3191 gene sets pertaining to the immune response, inflammation and cytokine production depicted in Figure 6F.**

**Supplementary File 7. Detailed information about the RNA-seq datasets used in Figure 6G-H.** These datasets were obtained from GEO databases and include data from normal mice of various strains, which were maintained on either standard or high-fat diets (HFDs).
